# The chromatin reader Dido3 is a regulator of the gene network that controls B cell differentiation

**DOI:** 10.1101/2021.02.23.432411

**Authors:** Fernando Gutiérrez del Burgo, María Ángeles García-López, Tirso Pons, Enrique Vázquez de Luis, Carlos Martínez-A, Ricardo Villares

**Affiliations:** Centro Nacional de Biotecnología/CSIC, Darwin 3, Cantoblanco, EL28049, Madrid, Spain; Centro Nacional de Investigaciones Cardiovasculares, Instituto de Salud Carlos III, Melchor Fernández Almagro 3, Madrid 28029, Spain

## Abstract

The development of hematopoietic cell lineages is a highly complex process that is governed by a delicate interplay of various transcription factors. The expression of these factors is influenced, in part, by epigenetic signatures that define each stage of cell differentiation. In particular, the formation of B lymphocytes depends on the sequential silencing of stemness genes and the balanced expression of interdependent transcription factors, along with DNA rearrangement. We have investigated the impact that the deficiency of DIDO3, a protein involved in chromatin status readout, has on B cell differentiation within the hematopoietic compartment of mice. Our findings revealed significant impairments in the successive stages of B cell development. The absence of DIDO3 resulted in remarkable alterations in the expression of essential transcription factors and differentiation markers, which are crucial for orchestrating the differentiation process. In addition, the somatic recombination process, which is responsible for generation of antigen receptor diversity, was also adversely affected. These observations highlight the vital role of epigenetic regulation, in particular the involvement of DIDO3, in ensuring proper B cell differentiation. This study show new mechanisms underlying disruptive alterations which deepen our understanding of hematopoiesis and may potentially lead to insights that aid in the development of therapeutic interventions for disorders involving aberrant B cell development.

## INTRODUCTION

B cells originate from hematopoietic stem cells (HSC) through a highly regulated process involving the activation or silencing of key transcription factors and concurrent DNA recombination events. HSC can differentiate into multipotent progenitors (MPP) that transition through various lymphoid progenitor cell stages, including lymphoid primed multi-potential progenitor (LMPP), common lymphoid progenitor (CLP), prepro-B, pro-B, pre-B, and immature B cells (Inlay et al., 2009).

During the CLP stage, somatic rearrangements occur between the diversity (DH) and joining (JH) gene segments of the heavy chain (HC) locus, marking the initiation of B cell lineage commitment (Allman et al., 2003). CLPs have the potential to differentiate into multiple cell types, such as B lymphocytes, T lymphocytes, NK cells, and dendritic cells (Manz et al., 2001; Kondo et al., 1997). In mice, the upregulation of B220 expression in a CLP subpopulation allows their differentiation into prepro-B cells. Prepro-B cells, characterized by the expression of tyrosine kinase AA4.1 (CD93), are committed to B cell differentiation (Rumfelt et al., 2006). The transition to the early pro-B state is marked by CD19 expression and the completion of DH-JH rearrangement (Otero et al. 2003). IL7R signaling contributes to the survival and transient proliferation of early pro-B cells, as well as the sequential reorganization of immunoglobulin (Ig) genes (Bertolino et al., 2005). The productive rearrangement of VH heavy chain variable segments with DJH (VH-DJH) in the late pro-B stage is crucial for the formation of a functional pre-B cell receptor (pre-BCR), which is essential for further survival and differentiation (Lutz et al., 2011).

Epigenetic mechanisms, including post-translational histone modifications, play a crucial role in controlling chromatin structure, DNA accessibility, and gene expression in a cell type-specific manner. Proteins that specifically recognize and interpret these modifications are known to be involved in development, differentiation, and tumorigenesis (Berdasco et al., 2010).

*Dido1* is a highly conserved gene found in various organisms, and its shortest encoded protein isoform, DIDO1, contains nuclear localization and export signals (NLS, NES), as well as a plant Zn-finger homeodomain (PHD). DIDO1 interacts with chromatin primarily through di/trimethylated lysine of histone H3 (H3K4me2/3) and functions as a chromatin reader (Tencer et al., 2017). Although the exact protein complexes associated with DIDO1 remain unknown, all three natural isoforms contain both the PHD and NLS domains. DIDO2, the least abundant isoform, includes a transcription elongation factor S-II central domain (TFIIS2M) required for the nucleolytic activity of RNA polymerase II, as well as a Spen paralog and ortholog C-terminal (SPOC) domain, a reader of RNA polymerase C-terminal domain (CTD) (Appel et al., 2023), with transcriptional corepressor activity. DIDO3, the main isoform, encompasses all the domains present in DIDO2, along with a coiled-coil (CC) region that likely has a structural role. Additionally, DIDO3 contains low complexity regions rich in proline, alanine, and arginine residues.

Attempts to delete *Dido1* have resulted in cell lethality. N-terminal truncation leads to the expression of a shorter protein driven by an internal promoter, which is associated with aneuploidy, centrosome amplification, and centromere-localized breaks (Guerrero et al., 2010). Mice with the *Dido1*ΔNT mutation exhibit myeloid neoplasms, and alterations in DIDO1 are linked to myelodysplastic syndrome in humans (Rojas et al., 2005). Deletion of the 3’ terminal exon (ΔE16) of *Dido1* results in a truncated form of DIDO3 and leads to embryonic lethality by gestation day 8.5 (Fütterer et al., 2012). Embryonic stem cells derived from these embryos lacking DIDO3 C-terminal fail to differentiate properly but retain their self-renewal capacity (Fütterer et al., 2021; 2017).

Given the embryonic lethality associated with *Dido1*ΔE16 deletion, we investigated the impact of DIDO3 deficiency on the hematopoietic lineage using mice carrying a floxed 3’-terminal exon 16 (flE16) of *Dido1* (Mora Gallardo et al., 2019) and a Cre recombinase gene under the control of the *Vav1* promoter. In these mice, the B cell population exhibited significant reductions, suggesting that DIDO3 plays a crucial role in chromatin remodeling necessary for B cell differentiation, thereby influencing developmental cell plasticity and lineage fate decisions.

## MATERIALS AND METHODS

### Mice

In brief, a mouse strain bearing an 3’-terminal exon 16 (E16) flanked by *loxP* sites was generated and interbred with Cre^VAV1^-producing mice heterozygous for the E16 deletion. Hematopoietic populations were isolated from the offspring. Mice were handled according to national and European Union guidelines, and experiments were approved by the Comité Ético de Experimentación Animal, Centro Nacional de Biotecnología, Consejo Superior de Investigaciones Científicas and by the Comunidad de Madrid (PROEX 055-14).

### Flow cytometry and cell sorting

Bone marrow cells from each sample were diluted in a solution of 0.5% PBS-BSA, 0.065% NaN_3_. To avoid nonspecific antibody binding, samples were blocked with Fc-Block (CD16/32; BD Pharmingen). Antibodies used were from eBioscience (B220-eF450, CD3-biotin, CD5-percpCy5.5, CD19-AlexaFluor700, CD93-APC, CD127 (IL7R)-PeCy7, Sca1APC), Beckman Coulter (B220-biotin, B220-PE, CD4-biotin, CD8-PE, CD8-biotin, cKit-APC, Gr1-biotin, Ly6C-biotin), BioLegend (CD4-PeCy7, CD11b-PeCy7), and BD Pharmingen (CD11b-biotin, CD21-APC, CD23-PE, CD43biotin, IgG1-PE, IgG3-PE, Ly6G-PE, Nk1.1-APC). Biotinylated antibodies were visualized with streptavidin-eF450 (eBioscience), streptavidin-PE (Southern Biotech) or streptavidin-PE-CF594 (eBioscience). Cell death was monitored using annexin V-APC (Immunostep) and DAPI. In *in vivo* assays, the cell cycle was analyzed by measuring BrdU incorporation into cell DNA 18 h after intraperitoneal injection of 200 mg BrdU (Invitrogen). Cells were labeled following the protocol for the BrdU Staining Kit for flow cytometry (Invitrogen). Analytical cytometric data was acquired using a Gallios cytometer (Beckman Coulter). Flow cytometry analyses were performed using FlowJo v10.6.0.

The same antibody combination used in flow cytometry was applied to the mouse bone marrow cells to select the target population in staining buffer (PBS containing 10% FBS), which was then sorted using a BD FACSAria Fusion cytometer. Data were analyzed with BD FACSDiva 8.0.1.

### *In vitro* differentiation of bone marrow cells

Bone marrow cells from control (WT) and *Dido1*ΔE16 mice (lin^−^: CD3, CD11b, CD19, CD45R (B220), Ly6G/C (Gr1), TER119), were purified with the EasySep Mouse Hematopoietic Progenitor Cell Isolation kit (STEMCELL Technologies) according to the manufacturer’s protocol. Isolated cells were grown in a semi-solid methyl cellulose matrix for the Mouse Hematopoietic CFU Assay in Methocult Media 3630 (STEMCELL Technologies), following the manufacturer’s instructions. The medium is supplemented with IL7, which promotes growth of B lymphocyte progenitor cells as pre-B colony-forming units (CFU-pre-B).

### RT-qPCR

RNA was extracted from sorted bone marrow LSK cells (Lin (B220, Ly6G/C (Gr1), TER119, CD3, CD11c,CD11b, F480)^-^, Sca-1^+^, cKit^+^, IL-R7^-^ with the RNeasy Plus Mini Kit (Qiagen) according to manufacturer’s instructions; quality was determined by electropherogram (Agilent 2100 Bionalyzer). To obtain cDNA, the SuperScriptIII FirstStrand Synthesis System for RT-PCR kit (Invitrogen) was used. Up to 1 µg total RNA was used as template in a final reaction of 20 µl. qPCR reactions were carried out 384-well plates in an ABI PRISM 7900HT system (Applied Biosystems) using SYBR Green qPCR Master Mix (Applied Biosystems). ABI SDS 2.0 software was used for data acquisition and analysis.

### RNA-seq

Pre-B (B220^+^, CD19^+^, IgM^-^) cells extracted from the bone marrow of WT and *Dido1*ΔE16 mice were isolated by sorting. RNA was purified using the RNeasy mini plus kit (Qiagen). Samples with about 100 ng of RNA with Integrity Number (RIN) ≥ 9 were processed for massive sequencing at BGI Genomics (Hong Kong) using the HiSeq 2000 platform.

### ATAC-seq

LSK cells of WT and *Dido1*ΔE16 mice were isolated by sorting and processed with the DNA Library Prep Kit (Illumina). The tagged DNA was purified with the DNA Clean ConcentratorTM5 Kit (Zymo Research). The samples were pre-amplified with the Nextera Index kit and the Nextera DNA Library Prep kit (both from Illumina), incorporating index sequences at the ends. DNA from the PCR reaction was purified using the DNA Clean ConcentratorTM5 Kit. To quantify tagged DNA, the 1x dsDNA HS Assay kit (Qubit) was used. Samples were then enriched in DNA sizes between 200 and 600 bp (corresponding to 1 to 3 nucleosomes) using AMPure XP magnetic beads (Beckman-Coulter), following the manufacturer’s guidelines. To sequence tagged DNA, a mixture was used of libraries with a balanced proportion of three samples each of WT and *Dido1*ΔE16 LSK cells, each with ∼100,000 pmol/l tagged DNA fragments. Massive sequencing was performed by the CNIC Genomics Service on a HiSeq 2500 platform using a Hi-Seq-Rapid PE Flowcell 2×50 kit.

### ChIP-seq

LSK cells of WT and *Dido1*ΔE16 from bone marrow mice were isolated by sorting. ChIP assays were performed with True MicroChIP-seq Kit (DIAGENODE Cat# C01010132) according to the manufacturer’s protocol. Briefly, samples were fixed with 1% formaldehyde for 10 minutes. Chromatin was sheared using Bioruptor® Pico sonication device (DIAGENODE Cat# B01060010) combined with the Bioruptor® Water cooler for 5 cycles using a 30’’ [ON] 30’’ [OFF] settings. The shearing efficiency of IP DNA was analyzed by PCR and fragments of 300-500 pb were visualized in 2% agarose gel.

Antibodies for histone marks (H3K4me3, True MicroChIP-seq Kit and H3K27me3, Abcam (ab195477)) were used. Equal amounts of 0.5 μg each of the rabbit IgG negative control and the H3K4me3 positive control antibody were added. For ChIP sequencing, libraries were prepared by the Genomics Unit of the Scientific Park Madrid and sequenced using NextSeq 2000 High Output Run Mode V4 (Illumina) as single-end 100-bp reads.

### Computational analysis

Raw fastq reads were aligned to the mouse reference genome (GRCm38/mm10 assembly, http://genome.ucsc.edu) using Bowtie2 (bowtie-bio.sourceforge.net/bowtie2) and Burrows-Wheeler aligner BWA-MEM 0.7.15 (github.com/lh3/bwa.git) with standard settings, and converted to BAM files using Picard tools 2.9.0 (http://broadinstitute.github.io/picard). Duplicate reads were removed using Picard tools. BED files were imported to RStudio and annotated using the R/Bioconductor package ChIPseeker (github.com/YuLab-SMU/ChIPseeker)(Yu, Wang, and He 2015). The promoter region was set to −1 kb to 200 bp of the TSS. We also used the Bioconductor packages org.Mm.eg.db and TxDb.Mmusculus.UCSC.mm10.knownGene for peak annotations. Gene ontology (GO), pathway annotation, and enrichment analyses were performed with GSEA (gsea-msigdb.org) using the Molecular Signatures Database (MSigDB) (go-basic.obo, downloaded Jan 15, 2020). Significance of overlap between data sets was calculated using the enrichPeakOverlap function implemented in ChIPseeker, setting the number of random permutations (nShuffle) of the genomic locations to 10,000. To assess read coverage distribution across the genome, bigWig files (10-bp genomic bins) were generated with bamCoverage/deepTools 2.3.1 (Ramírez et al. 2016) and normalized for differences. The aligned sequence reads, coverage, ChIP-seq and ATAC-seq peaks were visualized with IGV (https://igv.org; (Robinson et al. 2011). We used Metascape (metascape.org) for multi-omics analysis (Zhou et al. 2019).

#### RNA-seq data analysis

Sequenced reads were processed in parallel using the Galaxy RNA-seq pipeline (usegalaxy.org) and a combined approach with BWA-MEM, StringTie (ccb.jhu.edu/software/stringtie/) and edgeR (bioconductor.org/packages/edgeR). The combined approach was previously used to process RNA-seq data of *Dido1*ΔE16 in embryonic stem cells (GEO: GSE152346). Differentially expressed genes (DEG) were defined according to the absolute value of their log2 (fold change) ≥0.6 and FDR <0.05, unless otherwise stated. A Venn diagram was constructed to obtain co-expressed DEG among samples using the webtool at http://bioinformatics.psb.ugent.be/webtools/Venn/, provided by Bioinformatics & Evolutionary Genomics (Gent, BE). The biological significance of DEG was explored using STRING v.11.0 (string-db.org).

#### ATAC-seq data analysis

The model-based analysis of ChIP-seq (MACS; github.com/taoliu/MACS/) (Yong Zhang et al., 2008) version 2.1.2 and HOMER 4.11(Heinz et al., 2010) were used to calculate accessible regions of chromatin between WT and *Dido1*ΔE16 LSK cells. The overlap between accessible regions identified by MACS and HOMER was 80-90% (see Supplementary Table 4). To determine the equivalent accessible regions in the replicas, the Intersect function of BedTools (Quinlan 2014) and the BED files produced by MACS were used. Sequences which overlap the ENCODE black list regions (mm10-blacklist.bed.gz) were removed with BedTools before calculating accessible regions of chromatin. We accessed the significant change of chromatin accessibility between equivalent regions using DESeq2 and edgeR. It was defined as significantly changed if the peak showed fold change >2 and a false discovery rate (FDR) <0.05. Genes, promoters, UTRs, introns, and exons were allocated to the accessible regions with ChIPseeker.

#### ChIP-seq data analysis

BAM files were processed using MACS (github.com/taoliu/MACS/) (Yong Zhang et al., 2008) version 3.0.0b3 for enrichment scoring and peak calling. Briefly, peaks were then called using the callpeak function in MACS, with an extension size of 147. Differential binding between experimental conditions was performed using the bdgdiff function in MACS with a gap distance of 73bp and a minimum region length of 147bp. Read count normalization was performed on alignment files to account for sequencing depth differences, and then background corrected based on input using the bdgdiff function of MACS. Peaks overlap with ENCODE black list regions (mm10-blacklist.bed.gz) were removed from downstream analysis using Bedtools (Quinlan, 2014).

### Statistical analysis

Results are presented as mean ± standard error of the mean (SEM). Statistical significance was determined by 1-way analysis of variance (ANOVA) or Student *t* test using GraphPad Prism, version 6 software. P <0.05 was considered statistically significant.

### Data availability

ATAC-seq and RNA-seq data are deposited in the Gene Expression Omnibus database under accession code GSE157228, while ChIP-seq data is available under accession code GSE272156.

## RESULTS

### Loss of the DIDO3 protein results in anemia and severe peripheral lymphopenia

To examine *in vivo* the role of DIDO3 in hematopoiesis, we deleted E16 of *Dido1* in murine HSC early in development. Mice homozygous for the floxed allele (*Dido1*flE16/flE16) were crossed with mice carrying one functional and one mutated allele of *Dido1* (*Dido1*ΔE16, which lacks the C terminus of DIDO3) and the *Vav1:iCre* transgene. The resulting *Dido1*flE16/ΔE16;R26R-EYFP;iCre^Vav1^ mice (hereafter, *Dido1*ΔE16) were used as experimental subjects, and their *Dido1*flE16/+;R26R-EYFP;iCre^Vav1^ littermates (hereafter, WT) were used as controls, unless otherwise specified.

Ablation of *Dido1* E16 in early hematopoietic development was confirmed by qPCR on DNA samples from sorted bone marrow LSK cells, which showed a >99% reduction in *Dido1*ΔE16 DNA content (Suppl. Fig. 1A). Hematological analyses showed a significant decrease in cellular components when *Dido1*ΔE16 mice were compared to heterozygous littermate controls. Leukocyte numbers were reduced to 33% and erythrocyte numbers to 92% of their values. Within the leukocyte fraction, the lymphocyte subpopulation was responsible for the observed defect, with reduced cell numbers at 27% (Suppl. Fig. 1B). Peripheral blood samples from WT and *Dido1*ΔE16 mice were analyzed by flow cytometry for expression of the EYFP reporter; >97% of circulating cells were EYFP-positive in lymphoid and myeloid gates (Suppl. Fig. 1B, C).

As assessed by complete blood count and flow cytometry data, *Dido1*ΔE16 mice showed markedly decreased counts of lymphocyte subsets, most extreme in B220 (95% decrease), but also in CD4 (60%), CD8 (75%) and Nk1.1 (27%). The number of monocytes and granulocytes remained unchanged (Suppl. Fig. 1B, C). Furthermore, no differences were observed in the percentages of the different leukocyte populations between homozygous *Dido1* WT/WT mice and heterozygous WT/*Dido1*ΔE16 mice (data not shown).

### Loss of Dido 3 results in reduced total bone marrow cellularity

Since *Dido1* was previously linked most closely to stem cell developmental and differentiation functions, we analyzed the B-cell lineage in bone marrow. We used flow cytometry to identify LSK and common lymphoid progenitor lin-Sca1+cKit+IL7R+ (CLP) cell populations (Fig. 1A, B), and found no significant differences in the relative proportions of their respective populations. However, Dido1ΔE16 mice showed reduced total bone marrow cellularity and significantly lower absolute numbers of LSK cells (Fig. 1C).

**Figure 1.**
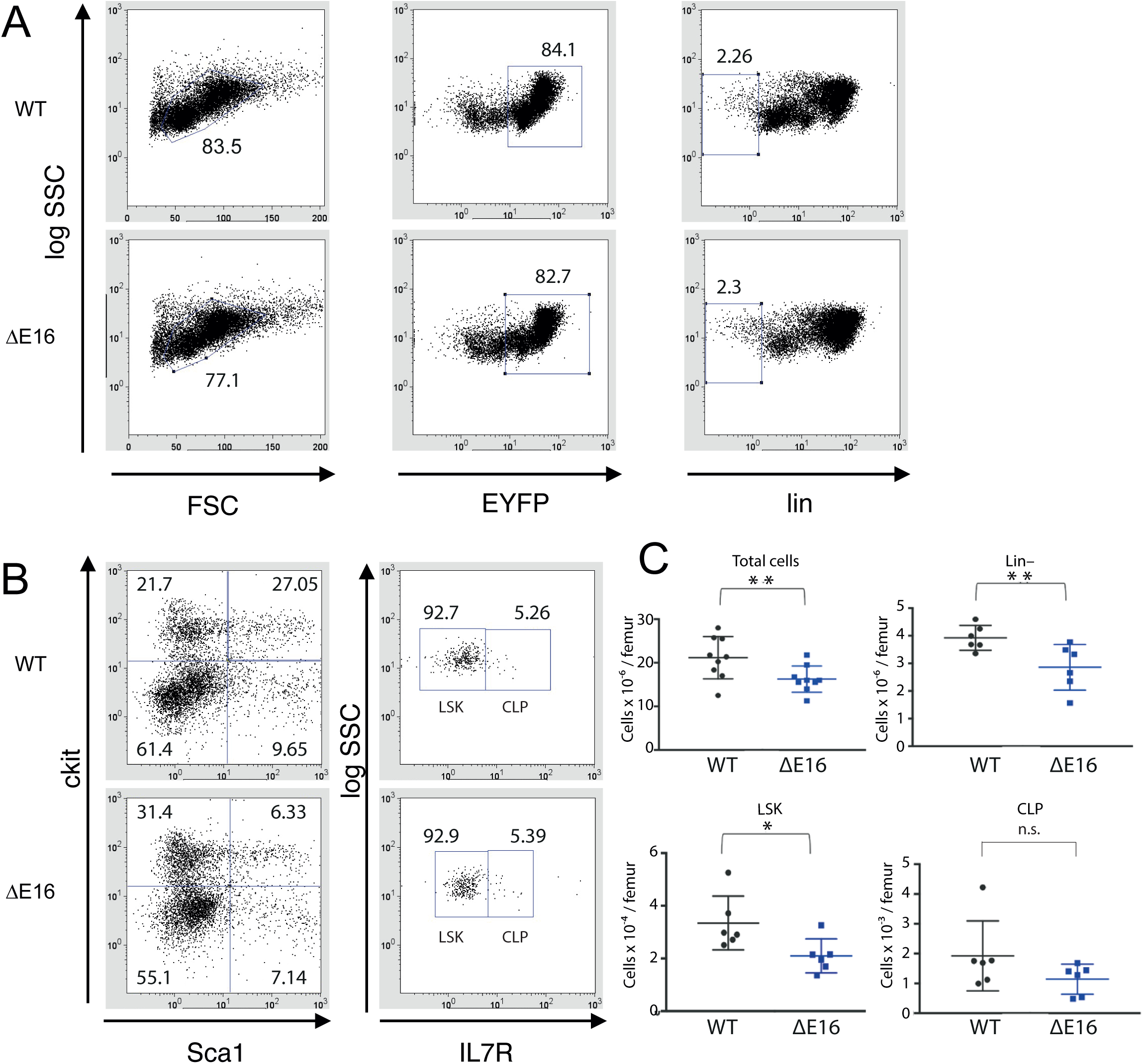
Characterization of hematopoietic progenitor cell populations. **A.** Forward/side scatter (FSC/SSC) plot of total cells in WT and *Dido1*ΔE16 mice and EYFP^+^ cells. Within EYFP^+^ cells, lin⁻ (Gr1⁻ Ter119⁻CD11b⁻CD3⁻B220⁻) were analyzed for Sca1, cKit, and IL7R expression to characterize LSK (Sca1^+^cKit^+^IL7R⁻) and CLP (Sca1^+^cKit^+^IL7R^+^) populations. **B.** Lin^-^ cells were analyzed for Sca1, c-kit, and IL7R expression to characterize LSK (Sca1^+^cKit^+^IL7R⁻) and CLP (Sca1^+^cKit^+^IL7R^+^) populations. **C.** Quantification of total (WT n = 9, *Dido1*ΔE16 n = 9), lin⁻, LSK and CLP cells (WT n = 6, *Dido1*ΔE16 n = 6) per single femur (t-test ** p<0.01, * p<0.05). Bars, mean ± SD.

We analyzed the distribution of the different subpopulations of B-cell precursors, namely prepro-B (EYFP^+^B220^+^CD19^-^CD93^+^IgM^-^), pro-B (EYFP^+^B220^+^CD19^+^CD43^+^IgM^-^), pre-B (EYFP^+^B220^+^CD19^+^CD43^-^IgM^-^) and immature B (EYFP^+^B220^+^CD19^+^IgM^low^) cells (Fig. 2A). The number of B220+ cells was substantially lower in *Dido1*ΔE16 mice than in controls (Fig. 2B). The most affected populations were pre-B and immature B cells, with prepro-B and pro-B cell numbers being similar in mice of both genotypes (Fig. 2C).

**Figure 2.**
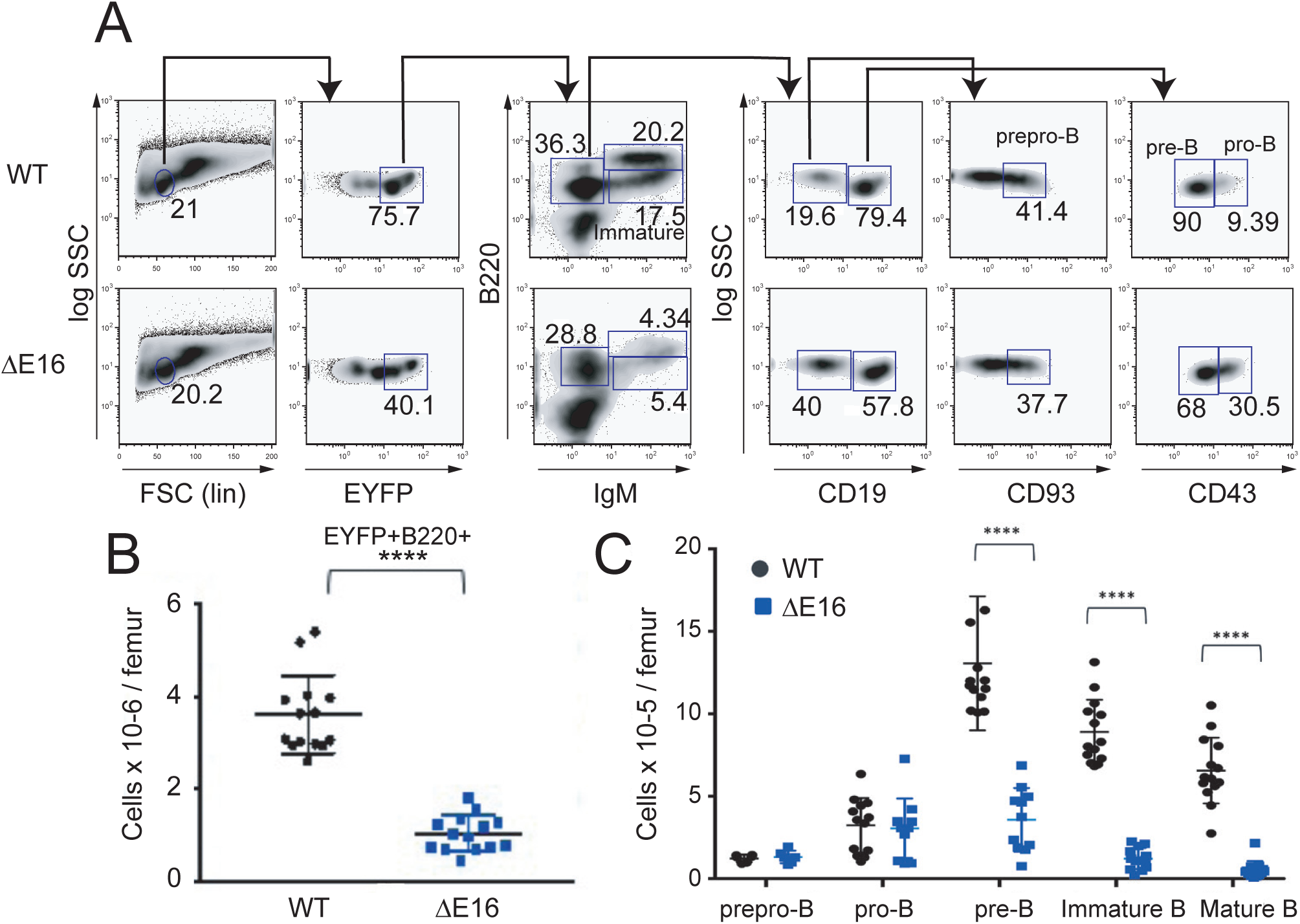
Characterization strategy by flow cytometry for lymphoid progenitor populations in mouse bone marrow. **A.** Forward/side scatter (FSC/SSC) plot of total cells in WT and *Dido1*ΔEI6 mice and EYFP^+^ cells. EYFP^+^ cells were gated by IgM and B220 markers to distinguish populations of mature EYFP^+^B220^+^IgM^high^ cells and immature EYFP^+^B220^+^IgM^low^ cells. Of the EYFP^+^B220^+^IgM⁻ subpopulation, prepro-B cells were identified as CD19⁻CD93^+^, pre-B as CD19^+^CD43⁻, and pro-B as CD19^+^CD43^+^. **B.** Numbers of EYFP^+^B220^+^ cells per femur in WT compared to *Dido1*ΔE16 mice (n = 14; ****, p<0.0001). **C.** Total bone marrow B precursors and mature recirculating B cells isolated from a single femur. Prepro-B (n = 6); pro-B, pre-B (WT n = 13, *Dido1*ΔE16 n = 11); immature and mature B cells (WT n = 14, *Dido1*ΔE16 n = 13). **B, C.** Mean ± SD are shown, t-test **** p<0.0001.

### DIDO3 Deficiency increases apoptosis and limits proliferation of immature B cells in bone marrow

Signaling from the pre-BCR complex promotes the survival and proliferation of cells that have successfully rearranged the *Igh* locus. This is a first checkpoint in which those rearrangements that give rise to a non-functional heavy chain lead to death by apoptosis. In a second checkpoint, immature B cells whose BCR is activated by an autoantigen in the bone marrow can undergo receptor editing to modify its light chain, or they can be deleted by apoptosis (Nemazee, 2017). To detect cell death in distinct precursor populations, we used flow cytometry on bone marrow samples from *Dido1*ΔE16 and WT mice, labeled with appropriate antibodies, and with annexin V and DAPI. Apoptosis levels were slightly higher, although non-significant, in *Dido1*ΔE16 mice. In the immature B cell population, both the number of apoptotic cells and the annexin V mean fluorescence intensity (MFI) in bone marrow of *Dido1*ΔE16 mice showed a significant increase relative to WT (Fig. 3A, B). This result suggests that the size of the immature B cell population in *Dido1*ΔE16 mice is limited by an increase in the number of cells that undergo apoptotic processes at this stage. Our data support the idea that DIDO3 is necessary to efficiently overcome the BCR signaling-dependent checkpoint.

**Figure 3.**
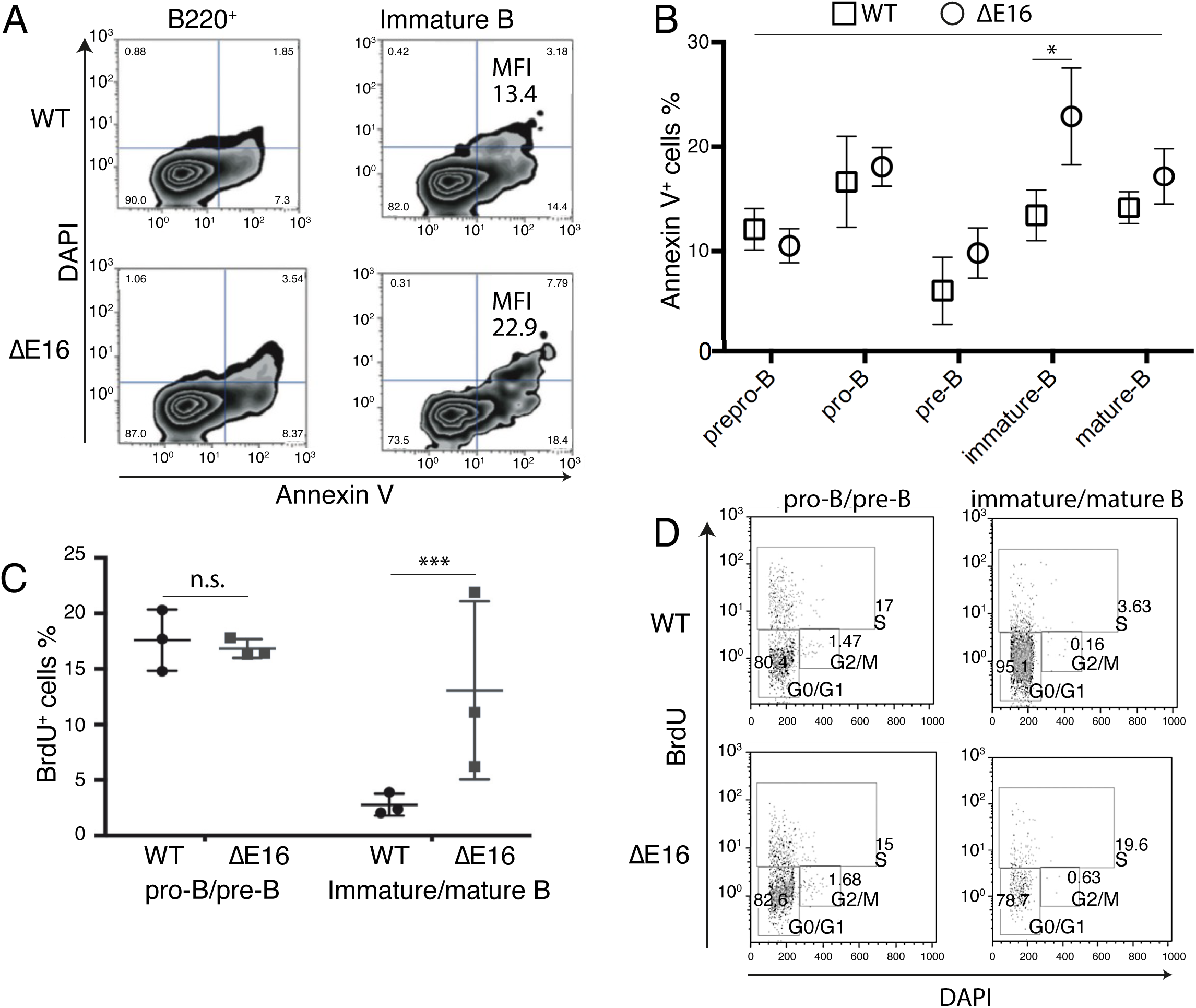
Effect of lack of DIDO3 on apoptosis and proliferation. **A**. Representative graphs of annexin V binding total B220+ (left) and immature B cells (right) from bone marrow of WT and *Dido1*ΔE16 mice. Mean fluorescence intensity (MFI) immature cells: 13.41 (WT, top) and 22.9 (*Dido1*ΔE16, bottom). Lower left, viable cells. Lower right, cells in early apoptosis (which bind annexin V and exclude DAPI). Upper right, cells in late apoptosis (which bind annexin V and incorporate DAPI). Upper left, necrotic cells. **B**. Mean percentages of annexin V+ cells in different precursor subpopulations (mean ± SD, n = 3, t-test * p<0.05). **C.** Percentages (mean ± SD; n = 3) of cells incorporating BrdU in pro-B/pre-B and immature/mature B cell populations, from bone marrow of WT and *Dido1*ΔE16 mice 18 h after intraperitoneal BrdU injection. **D.** Representative flow cytometry analysis of the cell cycle in pro-B/pre-B precursors (EYFP+B220+CD19+IgML) and immature/mature B cells (EYFP+B220+IgM+) from bone marrow of BrdU-treated WT and *Dido1*ΔE16 mice.

We used BrdU (5-bromo-2’-deoxyuridine) incorporation in DNA *in vivo* to identify and examine proliferating cells, and DAPI to determine DNA content and cell cycle stages. The percentage of cells that incorporated BrdU after 18 h post-intraperitoneal inoculation was quantified for pro-B/pre-B and immature/mature B cells. Cells from *Dido1*ΔE16 mice in pro-B/pre-B states showed BrdU incorporation similar to that of WT mice (Fig. 3C). In the immature cell population, there was a non-significant increase in the percentage and the MFI of cells that incorporated BrdU (Fig. 3D). The percentage of S phase cells of the pro-B/pre-B populations in both mice groups was similar, although an increase in the percentage of Dido1ΔE16 mature/immature B cells was observed in this phase (Fig. 3D). The increase in the percentage of immature B cells in the S phase in *Dido1*ΔE16 compared to control mice and the increase in cells that undergo apoptosis suggest that immature B cells from *Dido1*ΔE16 mice undergo an S phase block that eventually leads to apoptosis.

*In vitro* differentiation assays were carried out to determine whether the *Dido1*ΔE16 mouse B cell precursors showed alterations in their capacity to differentiate towards the B lineage. We purified cells lacking lineage-specific markers (lin^-^) by negative selection and plated them in a semi-solid matrix and medium optimized for B cell development, supplemented with IL7. In cultures from mice, between 25 and 35 colonies were counted in plates seeded with 2 x 10^6^ lin^−^ cells/ml; between 5 and 10 smaller cell groups/plate were also registered. No colonies were found in plates seeded with lin^-^ precursors from *Dido1*ΔE16 mice, although we detected some clusters consisting of a small number of disaggregated and undifferentiated cells (Suppl. Fig 2). Thus, B-lineage (B220^+^) cells from conditional mutant mice showed differentiation defects when stimulated in vitro, with blocked proliferation and maturation to B precursors under these conditions.

### Identification of essential regulators of stem cell function and the B cell pathway by ATAC-seq analysis of bone marrow LSK cells

As DIDO1 was demonstrated to be a chromatin interactor that recognizes accessible regions of chromatin with H3K4me3 histone marks (Tencer et al., 2017), we analyzed this population by Omni-ATAC-seq (Corces et al., 2017). Two biological replicas with accessible regions were used for differential analysis between WT and *Dido1*ΔE16 LSK cells.

Accessible regions were annotated with the ChIPseeker program (Yu et al., 2015) by associating the position of the accessible regions in each replica with the structure of the genes (promoter, 5’-UTR, 3’-UTR, exon, intron, or intergenic) in the mouse genome mm10. The largest percentage of accessible regions was concentrated in gene promoters and introns and in intergenic regions, in both WT and *Dido1*ΔE16 LSK cells (Fig. 4A). In the *Dido1*ΔE16 samples, however, there was a relative reduction in the number of introns and intergenic regions relative to promoters, which was related to the preferential association of DIDO1 to enhancer sequences over promoters (Engelen et al., 2015).

**Figure 4.**
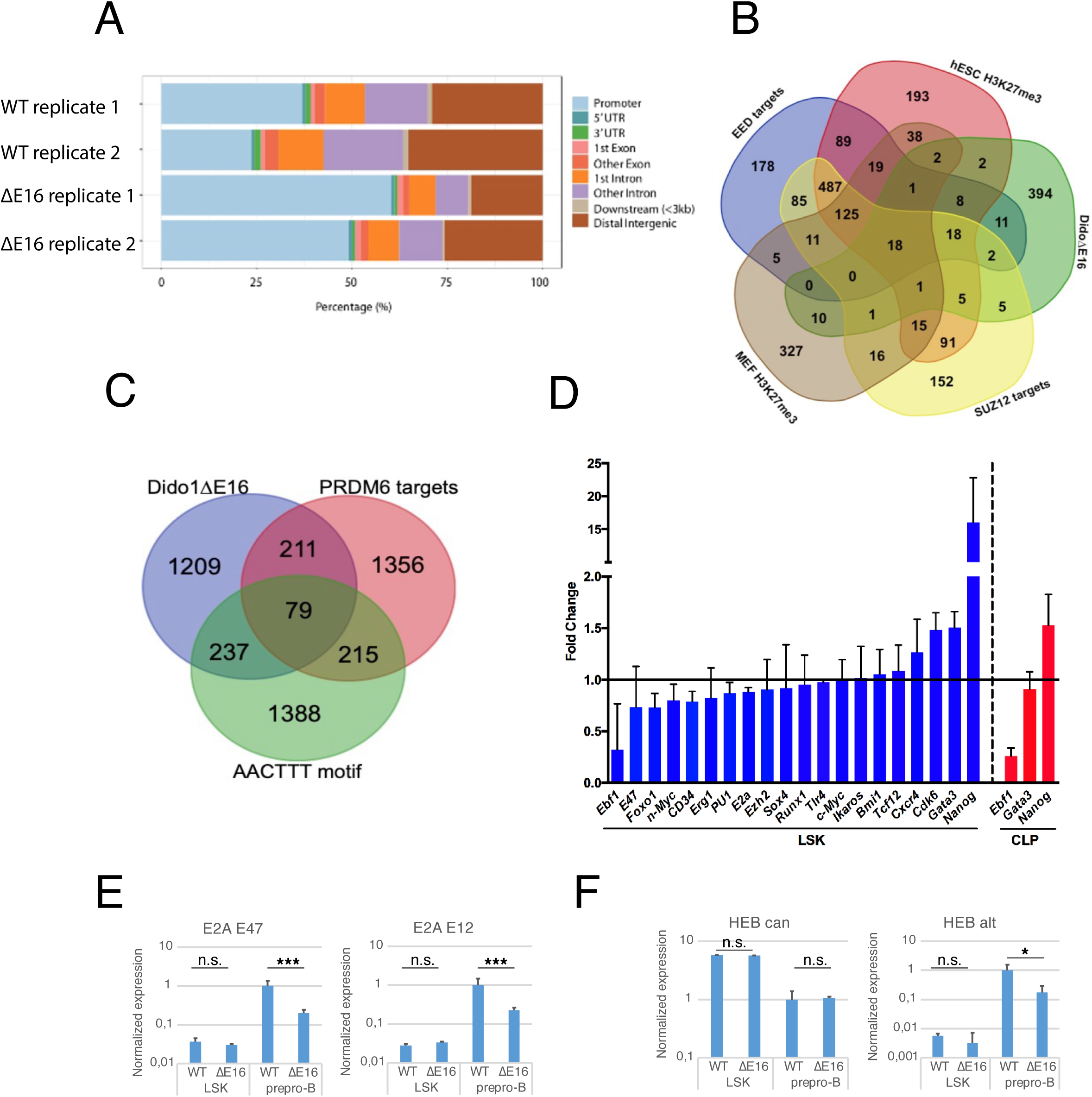
Effect of DIDO3 deficiency on chromatin accessibility and B cell developmental gene network. **A.** Distribution by distance to transcription start sites of chromatin open in WT and closed in *Dido1*ΔE16 LSK cells, as determined by ATAC-seq (mean values, n = 3). **B.** Venn diagrams show overlaps in promoter regions of differentially accessible chromatin with genesets associated with PRC2 activity. **C.** Overlap of differentially accessible chromatin in intronic and intergenic regions. **D.** Validation by real-time qPCR of the expression of potentially affected genes. **E.** Differential expression of *Tcf3* and its two isoforms in prepro-B cells and in lin⁻ precursors. Values are relative to WT prepro-B levels. **F**. Expression of various *Tcf12* isoforms in *Dido1*ΔE16 prepro-B cells, each relative to expression of the same region in WT cells.

A Gene Set Enrichment Analysis (GSEA) of the ATAC-seq data was performed. GSEA is a computerized method that determines whether *a priori* defined sets of genes show concordant and significant differences between two biological states (Subramanian et al. 2005). In seeking overlaps with “curated gene sets” (C2 collection) of MSigDB database, we found that the promoters with the least accessibility in *Dido1*ΔE16 LSK cells coincided with (1) genes with mixed epigenetic marks (H3K4me2 and H3K27me3) in mouse embryonic fibroblast (MEF) HCP genes (highCpG-density promoters), p=2.66 x 10^−57^; (2) silencing marks (H3K27me3) in embryonic cells, p=1.99 x 10^−38^; (3) human EED targets (a core subunit of PRC2), p=1.61x 10^−33^, and (4) human SUZ12 targets (also a core subunit of PRC2), p=5.56 x 10^−31^ (Supplementary Table 1). In accordance with these data, our previous microarray dataset (GEO: GSE85029, Fütterer et al., 2017) analyzed here by GSEA, indicate that genes silenced in DIDO3 deficient ESC correlate significantly with those silenced in mouse Suz12-deficient ESC; the genes overexpressed in these cells also coincided with those overexpressed in the Suz12–/– cells. There thus appears to be a relationship between epigenetic regulation of gene expression by PRC2 and the presence of DIDO3. We also constructed a Venn diagram that represents differentially-expressed genes (DEG) associated with distinct gene sets of MSigDB database (Fig. 4B). There was also correlation with (5) genes expressed after p53 activation (p<1.48 x 10^−24^), in agreement with the *Cdkn2a* alterations observed in *Dido1*ΔE16 mouse samples at later developmental stages (see below).

When analysis was limited to non-promoter regions (intra-or intergenic), using the MSigDB “regulatory target gene sets” C3 collection, there was significant overlap with (1) the targets of the putative histone-lysine N-methyltransferase PRDM6 (p<7.5 x 10^−24^), and (2) the set of genes with at least one occurrence of the highly conserved motif M17 AACTTT, a potential binding site for an unknown transcription factor in the region spanning up to 4 kb around their transcription start sites (p<5.91 x 10^−69^) (Fig. 4C). Within the 79 genes common to the three gene sets, we found a number of essential regulators for stem cell function and B cell pathway entrance, including *Akt3, Runx1, Ebf1, Igfr1, Tcf4, Cdk6*, and *Etv6*. GSEA analysis of the few identified sites that show higher accessibility in *Dido1*ΔE16 LSK cells yielded no relevant results.

To validate the expression data in additional samples by RT-qPCR, we selected several genes in the list of regions with different Tn5 accessibility in WT and *Dido1*ΔE16 LSK cells: genes related to the Polycomb group (PcG) proteins (*Bmi1, Ezh2*), maintenance and differentiation of stem cells (*Runx1, Nanog, Ikzf2* (Helios), *Tcf12* (Heb)), differentiation towards lineage B (*Klf3, Il7r, Ebf1, Tcf3* (E2A, E12/E47), *Spi1* (PU.1), *Foxo1*) or towards lineage T (*Gata3*), and cell proliferation (*Mycn* (N-myc), *Myc, Cdk6*). The analysis showed that expression of genes related to differentiation towards the B lineage was non-significantly lower in *Dido1*ΔE16 than in WT LSK cells. Other genes such as *Gata3* or *Cdk6* were expressed more significantly in *Dido1*ΔE16 LSK cells, whereas *Nanog* levels were 16 times higher than in WT LSK cells (Fig. 4D). Dido*1*ΔE16 CLP samples showed reduced *Ebf1* expression, whereas *Nanog* levels were normal. As in almost every developmental process, the three “E-proteins” play a fundamental role in hematopoiesis (Bain et al., 1998). These transcription factors are characterized by a basic helix-loop-helix (bHLH) domain with capacity to homo- and hetero-dimerize with themselves and other HLH proteins, and to bind to DNA through E-boxes (CACGTG sequences). *Tcf3* (which encodes E2A isoforms E12 and E47) and *Tcf12* (which encodes HEB isoforms HEBcan and HEBalt) have been implicated in B cell development (Anderson et al., 2007). To test the expression of the distinct isoforms, we designed specific primers for conventional RT-qPCR assays. *Tcf3* expression was lower in WT and *Dido1*ΔE16 LSK cells than in prepro-B cells; this reduction was due equally to E12 and E47 isoforms (Fig. 4E). However, while the expression of the HEBcan isoform of Tcf12 (assessed by exon 1/2 expression levels) was not affected by DIDO3 absence, HEBalt isoform expression (exon 1/2 of X9 mRNA isoform levels) dropped to 10% of the WT in Dido1ΔE16 prepro-B cells (Fig. 4F).

### DIDO3 deficiency alters gene expression in PRC2 targets and impairs immunoglobulin locus transcription in pre-B cells

As the first checkpoint in B cell development is dependent on pre-BCR activation and takes place at the pre-B stage(Von Boehmer and Melcherset al., 2010), where we found the greatest effect of the absence of DIDO3 (seen as lack of progenitor cell progression), we sequenced the transcriptome of WT and *Dido1*ΔE16 cells in this phase.

Using pooled samples (2-3 mice/pool), we purified RNA from flow cytometry-sorted pre-B cells. Two biological replicates were used to construct libraries for massive sequencing. We found 120 genes with higher expression in WT and 346 genes upregulated in Dido1ΔE16 pre-B cells (Fig. 5A and Supplementary Table 2).

**Figure 5.**
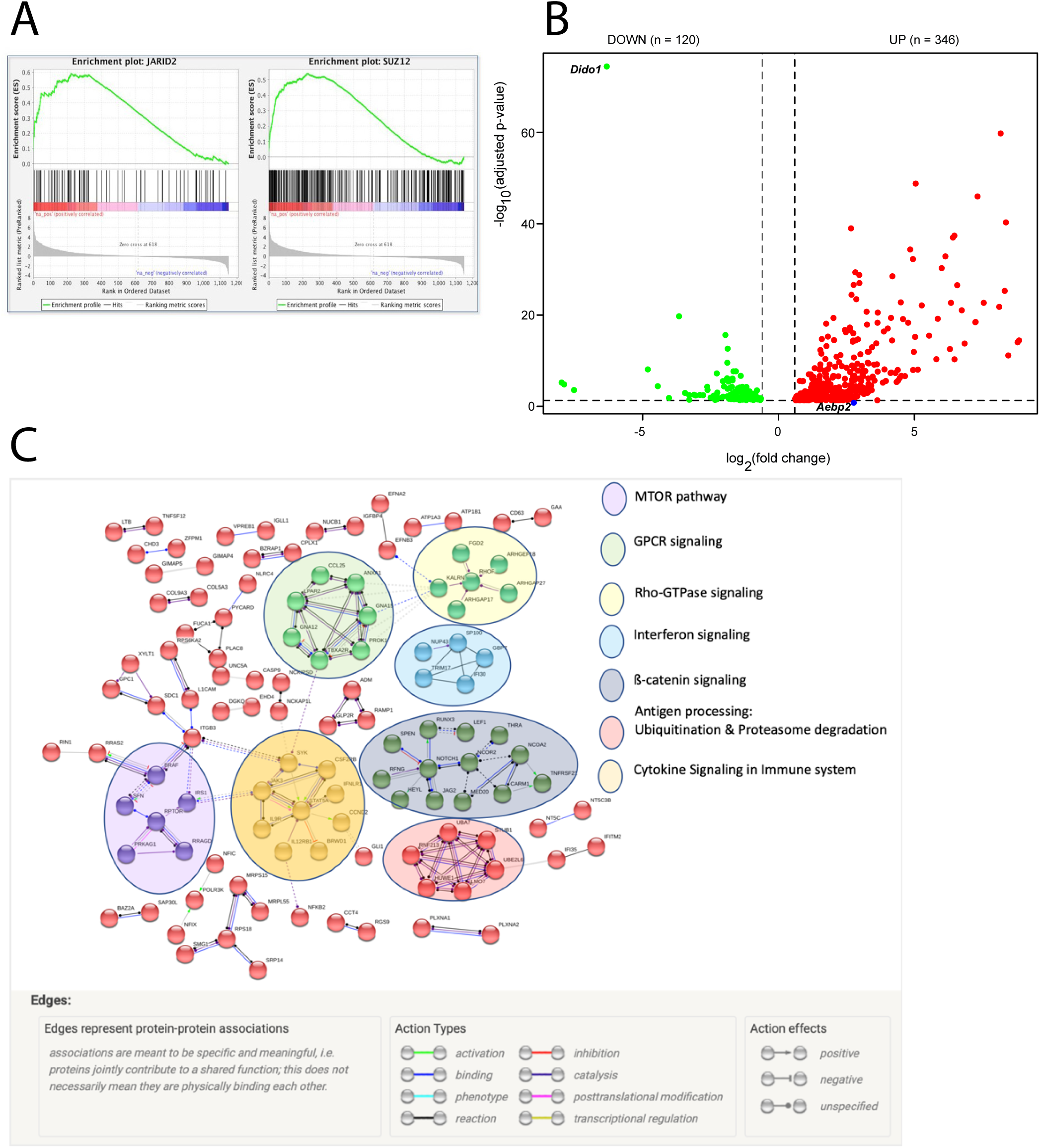
GSEA enrichment graphs of genesets relative to PRC2 activity and differentially expressed genes in pre-B cells. RNA-seq expression data ranked according to fold change in the transcription level in pre-B *Dido1*ΔE16 cells, relative to pre-B WT cells. Maximal correlation was observed between the expression of altered genes in pre-B *Dido1*ΔE16 cells and genes identified by ChIPseq as targets of PRC2 components JARID2 (p <0.001; FDR = 0.213) and SUZ12 (p <0.001: FDR = 0.217).

Protein-protein interactions in the STRING database v.11.0 were used to investigate the biological significance and functional association of differentially expressed genes. Gene Ontology (GO) terms with FDR <0.25 included positive regulation of leukocyte differentiation, cellular response to stimulus, positive regulation of lymphocyte activation, positive regulation of lymphocyte differentiation, positive regulation of T cell differentiation, positive regulation of B-cell differentiation, signal transduction, regulation of signaling, immune response, and regulation of leukocyte cell-cell adhesion. After gene clustering, the highlighted signaling pathways were MTOR, GPCR, Rho-GTPase, interferon, ß-catenin, cytokine signaling in the immune system, and antigen processing by ubiquitination and proteosome degradation (Fig. 5B).

For a second analysis we applied the GSEA tool, using a ranked list of genes with baseMean >1, ordered by fold change. By analysis against MSigDB M3 subcollection (GTRD, mouse transcription factor targets, https://www.gsea-msigdb.org/gsea/msigdb), we identified as downregulated gene sets of targets for PRC2 components JARID2 and SUZ12 (p<0.001) (Fig. 5C). *Aebp2*, a substoichometric PRC2 subunit that stimulates the K9 and K27 methylation of histone H3 (Grijzenhout et al., 2016), was upregulated (log_2_FC = 2.61 FDR = 0.067) in *Dido1*ΔE16 pre-B cells, although below the threshold of significance.

The relevance of pre-BCR in the pre-B developmental checkpoint and the PRC2 participation in V(D)J recombination (Su et al., 2003) led us to study the effect of DIDO3 deficiency on the transcription of *Igh* and *Igk* loci. Joining (J) and variable (V) regions of the *Igk* locus were not notably affected. There was over-representation of proximal *Igh* V regions (DQ52 and 7183 families), whereas intermediate and distal V fragments (mainly V_H_J558 and 3609 families) were under-represented in *Dido1*ΔE16 pre-B cells (Fig. 6A and Supplementary Table 3). We detected unexpectedly high transcription levels of DNA located between J and D fragments (Fig. 6B). In cells, this DNA segment is eliminated by recombination in the previous pro-B stage(Jung et al., 2006), or even in prepro-B, by RAG1/2 activity. Finally, we detected expression of the Igg2b (gamma-2b) gene, which suggested that isotype-switched IgG2b+ memory B cells co-purified with pre-B cells in numbers substantially higher than in WT samples (Fig. 6C).

**Figure 6.**
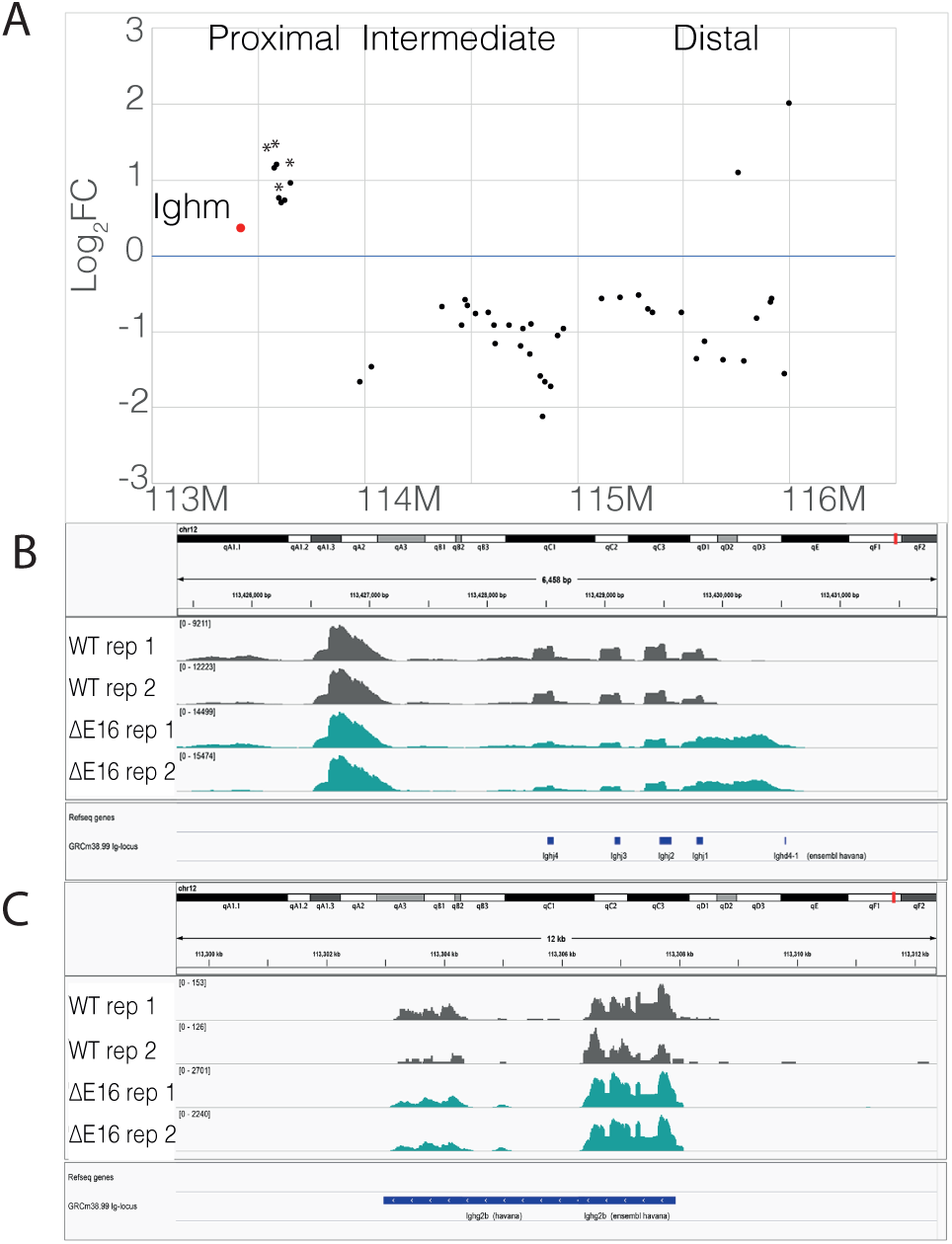
Analysis of the *Igh* locus in *Dido1*ΔE16 pre-B cells. **A.** Relative abundance of *Ighv* regions in the transcriptome of *Dido1*ΔE16 pre-B cells, shown as logFC of WT levels (*, p<0.05). Bottom, chromosome coordinates; top, the main regions (proximal, intermediate, distal) in which distinct V families map. **B**. Integrative Genomics Viewer (IGV; http://www.broadinstitute.org) plot with RNA-seq data showing transcription from the DNA region between D and J regions; J fragments and the more distal D fragment (D4) positions are represented. **C.** IGV plot with RNA-seq data showing transcription of the *Igg2b* gene in *Dido1*ΔE16 pre-B samples.

### Chromatin immunoprecipitation sequencing (ChIP-seq) analysis reveals the role of Dido 3 genes with H3K27me3

Trying to identify the effects of DIDO3 on PRC2 regulation, we performed H3K27me3 chromatin immunoprecipitation sequencing analysis of LSK cells isolated from WT and mutant *Dido1*ΔE16 mice (Figure 7, and Supplementary Tables 4 and 5). The largest percentage of H3K27me3 peaks was concentrated in gene introns and in intergenic regions in both WT and Dido1ΔE16 LSK cells (Supplementary Table 4). In WT, however, we observed slightly more genes with H3K27me3 enrichment relative to *Dido1*ΔE16 LSK cells. We found 58 regions (48 genes) with significantly more H3K27me3 enrichment in WT compared with *Dido1*ΔE16 and 44 regions (33 genes) that had more H3K27me3 enrichment in *Dido1*ΔE16 compared with WT. We also found 726 regions (436 genes) had no significant H3K27me3 enrichment in LSK cells (Figure 7, Supplementary Table 5, and Supplementary File 1). Genes highlighted in Supplementary Table 5 are involved in RNAPII transcription, RNA processing, histone binding and modification, cell identity and differentiation, immune response in B-cells, and maintenance of hematopoietic stem cells/hematopoiesis, among other functions. Some of these genes were differentially expressed in pre-B cells (see below).

**Figure 7.**
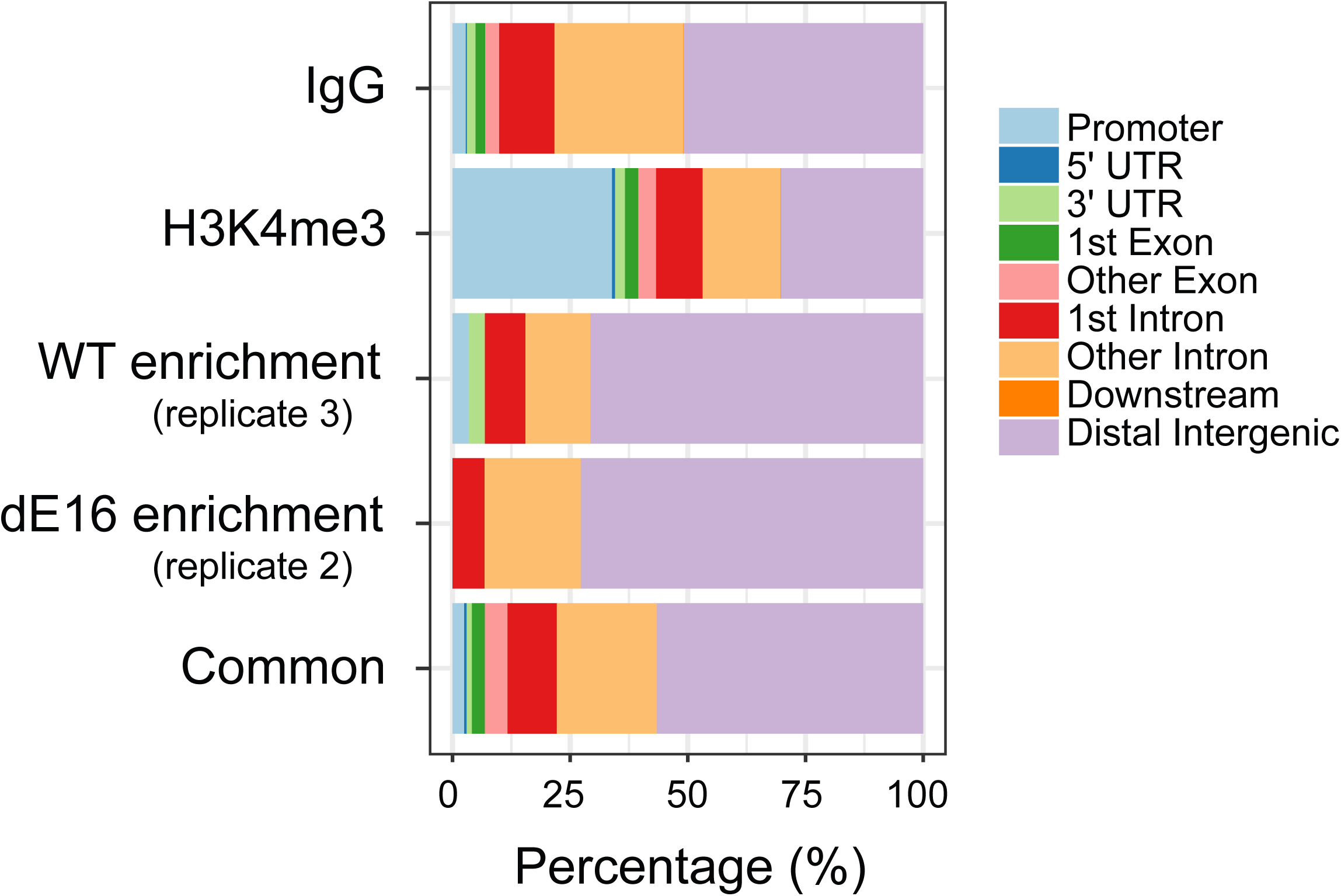
H3K27me3-enriched genomic regions in WT and *Dido1*ΔE16 LSK cells. Distribution of the enriched-peak distances to the nearest transcription start site as determined by ChIPseeker (Yu et al., 2015). The distribution of control IgG and H3K4me3 peaks in WT LSK cells is also shown.

We also compared the WT and *Dido1*ΔE16 H3K27me3 ChIP-seq data with other GEO datasets reported by two different studies. Specifically, we use Ezh2-KO (GSM2091489, GSM2091491) and Cebpa-KO LSK cells (GSM1054811, GSM1054814) datasets (Supplementary Table 6). The analysis indicate a preference for H3K27me3 binding in gene introns and intergenic regions, as well as no differences between wild-type and mutant LSK cells (Suppl. Fig. 3 and Supplementary Table 7). On the other hand, less than 10-12% of the H3K27me3 peaks in the mouse genome in WT and Dido1ΔE16 LSK cells overlap with Ezh2-KO and Cebpa-KO datasets (Supplementary Table 8). The result would be influenced by the different experimental conditions used (e.g., histone H3K27me3 antibody) in the three studies.

The combined analysis of differentially expressed genes (RNA-seq), chromatin accessible regions (ATAC-seq), and H3K27me3 ChIP-seq data reveals that only a few genes were detected simultaneously by different experiments, but a large number share a similar function or belong to a similar pathway (Figure 8). It is common in meta-analysis that we observe little direct overlap among studies due to the variations in the biological assays used (Figure 8A). However, we see more functional overlap (Figure 8B), as these studies probably pick up different subsets of gene members of the same biological processes. Figure 8C shows “immune effector process” (GO:0002252; logP = −11.5) as the most enriched GO term in downregulated genes in RNA-seq.

**Figure 8.**
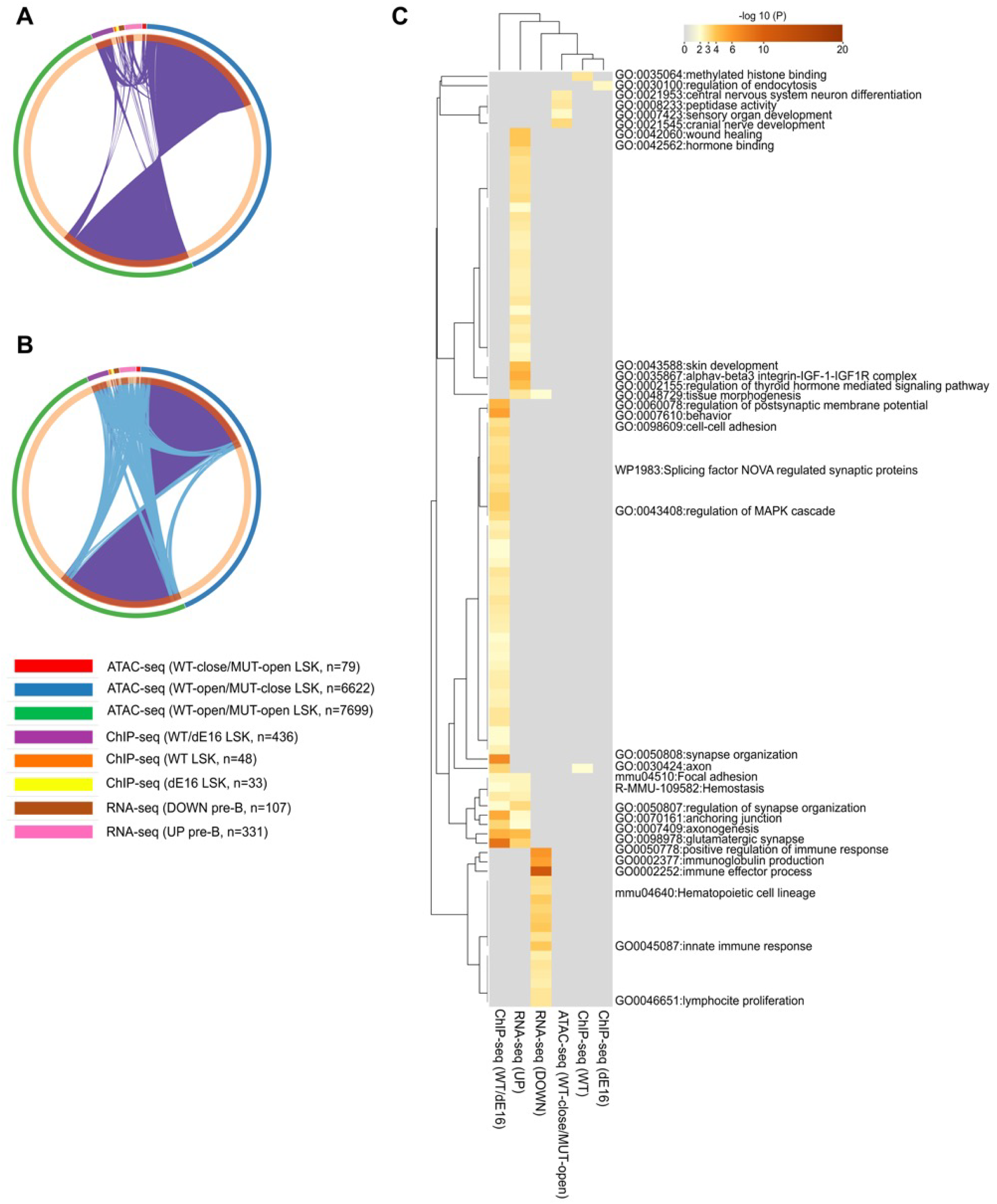
Metascape analysis of ATAC-seq, ChIP-seq, and RNA-seq experiments. **A.** overlaps between gene lists. **B.** overlaps between genes sharing the same enriched ontology terms (i.e., GO Biological Processes, Molecular Functions or Cellular Components, KEGG Pathway, Reactome Gene Sets, WikiPathways). On the outside, each arc represents the identity of each gene list, using the same color code as indicated in the legend. On the inside, each arc represents a gene list, where each gene member of that list is assigned a spot on the arc. Dark orange color represents the genes that are shared by multiple lists and light orange color represents genes that are unique to that gene list. Purple lines link the same gene that are shared by multiple gene lists (notice a gene that appears in two gene lists will be mapped once onto each gene list, therefore, the two positions are purple linked). Blue lines link the genes, although different, fall under the same ontology term (the term has to statistically significantly enriched and with size no larger than 100). The greater the number of purple links and the longer the dark orange arcs implies greater overlap among the input gene lists. Blue links indicate the amount of functional overlap among the input gene lists. **C.** hierarchical cluster of enriched ontology terms. The heatmap cells are colored by their p-values, gray cells indicate the lack of enrichment for that term in the corresponding gene list.

In WT and *Dido1*ΔE16 LSK cells, around 26-28% of the H3K27me3 binding sites overlap with chromatin-accessible areas in *Dido1*ΔE16 LSK cells (Supplementary Table 9). The differential chromatin-accessible areas “(close)/MUT(open)” in ATAC-seq coincide with only 1% of the H3K27me3 marks (Supplementary Table 9). In contrast, as shown in Supplementary Table 10, 37% of H3K4me3 binding sites overlap with chromatin-accessible regions in WT versus 19% in *Dido1*ΔE16 LSK cells, as expected.

Large areas of repressive H3K27 methylation and smaller areas of activating H3K4 methylation make up bivalent chromatin domains. In embryonic stem cells (ESC), these bivalent areas maintain developmental genes inactive while maintaining their readiness for activation. PRC2 is a crucial component in the mechanism of bivalent gene silencing and activation through the regulation of chromatin accessibility (Mas et al. 2018). Supplementary Table 11 lists bivalent genes upregulated or downregulated in RNA-seq.

## DISCUSSION

Although DIDO3, which is expressed ubiquitously in the organism, presumably has a more general function in determining chromatin structure, its dramatic influence on the B cell population led us to focus our study on this cell lineage. Speculations on the molecular mechanism involved have been treated elsewhere (Fütterer et al., 2024; García-López et al., 2024; Pacios-Bras et al., 2019; Gatchalian et al. 2013; Mora Gallardo et al., 2019; Sanchez de Diego et al., 2014; Gatchalian et al., 2013). Here we demonstrate that DIDO3 plays a role in HSC progression into B cell differentiation.

In lymphopoiesis B, EBF1 is the key transcription factor for B cell specification and compromise (Treiber et al., 2010). Ectopic expression of *Ebf1* in HSC directs differentiation towards the B lineage at the expense of other lineages (Zhang et al., 2003). *Ebf1^−/–^* mice maintain a developmental blockade of the B lineage in the prepro-B stage and lack mature B cells(Lin and Grosschedlet al., 1995). Our mouse model shows partial arrest in B cell development in the prepro-B and pro-B populations, comparable to the phenotype of *Ebf1*^+/–^ mice. Our RT qPCR assays show that *Ebf1* expression in purified *Dido1*ΔE16 CLP is approximately one-third of that observed in WT CLP cells, where the specification process towards the lymphoid lineage begins. Heterozygotic deletion of *Ebf1* resulted in an impaired response to IL7 *in vitro*(Åhsberg et al., 2013), which provides a possible explanation for the observed stage-specific reduction in cellular expansion. The loss of DIDO3 in the LSK state activates *Nanog* transcription, which negatively modulates *Ebf1*. Decreasing EBF1 levels would reduce the potential of *Dido1*ΔE16 CLP cells for differentiation into the B lineage.

All the canonical E proteins are important for lymphoid development (Kee, 2009), HEB (*Tcf12*) and E2-2 (*Tcf4*) are thought to contribute to the total E protein dose needed for B cell development, since the absence of either factor led to a decrease in pro-B cell numbers, but no obvious developmental block (Zhuang, Cheng, and Weintraub et al., 1996). Of the E proteins, E2A is necessary to maintain *Ebf1*, *Foxo1*, and *Pax5* expression, and hence the B cell genetic program. B cell development in Tcf3^+/–^;Tcf12^−/–^ mice is completely blocked at the CLP stage(Welinder et al., 2011). It is not known, however, whether this arrest is due to the lack of HEBcan, HEBalt, or both isoforms. In our case, neither is completely absent, but whereas HEBcan levels were normal, HEBalt levels dropped in *Dido1*ΔE16 prepro-B cells to 10% of that in, which suggests an undescribed function for this isoform. Lack of E2A function results in an inability to rearrange *Igh* gene segments, which can be rescued with a single dose of E2A (Bain et al., 1994). Although it remains to be determined whether this effect is dose-dependent, it could explain our results with *Dido1*ΔE16 pre-B cells, in which a substantial number of RNA reads correspond to the segment between D and J fragments, which was deleted efficiently in WT samples.

We found significant association of the transcriptional alterations in *Dido1*ΔE16 cells with a number of PRC2 targets and genes bearing H3K27me3 epigenetic marks, which suggests that DIDO3 is involved in determining specific PRC2 activity in the B cell lineage. Moreover, loss of EZH2 is known to cause impaired distal VH–DJH rearrangement and *Igh* locus contraction (Su et al., 2003). There is thus a preference to use proximal V fragments as in our model, which reinforces the view that DIDO3 acts in coordination with PRC2.

We have previously shown differential expression of the DIDO isoforms in MPD (Agnes Fütterer et al., 2005) and others reported that BCR-ABL1 translocation (Philadelphia chromosome; Ph) in myelocytic leukemia (CML) might be involved in modulating *DIDO1* expression in a JAK2V617F-independent manner (Berzoti-Coelho et al., 2018; 2016). Disorders in multiple signaling pathways and genetic abnormalities combined with the Ph are essential for the evolution of different types of leukemia; however, why MPD evolve specifically into CML, acute myeloid leukemia, acute leukemia leukocytosis, or mixed-phenotype acute leukemia is currently unclear. BCRABL1 kinase hyperactivity leads to the activation of signaling pathways and deregulation of cellular processes(Kumar et al., 2013; Sattler and Griffinet al., 2003), and hyperactivation of most of these pathways have been confirmed in CML and ALL mouse models. BCRABL1 activity channels mainly through JAK/STAT and PI3K/AKT/mTOR pathways, and through CCAAT/enhancer-binding proteins (C/EBP), which regulate normal myelopoiesis as well as myeloid disorders (Kang et al., 2016). Our Gene Ontology analysis of RNA-seq data showed that, in cells lacking DIDO3, the expression of PI3K/mTOR and JAK/STAT pathway components, as well as *Cebpb*, are affected at the transcriptional level. Statistical analysis of DNA methylation showed that target genes of NANOG (Tommasi et al., 2014), Polycomb-group target genes, and/or genes with an AACTTT promoter motif can have a methylation profile that seems to indicate predisposition to development of cancer, such as pancreatic ductal adenocarcinoma (Guler et al., 2020) and has previously been associated with the promoter of MEF2C and enriched at the promoters of genes involved in neurodevelopment (Mohajeri et al., 2022). Interestingly, this could be correlated with the anomalous brain development that we previously found in *Dido1* mutants (Villares et al., 2015).

In all, our data provide evidence of the involvement of DIDO3, a reader of histone post-translational modifications (PTM), in several processes of bone marrow hematopoiesis, including cell cycle progression, apoptosis, and differentiation. DIDO3 deficiency is associated with alterations in DNA accessibility and transcriptional defects of major B cell differentiation master regulators. This association seems to be related, at least in part, to the relationship of DIDO3 with PRC2. Remaining open questions include the DNA methylation and the general histone PTM landscape in DIDO3-deficient cells, as well as the consequences of DIDO3 deficiency in peripheral B cell differentiation and hematopoietic cancer development.

## Supporting information

Suppl3mentary Table 3

## ACKNOWLEDGEMENTS

We thank the Genomics Unit and the Service of Cardiovascular Physiology of the Animal Facility of the Spanish National Center for Cardiovascular Research (CNIC) for ATAC-seq and complete blood count processing, respectively. We also thank Catherine Mark for editorial assistance. This work was supported by grants from the Spanish Ministerio de Ciencia e Innovación (PID2019-110574RB-I00, SEV-2017-0712 and SAF2016-75456-R) and the Fundación Ramón Areces (XVIII Concurso Nacional para la Adjudicación de Ayudas a la Investigación en Ciencias de la Vida y de la Materia) and the Comunidad de Madrid (S2022/BMD-7321/MITIC-CM).

## AUTHOR CONTRIBUTIONS

FGB designed, performed, and analyzed most experiments and co-wrote the paper, MAG-L performed experimental work, TP performed computational data analysis and drafted the manuscript, CMA conceptualized experiments, analyzed data, co-wrote the paper and secured funding, RV designed the research, performed the experiments, and co-wrote the paper. All authors contributed to the analysis and discussion of the data.

## CORRESPONDING AUTHOR

Correspondence to Ricardo Villares.

