## Supplementary Figure 1 for "The chromatin reader Dido3 is a regulator of the gene network that controls B cell differentiation"

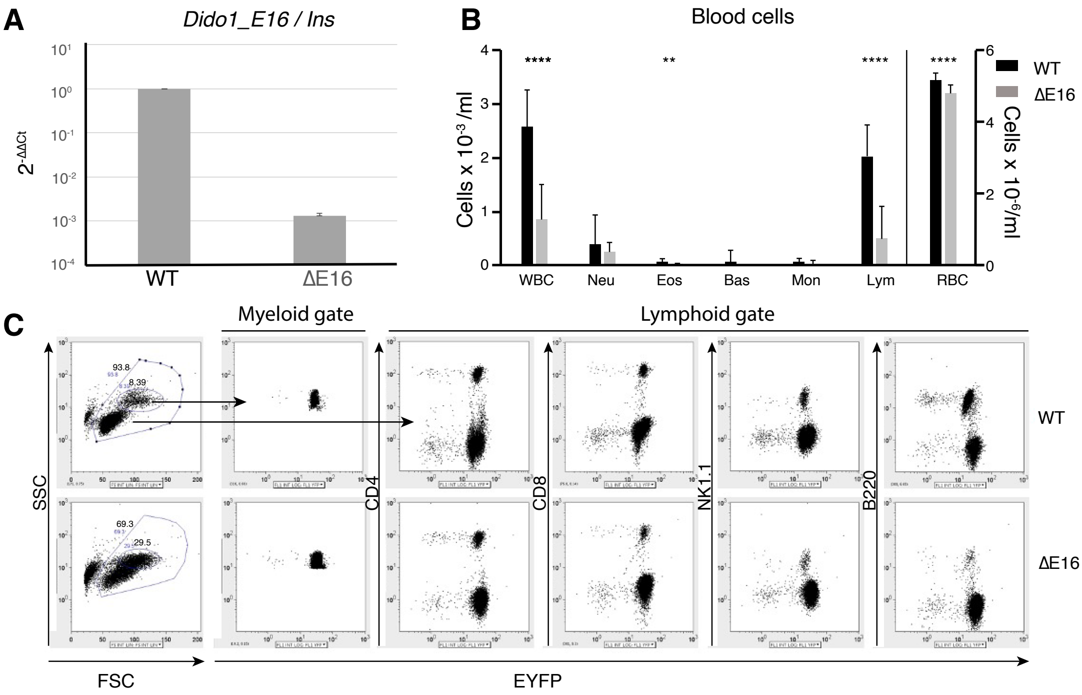


**Supplementary figure 1.** Deletion of *Dido1* exon 16 in hematopoietic lineage. **A.** Real-time qPCR of genomic DNA from bone marrow lin⁻ cells, normalized by ΔΔ Ct to the insulin gene (internal control) and WT exon 16 (internal reference). **B.** Complete blood count in WT (n = 18) and Dido1ΔE16 (n = 8) mice. Each column shows the mean number of total leukocytes (WBC), neutrophils (Neu), eosinophils (Eos), basophils (Bas), monocytes (Mon), lymphocytes (Lym) and eritrocytes/reticulocytes (RBC); bars indicate the standard error of the mean (SEM), t-test **** p <0.0001, ** p <0.01. **C.** Flow cytometry analysis of peripheral blood WBC from WT (top row) and Dido1ΔE16 (bottom row) mice, showing cells that express EYFP as reporter of recombinase activity and *Dido1* exon 16 deletion. Left to right: Forward/Side Scatter (FSC/SSC) profile and the EYFP reporter expression in the myeloid gate and in CD4+, CD8+, NK1.1+ and B220+ cells.
