## Supplementary Figure 2 for "The chromatin reader Dido3 is a regulator of the gene network that controls B cell differentiation"

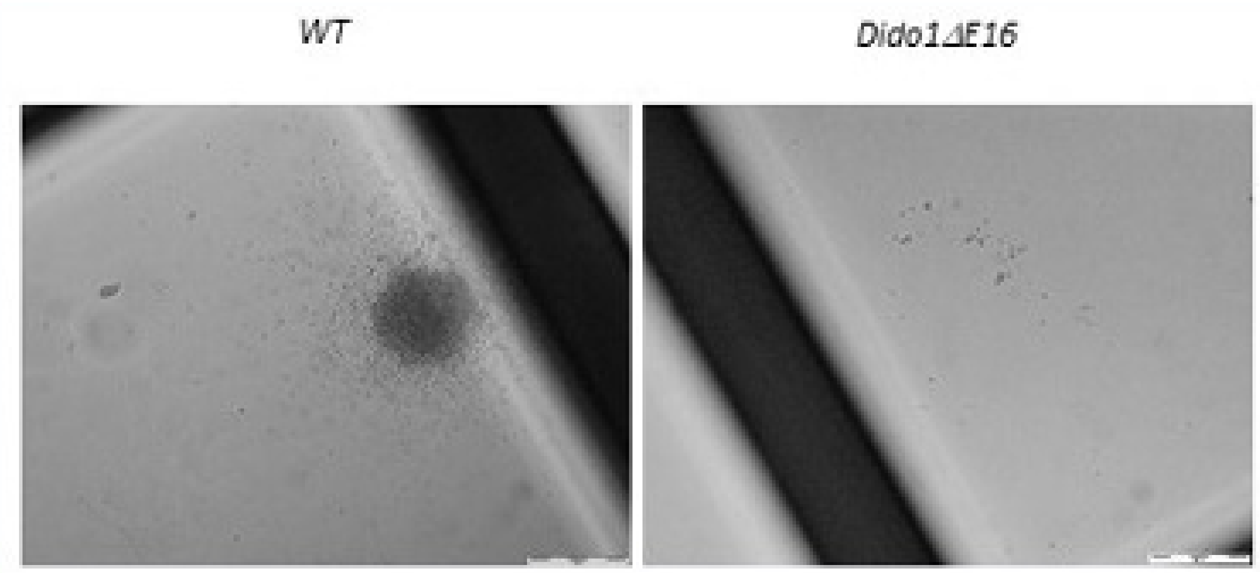


**Supplementary figure 2.** Representative micrographs of colonies obtained from lin cells from wt and Dido1ΔE16 mice. $Lin^-$ cells were plated in a semi-solid matrix and photographed after 10 days in culture. Progenitor cells able to generate pre-B colony-forming units gave rise to colonies in IL7-supplemented cultures.
