## Supplementary Figure 3 for "The chromatin reader Dido3 is a regulator of the gene network that controls B cell differentiation"

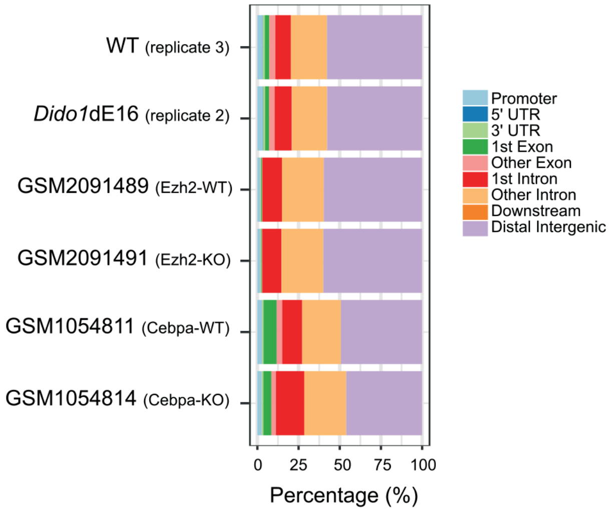


**Supplementary Figure 3.** H3K27me3-enriched genomic regions in LSK cells. Distribution of the enriched-peak distances to the nearest transcription start site as determined by ChIPseeker (Yu et al., 2015). The genomic coordinates of H3K27me3 peaks for Ezh2-WT, Ezh2-KO, Cebpa-WT, and Cebpa-KO LSK cells were downloaded from the ChIP-Atlas database (https://chip-atlas.org, (Zou et al., 2024)).
