## Supplementary Table 1 for "The chromatin reader Dido3 is a regulator of the gene network that controls B cell differentiation"

GSEA analysis of ATAC-seq data

Overlap Results

| | |
|---|---|
| Collection(s): | C2 |
| # overlaps shown: | 50 |
| # genesets in collections: | 4762 |
| # genes in comparison (n): | 1098 |
| # genes in universe (N): | 45956 |

| Gene Set Name | # Genes in Gene Set (K) | Description | # Genes in Overlap (k) | k/K | p-value | FDR q-value |
|---|---|---|---|---|---|---|
| MEISSNER_BRAIN_HCP_WITH_H3K4ME3_AND_H3K27ME3 | 1069 | H3 dimethylation at K4 (H3K4me2) and trimethylation at K27 (H3K27me3) in brain. | 135 | 0,1263 | 2,66E-57 | 1,27E-53 |
| BENPORATH_ES_WITH_H3K27ME3 | 1118 | (H3K27me3) mark in their promoters in human embryonic stem cells, as identified by ChIP on chip. | 113 | 0,1011 | 1,99E-38 | 4,74E-35 |
| CHEN_METABOLIC_SYNDROM_NETWORK | 1210 | (MEMN) claimed to have a causal relationship with the metabolic syndrom traits. | 111 | 0,0917 | 6,59E-34 | 1,05E-30 |
| BENPORATH_EED_TARGETS | 1062 | the Polycomb protein EED [GeneID=8726] in human embryonic stem cells. | 103 | 0,097 | 1,61E-33 | 1,91E-30 |
| PEREZ_TP53_TARGETS | 1174 | epithelium) upon expression of TP53 [GeneID=7157] off | 105 | 0,0894 | 3,65E-31 | 3,47E-28 |
| BENPORATH_SUZ12_TARGETS | 1038 | of the Polycomb protein SUZ12 [GeneID=23512] in human embryonic stem cells. | 98 | 0,0944 | 5,56E-31 | 4,42E-28 |
| SCHAEFFER_PROSTATE_DEVELOPMENT_48HR_DN | 428 | females exposed to the androgen dihydrotestosterone [PubChem=10635] for 48 h. | 63 | 0,1472 | 7,77E-31 | 5,28E-28 |
| YOSHIMURA_MAPK8_TARGETS_UP | 1305 | MAPK8 (JNK1) [GeneID=5599]. | 107 | 0,082 | 1,42E-28 | 8,46E-26 |
| NUYTTEN_EZH2_TARGETS_UP | 1037 | knockdown of EZH2 [GeneID=2146] by RNAi. | 94 | 0,0906 | 2,12E-28 | 1,12E-25 |
| BLALOCK_ALZHEIMERS_DISEASE_UP | 1691 | disease. | 123 | 0,0727 | 5,72E-28 | 2,73E-25 |
| WONG_ADULT_TISSUE_STEM_MODULE | 721 | regulated in a compendium of adult tissue stem cells. | 76 | 0,1054 | 3,21E-27 | 1,39E-24 |
| MEISSNER_NPC_HCP_WITH_H3K4ME2 | 491 | H3 dimethylation mark at K4 (H3K4me2) in neural precursor cells (NPC). | 62 | 0,1263 | 1,29E-26 | 5,12E-24 |
| CUI_TCF21_TARGETS_2_DN | 830 | isolated from TCF21 [Gene ID=6943] knockout mice. | 80 | 0,0964 | 5,08E-26 | 1,86E-23 |
| LIM_MAMMARY_STEM_CELL_UP | 489 | mouse and human species. | 60 | 0,1227 | 3,92E-25 | 1,33E-22 |
| MIKKELSEN_NPC_ICP_WITH_H3K4ME3 | 445 | histone H3 trimethylation mark at K4 (H3K4me3) in neural progenitor cells (NPC). | 57 | 0,1281 | 7,01E-25 | 2,22E-22 |
| KINSEY_TARGETS_OF_EWSR1_FLII_FUSION_DN | 329 | Genes down-regulated in TC71 and EWS502 cells (Ewing's sarcoma) by EWSR1-FLI1 [GeneID=2130;2314] as inferred from RNAi knockdown of this fusion protein. | 48 | 0,1459 | 1,29E-23 | 3,83E-21 |
| SMID_BREAST_CANCER_BASAL_DN | 701 | Genes down-regulated in basal subtype of breast cancer samles. | 69 | 0,0984 | 4,22E-23 | 1,18E-20 |
| BENPORATH_PRC2_TARGETS | 652 | Set 'PRC2 targets': Polycomb Repression Complex 2 (PRC) targets; identified by ChIP on chip on human embryonic stem cells as genes that: posess the trimethylated H3K27 mark in their promoters and are bound by SUZ12 [GeneID=23512] and EED [GeneID=8726] Polycomb proteins. | 65 | 0,0997 | 4,00E-22 | 1,06E-19 |
| LEE_BMP2_TARGETS_UP | 745 | Genes up-regulated in uterus upon knockout of BMP2 [GeneID=650]. | 69 | 0,0926 | 1,29E-21 | 3,23E-19 |
| DODD_NASOPHARYNGEAL_CARCINOMA_UP | 1821 | Genes up-regulated in nasopharyngeal carcinoma (NPC) compared to the normal tissue. | 114 | 0,0626 | 1,06E-20 | 2,52E-18 |
| GOBERT_OLIGODENDROCYTE_DIFFERENTIATION_DN | 1080 | Genes down-regulated during differentiation of Oli-Neu cells (oligodendroglial precursor) in response to PD174265 [PubChem=4709]. | 83 | 0,0769 | 1,45E-20 | 3,30E-18 |
| ZWANG_TRANSIENTLY_UP_BY_2ND_EGF_PULSE_ONLY | 1725 | Genes transiently induced only by the second pulse of EGF [GeneID =1950] in 184A1 cells (mammary epithelium). | 109 | 0,0632 | 4,04E-20 | 8,74E-18 |
| GOZGIT_ESR1_TARGETS_DN | 781 | Genes down-regulated in TMX2-28 cells (breast cancer) which do not express ESR1 [GeneID=2099]) compared to the parental MCF7 cells which do. | 68 | 0,0871 | 7,21E-20 | 1,49E-17 |
| GRAESSMANN_APOPTOSIS_BY_DOXORUBICIN_UP | 1142 | Genes up-regulated in ME-A cells (breast cancer) undergoing apoptosis in response to doxorubicin [PubChem=31703]. | 82 | 0,0718 | 1,51E-18 | 2,99E-16 |
| SMID_BREAST_CANCER_LUMINAL_B_DN | 564 | Genes down-regulated in the luminal B subtype of breast cancer. | 55 | 0,0975 | 1,78E-18 | 3,38E-16 |
| SMID_BREAST_CANCER_BASAL_UP | 648 | Genes up-regulated in basal subtype of breast cancer samles. | 59 | 0,091 | 2,63E-18 | 4,82E-16 |
| LIU_PROSTATE_CANCER_DN | 481 | Genes down-regulated in prostate cancer samples. | 50 | 0,104 | 4,61E-18 | 8,12E-16 |
| DURAND_STROMA_S_UP | 297 | Genes up-regulated in the HSC supportive stromal cell lines. | 39 | 0,1313 | 8,07E-18 | 1,37E-15 |
| YANG_BCL3_TARGETS_UP | 364 | Genes up-regulated in neonatal cardiac myocytes upon knockdown of BCL3 [GeneID=602] by RNAi. | 43 | 0,1181 | 9,24E-18 | 1,52E-15 |
| CHICAS_RB1_TARGETS_CONFLUENT | 567 | Genes up-regulated in confluent IMR90 cells (fibroblast) after knockdown of RB1 [GeneID=5925] by RNAi. | 54 | 0,0952 | 1,03E-17 | 1,64E-15 |
| MEISSNER_NPC_HCP_WITH_H3_UNMETHYLATED | 536 | Genes with high-CpG-density promoters (HCP) that have no histone H3 methylation marks in neural precursor cells (NPC). | 52 | 0,097 | 1,91E-17 | 2,88E-15 |
| MIKKELSEN_ES_ICP_WITH_H3K4ME3 | 718 | Genes with intermediate-CpG-density (ICP) promoters bearing histone H3 K4 trimethylation mark (H3K4me3) in embryonic stem cells (ES). | 61 | 0,085 | 1,94E-17 | 2,88E-15 |
| PLASARI_TGFB1_TARGETS_10HR_DN | 244 | Genes down-regulated in MEF cells (embryonic fibroblast) upon stimulation with TGFB1 [GeneID=7040] for 10 h. | 35 | 0,1434 | 2,36E-17 | 3,40E-15 |
| IVANOVA_HEMATOPOIESIS_STEM_CELL_AND_PROGENITO R | 681 | Genes in the expression cluster 'HSC and Progenitors Shared': up-regulated in hematopoietic stem cells (HSC) and progenitors from adult bone marrow and fetal liver. | 59 | 0,0866 | 2,67E-17 | 3,74E-15 |
| NUYTTEN_EZH2_TARGETS_DN | 1024 | Genes down-regulated in PC3 cells (prostate cancer) after knockdown of EZH2 [GeneID=2146] by RNAi. | 74 | 0,0723 | 5,40E-17 | 7,35E-15 |
| ZWANG_TRANSIENTLY_UP_BY_1ST_EGF_PULSE_ONLY | 1839 | Genes transiently induced only by the first pulse of EGF [GeneID =1950] in 184A1 cells (mammary epithelium). | 106 | 0,0576 | 8,61E-17 | 1,13E-14 |

| | | | | | | |
|---|---|---|---|---|---|---|
| CREIGHTON_ENDOCRINE_THERAPY_RESISTANCE_3 | 720 | The 'group 3 set' of genes associated with acquired endocrine therapy resistance in breast tumors expressing ESR1 and ERBB2 [GeneID=2099;2064]. | 60 | 0,0833 | 8,75E-17 | 1,13E-14 |
| MCBRYAN_PUBERTAL_BREAST_4_5WK_UP | 271 | Genes up-regulated during pubertal mammary gland development between week 4 and 5. | 36 | 0,1328 | 1,02E-16 | 1,27E-14 |
| ONDER_CDH1_TARGETS_2_DN | 464 | Genes down-regulated in HMLE cells (immortalized nontransformed mammary epithelium) after E-cadhedrin (CDH1) [GeneID=999] knockdown by RNAi. | 47 | 0,1013 | 1,28E-16 | 1,56E-14 |
| LINDGREN_BLADDER_CANCER_CLUSTER_2B | 392 | Genes specifically up-regulated in Cluster IIb of urothelial cell carcinom (UCC) tumors. | 43 | 0,1097 | 1,43E-16 | 1,71E-14 |
| GRYDER_PAX3FOXO1_ENHANCERS_IN_TADS | 975 | Expressed genes (FPKM>1) associated with high-confidence PAX3-FOXO1 sites with enhancers in primary tumors and cell lines, restricted to those within topological domain boundaries | 71 | 0,0728 | 1,60E-16 | 1,86E-14 |
| RIGGI_EWING_SARCOMA_PROGENITOR_UP | 430 | Genes up-regulated in mesenchymal stem cells (MSC) engineered to express EWS-FLI1 [GeneID=2130;2321] fusion protein. | 45 | 0,1047 | 1,68E-16 | 1,90E-14 |
| WEST_ADRENOCORTICAL_TUMOR_DN | 546 | Down-regulated genes in pediatric adrenocortical tumors (ACT) compared to the normal tissue. | 51 | 0,0934 | 1,87E-16 | 2,08E-14 |
| NABA_MATRISOME | 1028 | Ensemble of genes encoding extracellular matrix and extracellular matrix-associated proteins | 73 | 0,071 | 2,22E-16 | 2,41E-14 |
| BAELDE_DIABETIC_NEPHROPATHY_DN | 434 | Genes down-regulated in glomeruli of kidneys from patients with diabetic nephropathy (type 2 diabetes mellitus). | 45 | 0,1037 | 2,38E-16 | 2,52E-14 |
| RODRIGUES_THYROID_CARCINOMA_POORLY_DIFFERENTIATED_DN | 805 | Genes down-regulated in poorly differentiated thyroid carcinoma (PDTC) compared to normal thyroid tissue. | 63 | 0,0783 | 2,97E-16 | 3,07E-14 |
| BLALOCK_ALZHEIMERS_DISEASE_DN | 1237 | Genes down-regulated in brain from patients with Alzheimer's disease. | 81 | 0,0655 | 4,81E-16 | 4,87E-14 |
| BOQUEST_STEM_CELL_CULTURED_VS_FRESH_UP | 425 | Genes up-regulated in cultured stromal stem cells from adipose tissue, compared to the freshly isolated cells. | 44 | 0,1035 | 5,42E-16 | 5,37E-14 |
| ACEVEDO_FGFR1_TARGETS_IN_PROSTATE_CANCER_MODEL_DN | 308 | Genes down-regulated during prostate cancer progression in the JOCK1 model due to inducible activation of FGFR1 [GeneID=2260] gene in prostate. | 37 | 0,1201 | 1,04E-15 | 1,01E-13 |
| BONOME_OVARIAN_CANCER_SURVIVAL_SUBOPTIMAL_DEBULKING | 510 | Genes whose expression in suboptimally debulked ovarian tumors is associated with survival prognosis. | 48 | 0,0941 | 1,08E-15 | 1,03E-13 |
