## Supplementary Table 2 for "The chromatin reader Dido3 is a regulator of the gene network that controls B cell differentiation"

Differentially expresed transcripts. RNA-seq data.

| Upregulated in Dido1ΔE16 preB | | | | | | |
|---|---|---|---|---|---|---|
| **TranscriptID** | **Symbol** | **logFC** | **logCPM** | **F** | **PValue** | **FDR** |
| **ENSMUST00000044123.1** | Trhr2 | 8,178254997 | 3,689639849 | 291,5462108 | 2,38E-65 | 1,69E-60 |
| **ENSMUST00000036215.7** | Foxj1 | 5,05070799 | 4,24479595 | 240,3549946 | 3,38E-54 | 1,6E-49 |
| **ENSMUST00000069741.3** | E130018O15Rik | 7,328231004 | 3,400752558 | 227,4450916 | 2,97E-51 | 1,06E-46 |
| **ENSMUST00000045068.9** | Cplx3 | 8,373271374 | 3,009840974 | 200,5853343 | 1,8E-45 | 5,13E-41 |
| **ENSMUST00000043315.14** | Serpina3g | 2,670885696 | 6,717498823 | 193,9837055 | 4,37E-44 | 1,04E-39 |
| **ENSMUST00000000199.7** | Ncs1 | 6,476854223 | 3,228629155 | 186,3587587 | 2,01E-42 | 4,1E-38 |
| **ENSMUST00000149378.1** | Gm13398 | 6,422683528 | 3,251844647 | 184,800757 | 5,98E-42 | 1,06E-37 |
| **ENSMUST00000201921.1** | 4930478P22Rik | 4,849938561 | 3,724752284 | 171,9832757 | 2,77E-39 | 4,38E-35 |
| **ENSMUST00000003501.8** | Elavl3 | 6,145498595 | 3,091770125 | 164,8498319 | 9,99E-38 | 1,42E-33 |
| **ENSMUST00000145549.1** | B230312C02Rik | 4,950126862 | 4,267465985 | 215,6493373 | 4,32E-37 | 5,59E-33 |
| **ENSMUST00000100396.3** | 4930407I10Rik | 6,007672773 | 3,012110222 | 152,6974129 | 4,51E-35 | 5,35E-31 |
| **ENSMUST00000203350.1** | Gm44067 | 2,829409924 | 5,51242358 | 148,3735112 | 3,97E-34 | 4,35E-30 |
| **ENSMUST00000098778.2** | Gm10676 | 2,975303294 | 5,190339212 | 145,529287 | 1,66E-33 | 1,69E-29 |
| **ENSMUST00000114973.8** | Kalrn | 4,193488629 | 3,806229123 | 144,124296 | 3,37E-33 | 3,2E-29 |
| **ENSMUST00000145988.8** | Dnhd1 | 2,979992739 | 5,009071296 | 137,1784691 | 1,11E-31 | 9,9E-28 |
| **ENSMUST00000179531.1** | Rnpepl1 | 2,766281975 | 5,374760616 | 135,2117135 | 2,99E-31 | 2,51E-27 |
| **ENSMUST00000181436.1** | Gm26583 | 6,580283705 | 2,667763471 | 135,657234 | 3,65E-31 | 2,89E-27 |
| **ENSMUST00000093832.10** | Lman1l | 8,327862413 | 2,278974964 | 129,007599 | 6,81E-30 | 5,1E-26 |
| **ENSMUST00000060125.6** | Scn4b | 2,689071631 | 5,386890346 | 128,0014945 | 5,13E-29 | 3,65E-25 |
| **ENSMUST00000017637.12** | Igfbp4 | 2,865737935 | 4,840380216 | 120,4695138 | 5,03E-28 | 3,41E-24 |
| **ENSMUST00000057612.8** | Ssc5d | 4,502593079 | 3,19675086 | 117,2099648 | 2,6E-27 | 1,68E-23 |
| **ENSMUST00000187142.2** | Zfp469 | 7,556092546 | 2,19423831 | 116,6247506 | 3,49E-27 | 2,11E-23 |
| **ENSMUST00000046892.9** | Cplx1 | 6,350367745 | 2,47891755 | 116,6594687 | 3,55E-27 | 2,11E-23 |
| **ENSMUST00000207625.1** | A930030B08Rik | 5,274463635 | 2,786384842 | 113,8938095 | 1,38E-26 | 7,88E-23 |
| **ENSMUST00000176061.1** | Gm20681 | 8,12726194 | 2,058727587 | 112,3932968 | 2,95E-26 | 1,61E-22 |
| **ENSMUST00000114574.2** | Glp1r | 6,747750682 | 2,213000516 | 108,8820554 | 1,73E-25 | 9,14E-22 |
| **ENSMUST00000144818.1** | Gm14168 | 3,243553083 | 4,087137291 | 107,3749792 | 3,7E-25 | 1,88E-21 |
| **ENSMUST00000231439.1** | Igll1 | 3,654322209 | 3,723318304 | 109,8775049 | 5,94E-25 | 2,92E-21 |
| **ENSMUST00000103430.1** | Ighj1 | 4,142917982 | 3,161137658 | 101,0360908 | 9,07E-24 | 4,17E-20 |
| **ENSMUST00000030551.10** | Alpl | 2,024903863 | 6,33004764 | 100,9736693 | 9,76E-24 | 4,34E-20 |
| **ENSMUST00000088904.9** | Espnl | 5,864652669 | 2,312617787 | 100,0042741 | 1,53E-23 | 6,59E-20 |
| **ENSMUST00000073935.6** | Gsg1l | 4,583973176 | 2,863310035 | 99,65603652 | 1,82E-23 | 7,63E-20 |
| **ENSMUST00000231330.1** | Gm35455 | 7,257248574 | 1,915417082 | 96,70873365 | 8,07E-23 | 3,28E-19 |
| **ENSMUST00000044970.6** | Mgat3 | 7,248207023 | 1,912963946 | 96,40751334 | 9,39E-23 | 3,72E-19 |
| **ENSMUST00000080368.12** | Atp8a2 | 4,772741967 | 2,689835341 | 95,81534358 | 1,27E-22 | 4,81E-19 |
| **ENSMUST00000045085.7** | Grin3b | 3,645437838 | 3,469184124 | 95,7902219 | 1,28E-22 | 4,81E-19 |
| **ENSMUST00000044384.4** | Aldh1b1 | 1,75479016 | 7,28316449 | 94,78412823 | 2,15E-22 | 7,85E-19 |
| **ENSMUST00000065587.4** | Ackr3 | 3,26082333 | 3,812359919 | 93,99867194 | 3,47E-22 | 1,24E-18 |
| **ENSMUST00000046383.11** | Tnfsf10 | 2,627373387 | 4,558816641 | 92,17799591 | 1,56E-21 | 5,42E-18 |
| **ENSMUST00000095360.10** | Igf1 | 4,022933516 | 3,03523223 | 89,80536709 | 2,64E-21 | 8,94E-18 |
| **ENSMUST00000144897.1** | Slx1b | 2,44672511 | 4,826217701 | 88,32164445 | 5,58E-21 | 1,85E-17 |
| **ENSMUST00000203263.1** | Gm44066 | 2,643559159 | 4,429725713 | 87,93388583 | 6,79E-21 | 2,2E-17 |

| | | | | | | |
|---|---|---|---|---|---|---|
| **ENSMUST00000009875.4** | Kcnd1 | 3,878502091 | 3,060776876 | 86,71730507 | 1,26E-20 | 3,98E-17 |
| **ENSMUST00000049614.12** | B430306N03Rik | 2,477422298 | 4,677193091 | 85,91677149 | 1,88E-20 | 5,83E-17 |
| **ENSMUST00000070328.9** | Sh2d4b | 2,756022319 | 4,209991815 | 84,79227113 | 3,32E-20 | 1,01E-16 |
| **ENSMUST00000140833.1** | Gm15728 | 5,546501016 | 2,104448451 | 82,37768727 | 1,13E-19 | 3,28E-16 |
| **ENSMUST00000000188.11** | Ccnd2 | 1,818841158 | 6,329194025 | 81,45457634 | 1,83E-19 | 5,21E-16 |
| **ENSMUST00000057831.7** | Cilp2 | 5,029747674 | 2,245706746 | 81,06735435 | 2,19E-19 | 6,11E-16 |
| **ENSMUST00000180086.2** | H1f0 | 1,595263421 | 7,333076762 | 78,90285205 | 6,54E-19 | 1,79E-15 |
| **ENSMUST00000022699.9** | Gfra2 | 2,926129741 | 3,827212439 | 78,58993845 | 7,71E-19 | 2,07E-15 |
| **ENSMUST00000040821.4** | Heyl | 2,045638174 | 5,54571685 | 79,12551615 | 1,17E-18 | 3,07E-15 |
| **ENSMUST00000054491.5** | Sox18 | 8,857512753 | 1,34795791 | 77,4372545 | 1,37E-18 | 3,56E-15 |
| **ENSMUST00000179520.1** | Ighd4-1 | 4,181337013 | 2,652040852 | 77,36235869 | 1,43E-18 | 3,63E-15 |
| **ENSMUST00000025830.8** | Apba1 | 2,83487618 | 3,933545204 | 77,33293058 | 1,47E-18 | 3,68E-15 |
| **ENSMUST00000033054.9** | Adm | 2,745857335 | 4,051061047 | 77,31621369 | 1,52E-18 | 3,73E-15 |
| **ENSMUST00000105572.2** | Perm1 | 2,736342512 | 4,010776099 | 76,00926352 | 2,83E-18 | 6,83E-15 |
| **ENSMUST00000030420.8** | Epha8 | 8,802780653 | 1,308029304 | 75,36574302 | 3,92E-18 | 9,31E-15 |
| **ENSMUST00000145401.7** | Il9r | 3,600406092 | 3,923795062 | 100,1664368 | 6,83E-18 | 1,59E-14 |
| **ENSMUST00000166193.8** | Igfn1 | 6,853215762 | 1,529374109 | 74,07841672 | 7,53E-18 | 1,73E-14 |
| **ENSMUST00000020537.8** | Nsg2 | 1,765745532 | 6,112490555 | 71,46610797 | 2,83E-17 | 6,39E-14 |
| **ENSMUST00000206494.1** | Unc45a | 3,446151321 | 3,057578002 | 70,31115962 | 5,08E-17 | 1,13E-13 |
| **ENSMUST00000027533.8** | Klhl30 | 6,312169778 | 1,524034391 | 68,46552634 | 1,29E-16 | 2,79E-13 |
| **ENSMUST00000159547.2** | Vamp1 | 1,542047785 | 6,889653481 | 67,41663009 | 2,2E-16 | 4,68E-13 |
| **ENSMUST00000185943.1** | 5031425E22Rik | 3,323723001 | 3,043311935 | 65,79335522 | 5,04E-16 | 1,06E-12 |
| **ENSMUST00000028689.3** | Lrp4 | 3,288327318 | 3,094569561 | 65,64168988 | 5,42E-16 | 1,12E-12 |
| **ENSMUST00000051765.8** | Glp2r | 4,982508838 | 1,935230947 | 65,5190259 | 5,8E-16 | 1,18E-12 |
| **ENSMUST00000017561.14** | Plxdc1 | 2,130467259 | 4,719182841 | 64,85862685 | 8,06E-16 | 1,62E-12 |
| **ENSMUST00000075317.11** | Pdzd2 | 1,577930631 | 6,475284059 | 63,91053675 | 1,41E-15 | 2,79E-12 |
| **ENSMUST00000236745.1** | Ubash3a | 3,015582757 | 3,297585315 | 63,06607012 | 2E-15 | 3,91E-12 |
| **ENSMUST00000018711.14** | Gabarap | 2,282096628 | 4,308942553 | 62,17482126 | 3,15E-15 | 6,06E-12 |
| **ENSMUST00000127305.1** | Epn3 | 8,460520852 | 1,009187845 | 61,83380794 | 3,74E-15 | 7,11E-12 |
| **ENSMUST00000097648.5** | Ramp1 | 2,138698729 | 4,519124664 | 60,58985181 | 7,04E-15 | 1,32E-11 |
| **ENSMUST00000192126.1** | Gm18407 | 2,775747462 | 3,512532719 | 59,74526766 | 1,08E-14 | 2E-11 |
| **ENSMUST00000102942.7** | Psd4 | 1,726891331 | 5,629282353 | 58,2842288 | 2,27E-14 | 4,15E-11 |
| **ENSMUST00000126198.2** | Fam78b | 5,812836728 | 1,343676636 | 58,01062587 | 2,61E-14 | 4,71E-11 |
| **ENSMUST00000196046.1** | Gm43364 | 6,479984579 | 1,171433088 | 57,88118167 | 2,79E-14 | 4,97E-11 |
| **ENSMUST00000097474.8** | Rcsd1 | 2,233583036 | 4,159368437 | 54,99921882 | 1,21E-13 | 2,12E-10 |
| **ENSMUST00000132422.1** | Gps1 | 3,284351465 | 2,758402058 | 54,54650402 | 1,52E-13 | 2,64E-10 |
| **ENSMUST00000053856.5** | Pcdhb17 | 3,31656198 | 2,705065885 | 53,65228999 | 2,4E-13 | 4,06E-10 |
| **ENSMUST00000132080.1** | Tmem259 | 2,905692276 | 3,118682753 | 53,42706116 | 2,69E-13 | 4,5E-10 |
| **ENSMUST00000052528.4** | Gm9847 | 3,624529389 | 2,40044788 | 52,78942758 | 3,73E-13 | 6,17E-10 |
| **ENSMUST00000026315.7** | Dnase1l3 | 1,541661805 | 6,019372997 | 52,69721816 | 3,9E-13 | 6,38E-10 |
| **ENSMUST00000137906.1** | Arid5a | 1,883715516 | 4,820906235 | 52,59156905 | 4,11E-13 | 6,66E-10 |
| **ENSMUST00000103091.8** | Elmo2 | 3,524071926 | 2,440161058 | 52,32854761 | 4,7E-13 | 7,52E-10 |
| **ENSMUST00000075827.4** | Jag2 | 2,709793846 | 3,29670242 | 51,52895184 | 7,06E-13 | 1,12E-09 |
| **ENSMUST00000016172.8** | Celsr1 | 3,06916754 | 2,851172513 | 51,14074038 | 8,61E-13 | 1,35E-09 |
| **ENSMUST00000134427.1** | Gm11613 | 2,334296872 | 3,824140741 | 50,75561061 | 1,05E-12 | 1,6E-09 |
| **ENSMUST00000108567.8** | Zfp444 | 2,085246088 | 4,243471562 | 50,20504135 | 1,39E-12 | 2,1E-09 |
| **ENSMUST00000029658.13** | Enpep | 2,509797611 | 3,483001415 | 49,1351093 | 2,39E-12 | 3,59E-09 |
| **ENSMUST00000060474.13** | Septin6 | 1,537678051 | 5,80830831 | 48,91053335 | 2,68E-12 | 3,98E-09 |

| | | | | | | |
|---|---|---|---|---|---|---|
| **ENSMUST00000000153.8** | Gna12 | 1,332689888 | 6,748772682 | 48,70528104 | 2,98E-12 | 4,37E-09 |
| **ENSMUST00000071135.5** | Tubb4a | 2,778146887 | 3,099110788 | 48,58844318 | 3,16E-12 | 4,59E-09 |
| **ENSMUST00000211765.1** | Nucb1 | 1,823622664 | 4,731981699 | 47,86251092 | 4,58E-12 | 6,52E-09 |
| **ENSMUST00000202026.3** | 4930478P22Rik | 4,326020601 | 1,697352809 | 47,61659668 | 5,2E-12 | 7,34E-09 |
| **ENSMUST00000030585.7** | A3galt2 | 3,901896341 | 1,92571106 | 47,4104692 | 5,78E-12 | 8,07E-09 |
| **ENSMUST00000020768.3** | Pgam2 | 1,511219167 | 5,805748496 | 47,19041525 | 6,45E-12 | 8,83E-09 |
| **ENSMUST00000030677.6** | Map3k6 | 5,128849384 | 1,20890097 | 47,07878579 | 6,83E-12 | 9,26E-09 |
| **ENSMUST00000025805.7** | Cnih2 | 4,390761895 | 1,571238692 | 46,78615298 | 7,93E-12 | 1,06E-08 |
| **ENSMUST00000039752.3** | Slc16a8 | 4,955339846 | 1,298610136 | 46,7491308 | 8,08E-12 | 1,08E-08 |
| **ENSMUST00000211939.1** | Polr2c | 3,127553302 | 2,612228058 | 46,30081159 | 1,02E-11 | 1,34E-08 |
| **ENSMUST00000024575.7** | Rps6ka2 | 1,724961592 | 4,949365839 | 46,22813017 | 1,05E-11 | 1,38E-08 |
| **ENSMUST00000127208.7** | Lrrc14 | 2,921898408 | 2,831551061 | 46,08399964 | 1,13E-11 | 1,47E-08 |
| **ENSMUST00000037007.3** | Evpl | 3,199786031 | 2,525025391 | 45,620841 | 1,44E-11 | 1,85E-08 |
| **ENSMUST00000055619.4** | Hic1 | 3,146427583 | 2,54680267 | 45,31331751 | 1,68E-11 | 2,1E-08 |
| **ENSMUST00000132562.1** | Wrap73 | 3,651890768 | 2,045044936 | 44,96172096 | 2,01E-11 | 2,47E-08 |
| **ENSMUST00000224954.1** | Daam2 | 2,639852539 | 3,123062049 | 44,50121968 | 2,54E-11 | 3,1E-08 |
| **ENSMUST00000047282.11** | Mthfsd | 1,706383836 | 4,895833979 | 44,40456761 | 2,67E-11 | 3,23E-08 |
| **ENSMUST00000046937.3** | Tssk1 | 4,152835828 | 1,611894621 | 43,99331675 | 3,3E-11 | 3,95E-08 |
| **ENSMUST00000232008.1** | 4933432I09Rik | 4,123826935 | 1,644247453 | 43,5698768 | 4,09E-11 | 4,86E-08 |
| **ENSMUST00000040059.8** | Hyal3 | 4,306498157 | 1,494729643 | 43,48327016 | 4,28E-11 | 5,04E-08 |
| **ENSMUST00000055872.2** | Galr2 | 3,091181839 | 2,541521959 | 43,07381194 | 5,28E-11 | 6,11E-08 |
| **ENSMUST00000109764.7** | Nfix | 1,7474685 | 4,639411263 | 42,50577762 | 7,05E-11 | 8,1E-08 |
| **ENSMUST00000120430.1** | Gm7901 | 1,119022712 | 7,808153921 | 42,14990359 | 8,46E-11 | 9,64E-08 |
| **ENSMUST00000184613.1** | Mrip-ps | 1,188132892 | 7,181988937 | 42,07389417 | 8,81E-11 | 9,97E-08 |
| **ENSMUST00000122029.1** | Gm6939 | 3,104980149 | 2,396644455 | 41,14392486 | 1,42E-10 | 1,58E-07 |
| **ENSMUST00000047309.5** | Nat14 | 2,795271917 | 2,755541996 | 40,68164561 | 1,79E-10 | 1,98E-07 |
| **ENSMUST00000201142.3** | 1700028E10Rik | 2,743697883 | 2,808371646 | 40,61036861 | 1,86E-10 | 2,04E-07 |
| **ENSMUST00000053926.11** | Gm49486 | 4,520082979 | 1,257922682 | 40,51833229 | 1,95E-10 | 2,1E-07 |
| **ENSMUST00000032877.10** | Ddias | 1,522241681 | 5,29710636 | 40,41477923 | 2,06E-10 | 2,21E-07 |
| **ENSMUST00000165838.8** | Metrn | 2,013891208 | 3,934040724 | 40,28601272 | 2,2E-10 | 2,33E-07 |
| **ENSMUST00000238352.1** | Gm10570 | 4,65685676 | 1,140663365 | 40,24497226 | 2,24E-10 | 2,37E-07 |
| **ENSMUST00000099373.11** | Cnnm2 | 3,11348923 | 2,358574148 | 40,20048908 | 2,29E-10 | 2,4E-07 |
| **ENSMUST00000206275.1** | Bola2 | 2,04896394 | 3,857536965 | 40,12932128 | 2,38E-10 | 2,47E-07 |
| **ENSMUST00000086281.4** | Zfp599 | 2,204170722 | 3,581558944 | 40,03868374 | 2,49E-10 | 2,57E-07 |
| **ENSMUST00000095076.9** | Epb41l4b | 3,467940505 | 1,959970582 | 39,68324766 | 2,99E-10 | 3,06E-07 |
| **ENSMUST00000112682.3** | Slc25a18 | 1,746638239 | 4,461506871 | 39,37063282 | 3,51E-10 | 3,56E-07 |
| **ENSMUST00000057311.3** | Sfn | 1,293032336 | 6,207664903 | 39,23106449 | 3,79E-10 | 3,8E-07 |
| **ENSMUST00000225289.1** | Rab24 | 3,467489401 | 1,933190752 | 38,68112905 | 4,99E-10 | 4,94E-07 |
| **ENSMUST00000153578.7** | Tspoap1 | 1,991732965 | 3,934592249 | 39,14459395 | 5,11E-10 | 5,02E-07 |
| **ENSMUST00000168386.8** | Prr36 | 4,461707258 | 1,163163503 | 38,15753377 | 6,53E-10 | 6,33E-07 |
| **ENSMUST00000086040.5** | F5 | 1,239453192 | 7,159198127 | 41,20653877 | 7,58E-10 | 7,2E-07 |
| **ENSMUST00000204059.2** | Add2 | 3,308950451 | 2,010560846 | 37,76298621 | 7,99E-10 | 7,54E-07 |
| **ENSMUST00000051442.6** | Pcdhb16 | 2,981830872 | 2,384961332 | 37,70910916 | 8,22E-10 | 7,7E-07 |
| **ENSMUST00000179719.1** | Hyi | 2,303165648 | 3,285830862 | 37,63990033 | 8,51E-10 | 7,92E-07 |
| **ENSMUST00000194216.1** | Gm37795 | 2,443973485 | 3,052818063 | 37,35251713 | 9,87E-10 | 8,95E-07 |
| **ENSMUST00000201742.1** | Gm43860 | 1,504986218 | 5,056842678 | 36,35552456 | 1,65E-09 | 1,47E-06 |
| **ENSMUST00000039840.14** | Enpp6 | 2,810178276 | 2,502889818 | 36,13162121 | 1,85E-09 | 1,64E-06 |
| **ENSMUST00000028403.2** | Cybrd1 | 2,701860557 | 2,708552249 | 36,43646512 | 1,87E-09 | 1,65E-06 |

| | | | | | | |
|---|---|---|---|---|---|---|
| **ENSMUST00000056890.9** | Fbxl22 | 1,324652146 | 5,787102885 | 35,89699984 | 2,08E-09 | 1,83E-06 |
| **ENSMUST00000030142.3** | Epb41l4b | 1,114166377 | 7,034874192 | 35,83237583 | 2,15E-09 | 1,88E-06 |
| **ENSMUST00000204971.1** | Pyroxd1 | 3,535965666 | 1,681534547 | 35,46673706 | 2,6E-09 | 2,23E-06 |
| **ENSMUST00000055436.4** | Hpdl | 4,395973385 | 1,110148268 | 35,18551657 | 3E-09 | 2,56E-06 |
| **ENSMUST00000217282.1** | Gm38431 | 3,047412451 | 2,155405091 | 35,14967029 | 3,05E-09 | 2,59E-06 |
| **ENSMUST00000026985.8** | Cplx2 | 0,786366167 | 11,12044583 | 35,08636215 | 3,16E-09 | 2,66E-06 |
| **ENSMUST00000062117.13** | Rap2a | 1,823566263 | 4,054593856 | 35,0695298 | 3,18E-09 | 2,67E-06 |
| **ENSMUST00000150442.1** | Gm11642 | 3,953466338 | 1,331947062 | 34,99043619 | 3,32E-09 | 2,76E-06 |
| **ENSMUST00000115421.2** | Steap4 | 0,906258524 | 9,153220041 | 34,91076411 | 3,45E-09 | 2,86E-06 |
| **ENSMUST00000023259.14** | Lynx1 | 0,872210902 | 9,622204881 | 34,75964575 | 3,76E-09 | 3,06E-06 |
| **ENSMUST00000050519.7** | Elfn1 | 4,348535247 | 1,06872759 | 33,9158706 | 5,77E-09 | 4,64E-06 |
| **ENSMUST00000238703.1** | Gm50464 | 3,843958267 | 1,322656373 | 33,29999737 | 7,9E-09 | 6,22E-06 |
| **ENSMUST00000180612.2** | 9330175E14Rik | 1,438984143 | 5,022930753 | 32,87358614 | 9,84E-09 | 7,7E-06 |
| **ENSMUST00000128342.1** | Gm16576 | 1,670149231 | 4,238729656 | 32,30430049 | 1,32E-08 | 1,02E-05 |
| **ENSMUST00000103105.9** | Aoc3 | 3,313840055 | 1,722766459 | 32,07082628 | 1,49E-08 | 1,15E-05 |
| **ENSMUST00000077502.4** | Dqx1 | 1,869603207 | 3,750285324 | 31,75555964 | 1,75E-08 | 1,33E-05 |
| **ENSMUST00000066330.14** | Mprip | 1,240870191 | 5,813474083 | 31,70793684 | 1,79E-08 | 1,36E-05 |
| **ENSMUST00000123349.1** | Gramd4 | 1,580677503 | 4,425720133 | 31,62199216 | 1,87E-08 | 1,41E-05 |
| **ENSMUST00000027809.7** | Opn3 | 1,488250879 | 4,718897142 | 31,57717888 | 1,92E-08 | 1,42E-05 |
| **ENSMUST00000101339.7** | Nhsl2 | 1,775886223 | 3,947564817 | 31,42541421 | 2,07E-08 | 1,52E-05 |
| **ENSMUST00000201164.1** | 4930553P18Rik | 2,691706319 | 2,400043714 | 31,37788503 | 2,12E-08 | 1,55E-05 |
| **ENSMUST00000066386.5** | Lysmd1 | 1,916718853 | 3,643093553 | 31,3707699 | 2,13E-08 | 1,55E-05 |
| **ENSMUST00000192173.1** | Gm37233 | 1,756222894 | 3,963099576 | 31,10297182 | 2,45E-08 | 1,76E-05 |
| **ENSMUST00000209248.1** | Rnf223 | 2,456071228 | 2,702815818 | 31,03311763 | 2,54E-08 | 1,82E-05 |
| **ENSMUST00000039178.11** | Tnn | 3,021037311 | 1,966058195 | 30,76333939 | 2,92E-08 | 2,08E-05 |
| **ENSMUST00000165968.1** | Serpina3e-ps | 2,644285216 | 2,411907374 | 30,63693545 | 3,11E-08 | 2,21E-05 |
| **ENSMUST00000027952.11** | Plxna2 | 2,451827751 | 2,679388168 | 30,55589348 | 3,25E-08 | 2,29E-05 |
| **ENSMUST00000142367.7** | Palm3 | 3,415190644 | 1,474732627 | 30,13047218 | 4,05E-08 | 2,8E-05 |
| **ENSMUST00000231640.1** | Gm49745 | 2,835067494 | 2,0716474 | 29,54989546 | 5,45E-08 | 3,75E-05 |
| **ENSMUST00000087497.10** | Col11a2 | 1,918409716 | 3,511668596 | 29,49297622 | 5,61E-08 | 3,83E-05 |
| **ENSMUST00000119494.2** | Plk-ps1 | 1,649705765 | 4,078484833 | 29,00786005 | 7,21E-08 | 4,87E-05 |
| **ENSMUST00000048923.6** | Spred3 | 2,233625227 | 2,883617794 | 28,47234464 | 9,51E-08 | 6,22E-05 |
| **ENSMUST00000226062.1** | Psd | 1,382994821 | 4,796589922 | 28,04201831 | 1,19E-07 | 7,62E-05 |
| **ENSMUST00000110052.1** | Ocel1 | 1,202401787 | 5,630785468 | 28,02148255 | 1,2E-07 | 7,67E-05 |
| **ENSMUST00000159283.7** | Manf | 2,099437434 | 3,071940068 | 27,73969493 | 1,39E-07 | 8,75E-05 |
| **ENSMUST00000232332.2** | Gm1043 | 1,515355313 | 4,323381884 | 27,56396107 | 1,52E-07 | 9,5E-05 |
| **ENSMUST00000051259.9** | Adgrg3 | 3,149464432 | 1,582950951 | 27,5342098 | 1,54E-07 | 9,6E-05 |
| **ENSMUST00000036248.12** | Pmepa1 | 1,889435097 | 3,406260219 | 27,11245222 | 1,92E-07 | 0,000116893 |
| **ENSMUST00000130268.7** | Mypop | 1,651725153 | 3,9274394 | 26,93075453 | 2,11E-07 | 0,000126255 |
| **ENSMUST00000108288.8** | Lrfn1 | 2,648980494 | 2,118020507 | 26,87884245 | 2,17E-07 | 0,00012915 |
| **ENSMUST00000232529.1** | Car15 | 2,190762771 | 2,816033678 | 26,49825842 | 2,64E-07 | 0,000154042 |
| **ENSMUST00000093369.4** | Nefh | 3,255859372 | 1,408825797 | 26,37595979 | 2,81E-07 | 0,000162776 |
| **ENSMUST00000123325.8** | Ankrd33b | 1,385163378 | 4,612731563 | 26,3481133 | 2,85E-07 | 0,000163808 |
| **ENSMUST00000180852.1** | 2610037D02Rik | 2,883952188 | 1,80721041 | 26,32098562 | 2,89E-07 | 0,000165457 |
| **ENSMUST00000204482.2** | Tmem176a | 2,954310878 | 1,69945943 | 26,26396613 | 2,98E-07 | 0,000169733 |
| **ENSMUST00000058479.6** | Drc7 | 2,627568213 | 2,080968669 | 25,83238963 | 3,73E-07 | 0,000208904 |
| **ENSMUST00000149932.1** | Gm13184 | 3,001670523 | 1,615375774 | 25,43483112 | 4,58E-07 | 0,000252715 |
| **ENSMUST00000006669.5** | Pdk1 | 1,598024162 | 3,934299237 | 25,38804423 | 4,69E-07 | 0,000258066 |

| | | | | | | |
|---|---|---|---|---|---|---|
| **ENSMUST00000023072.6** | Parvb | 1,80953032 | 3,435541445 | 25,33830344 | 4,82E-07 | 0,00026396 |
| **ENSMUST00000177908.1** | Cfap73 | 1,863204877 | 3,343751007 | 25,25257978 | 5,03E-07 | 0,0002737 |
| **ENSMUST00000054487.9** | Ajuba | 3,432751962 | 1,148390741 | 25,21224091 | 5,14E-07 | 0,000278234 |
| **ENSMUST00000125447.2** | Macf1 | 1,488815289 | 4,207830645 | 25,15140066 | 5,31E-07 | 0,000286305 |
| **ENSMUST00000024860.8** | Ehd3 | 1,300951319 | 4,815575882 | 24,90150471 | 6,04E-07 | 0,000321987 |
| **ENSMUST00000110218.8** | Spef1 | 1,779618352 | 3,473243744 | 24,82857175 | 6,27E-07 | 0,000333154 |
| **ENSMUST00000193564.1** | Gm37736 | 2,774531055 | 1,80628983 | 24,78582995 | 6,41E-07 | 0,000339357 |
| **ENSMUST00000041124.12** | Zfp704 | 1,471828907 | 4,208156849 | 24,66828683 | 6,82E-07 | 0,000359985 |
| **ENSMUST00000028223.8** | Kynu | 1,607251987 | 3,812534738 | 24,11986018 | 9,05E-07 | 0,00046902 |
| **ENSMUST00000065793.11** | Phgdh | 0,978825536 | 6,451178839 | 24,04023603 | 9,44E-07 | 0,00048529 |
| **ENSMUST00000163139.7** | Plxna1 | 1,300548345 | 4,708694186 | 23,97406199 | 9,77E-07 | 0,000498658 |
| **ENSMUST00000104937.1** | Ankrd63 | 3,182732563 | 1,266945918 | 23,91952368 | 1E-06 | 0,000509172 |
| **ENSMUST00000003154.6** | Efna2 | 2,856305497 | 1,64539989 | 23,91329801 | 1,01E-06 | 0,000509172 |
| **ENSMUST00000124126.1** | Polr1c | 3,319268229 | 1,183690409 | 23,8860097 | 1,02E-06 | 0,000514616 |
| **ENSMUST00000020549.3** | Gzmm | 2,539601573 | 2,0484197 | 23,59066076 | 1,19E-06 | 0,000593671 |
| **ENSMUST00000022921.6** | Angpt1 | 1,366028426 | 4,423730217 | 23,43669635 | 1,29E-06 | 0,000638663 |
| **ENSMUST00000133221.2** | Trp53cor1 | 3,098395699 | 1,320261454 | 23,37125186 | 1,34E-06 | 0,000656207 |
| **ENSMUST00000218571.1** | Map2k2 | 2,343529236 | 2,300906166 | 23,26372174 | 1,41E-06 | 0,000689183 |
| **ENSMUST00000119613.1** | Gm5939 | 1,081571173 | 5,721270725 | 23,24257768 | 1,43E-06 | 0,000694423 |
| **ENSMUST00000094451.3** | Gpr157 | 2,000914006 | 2,899214486 | 23,10041596 | 1,54E-06 | 0,00074516 |
| **ENSMUST00000045713.3** | Nacad | 1,397895867 | 4,270906699 | 22,90602162 | 1,7E-06 | 0,000816118 |
| **ENSMUST00000059026.9** | Abi3 | 2,459223566 | 2,102308684 | 22,78780396 | 1,81E-06 | 0,000856356 |
| **ENSMUST00000004036.5** | Efnb3 | 1,593601349 | 3,729015516 | 22,77231922 | 1,82E-06 | 0,000860425 |
| **ENSMUST00000200257.1** | Gm19710 | 2,389135468 | 2,183344783 | 22,75505761 | 1,84E-06 | 0,000865322 |
| **ENSMUST00000152138.7** | Casc1 | 3,209823198 | 1,201112581 | 22,7080501 | 1,89E-06 | 0,000883833 |
| **ENSMUST00000187230.1** | Begain | 3,058379859 | 1,290756938 | 22,67051025 | 1,92E-06 | 0,000898314 |
| **ENSMUST00000217354.1** | Ppp2r3d | 1,490085117 | 3,979783517 | 22,51386827 | 2,09E-06 | 0,000965129 |
| **ENSMUST00000154277.1** | Galr2 | 2,720036002 | 1,686457289 | 22,47970309 | 2,12E-06 | 0,000979268 |
| **ENSMUST00000208686.1** | Prcp | 2,196527545 | 2,490823108 | 22,42829885 | 2,18E-06 | 0,001002584 |
| **ENSMUST00000122882.1** | 2210406O10Rik | 3,271197846 | 1,046784841 | 22,15524491 | 2,52E-06 | 0,001144697 |
| **ENSMUST00000014118.3** | Mcemp1 | 2,659118377 | 1,757489807 | 22,08215972 | 2,61E-06 | 0,001182165 |
| **ENSMUST00000231165.1** | Nfam1 | 1,226158307 | 4,800452911 | 21,91974 | 2,84E-06 | 0,001273771 |
| **ENSMUST00000159004.1** | Mrgpra2a | 2,648018483 | 1,876412857 | 22,36282257 | 2,92E-06 | 0,001304577 |
| **ENSMUST00000026886.7** | Itih5 | 1,178633588 | 5,016252047 | 21,81641371 | 3E-06 | 0,001331674 |
| **ENSMUST00000211974.1** | Adgrg3 | 2,726435361 | 1,610165285 | 21,80186389 | 3,02E-06 | 0,001337643 |
| **ENSMUST00000234131.1** | Slc8a1 | 2,580137573 | 1,850693752 | 21,79559037 | 3,03E-06 | 0,001337869 |
| **ENSMUST00000033495.14** | Pim2 | 0,844940609 | 7,347218913 | 21,6973006 | 3,19E-06 | 0,001398219 |
| **ENSMUST00000070084.10** | Ticam2 | 1,160760069 | 5,086970346 | 21,67381182 | 3,23E-06 | 0,001408107 |
| **ENSMUST00000074408.6** | Ifnlr1 | 2,184659642 | 2,438989006 | 21,6521755 | 3,27E-06 | 0,001419737 |
| **ENSMUST00000082432.4** | Dio2 | 2,000961368 | 2,76544012 | 21,60887215 | 3,34E-06 | 0,001443352 |
| **ENSMUST00000193350.1** | Gm37120 | 1,369415633 | 4,236353842 | 21,57156364 | 3,41E-06 | 0,001463789 |
| **ENSMUST00000041093.5** | Creb3l2 | 0,815078195 | 7,689664066 | 21,57031547 | 3,41E-06 | 0,001463789 |
| **ENSMUST00000028780.3** | Chac1 | 2,542078207 | 1,852268354 | 21,47526434 | 3,59E-06 | 0,001528955 |
| **ENSMUST00000142665.1** | Wrap73 | 1,614515928 | 3,569008251 | 21,40444732 | 3,72E-06 | 0,001581742 |
| **ENSMUST00000060524.10** | Trim10 | 2,23280333 | 2,290065577 | 21,04537315 | 4,5E-06 | 0,001881433 |
| **ENSMUST00000069817.14** | Prrc2b | 0,921954593 | 6,411183435 | 20,99414127 | 4,62E-06 | 0,001923849 |
| **ENSMUST00000193194.1** | AA914427 | 3,146122504 | 1,118916616 | 20,98483954 | 4,63E-06 | 0,001923849 |
| **ENSMUST00000169652.2** | Tifab | 1,622130984 | 3,485237747 | 20,70558945 | 5,36E-06 | 0,002193104 |

| | | | | | | |
|---|---|---|---|---|---|---|
| **ENSMUST00000237716.1** | Gramd3 | 1,704791207 | 3,267133208 | 20,41862597 | 6,22E-06 | 0,002504222 |
| **ENSMUST00000159977.1** | Pcmt1 | 2,424066666 | 1,933994746 | 20,35431869 | 6,44E-06 | 0,002568039 |
| **ENSMUST00000024774.13** | Guca1b | 2,026143064 | 2,60078282 | 20,21724586 | 6,91E-06 | 0,002751059 |
| **ENSMUST00000019068.6** | Alox15 | 2,4872212 | 1,833073366 | 20,19116051 | 7,41E-06 | 0,002922246 |
| **ENSMUST00000105545.11** | Phactr2 | 1,552111795 | 3,596681038 | 20,06800061 | 7,48E-06 | 0,002941441 |
| **ENSMUST00000069324.6** | Zfp580 | 1,759429981 | 3,099546922 | 19,90546691 | 8,14E-06 | 0,003167394 |
| **ENSMUST00000182751.1** | Rbpj-ps3 | 0,962880195 | 5,957685839 | 19,81859605 | 8,52E-06 | 0,003305615 |
| **ENSMUST00000132882.1** | Hyi | 2,585618917 | 1,637535361 | 19,80162095 | 8,59E-06 | 0,003326038 |
| **ENSMUST00000040128.11** | Atp8b4 | 2,417728374 | 2,038192274 | 20,24541723 | 8,85E-06 | 0,003407539 |
| **ENSMUST00000035672.4** | Ppl | 2,60080229 | 1,597221282 | 19,6156583 | 9,47E-06 | 0,003626536 |
| **ENSMUST00000233692.1** | Lrrc73 | 2,578944247 | 1,6071816 | 19,47485064 | 1,02E-05 | 0,003872694 |
| **ENSMUST00000050020.7** | Jaml | 2,252198496 | 2,089603582 | 19,1849666 | 1,19E-05 | 0,004459996 |
| **ENSMUST00000089510.4** | Cenpb | 0,691954558 | 8,828774413 | 19,12108393 | 1,23E-05 | 0,004575567 |
| **ENSMUST00000139210.7** | Socs2 | 2,851757636 | 1,221918022 | 18,96721204 | 1,33E-05 | 0,004908384 |
| **ENSMUST00000050234.3** | Jrk | 1,078053278 | 5,135798253 | 18,95126244 | 1,34E-05 | 0,004924073 |
| **ENSMUST00000127984.8** | Cbfa2t3 | 0,691218048 | 8,775555733 | 18,91562972 | 1,37E-05 | 0,005004006 |
| **ENSMUST00000173127.2** | Gm9574 | 2,090222372 | 2,324130758 | 18,82405284 | 1,43E-05 | 0,005183463 |
| **ENSMUST00000111881.3** | Nfkb2 | 1,812067868 | 2,871430599 | 18,70725661 | 1,52E-05 | 0,005496874 |
| **ENSMUST00000180252.2** | Tmem151b | 2,902782205 | 1,092301247 | 18,5867386 | 1,62E-05 | 0,005826049 |
| **ENSMUST00000023487.4** | Arhgap31 | 1,011545585 | 5,439308958 | 18,44933609 | 1,75E-05 | 0,006158626 |
| **ENSMUST00000172910.2** | Crybg3 | 1,05867018 | 5,158532199 | 18,40626317 | 1,78E-05 | 0,006278167 |
| **ENSMUST00000113658.7** | Gfpt1 | 1,043398623 | 5,24004564 | 18,38379541 | 1,81E-05 | 0,006336984 |
| **ENSMUST00000190196.4** | Prob1 | 1,596870647 | 3,300553402 | 18,32104184 | 1,87E-05 | 0,006469501 |
| **ENSMUST00000105617.7** | Ipcef1 | 0,864975741 | 6,386381639 | 18,29696272 | 1,89E-05 | 0,006529436 |
| **ENSMUST00000099858.3** | Prep | 0,658705716 | 9,17719194 | 18,2942255 | 1,89E-05 | 0,006529436 |
| **ENSMUST00000040971.13** | Capn5 | 0,965678577 | 5,691155889 | 18,26944101 | 1,92E-05 | 0,006598963 |
| **ENSMUST00000149137.2** | Gm13270 | 2,675173489 | 1,389537889 | 18,17959909 | 2,01E-05 | 0,00688442 |
| **ENSMUST00000044195.5** | Tmc7 | 0,972133477 | 5,633514391 | 18,17140072 | 2,02E-05 | 0,006897539 |
| **ENSMUST00000135229.7** | Dgcr2 | 1,443464753 | 3,684723069 | 18,13202921 | 2,06E-05 | 0,007024777 |
| **ENSMUST00000082152.4** | Ube2o | 1,238773818 | 4,283131946 | 18,00507573 | 2,2E-05 | 0,007447531 |
| **ENSMUST00000050511.6** | Kynu | 2,18494778 | 2,078697419 | 18,0026277 | 2,21E-05 | 0,007447531 |
| **ENSMUST00000004565.14** | Ralb | 1,261448967 | 4,195447806 | 17,86523433 | 2,37E-05 | 0,007929837 |
| **ENSMUST00000180430.1** | Ksr2 | 2,687576574 | 1,31561539 | 17,82193914 | 2,43E-05 | 0,008074414 |
| **ENSMUST00000024779.14** | Usp49 | 0,921717753 | 5,877910248 | 17,66871917 | 2,63E-05 | 0,008610777 |
| **ENSMUST00000165559.2** | Ctif | 2,102857236 | 2,200264373 | 17,60586527 | 2,72E-05 | 0,008870608 |
| **ENSMUST00000055475.8** | Gpr18 | 0,71191288 | 8,061417842 | 17,60409652 | 2,72E-05 | 0,008870608 |
| **ENSMUST00000204498.1** | Gm44280 | 2,10551515 | 2,15921874 | 17,52632093 | 2,83E-05 | 0,009216716 |
| **ENSMUST00000024708.5** | Tnfrsf21 | 1,584030434 | 3,229964413 | 17,37891121 | 3,06E-05 | 0,009892078 |
| **ENSMUST00000033915.8** | Gpm6a | 2,631341363 | 1,334824432 | 17,16215365 | 3,44E-05 | 0,011019398 |
| **ENSMUST00000021028.4** | Itgb3 | 1,052570974 | 4,979349472 | 17,14037813 | 3,47E-05 | 0,011089499 |
| **ENSMUST00000043400.7** | Asprv1 | 2,309018389 | 1,791814166 | 17,09104218 | 3,67E-05 | 0,011591079 |
| **ENSMUST00000208023.1** | Slc35a2 | 2,259435725 | 1,824266541 | 17,00877201 | 3,72E-05 | 0,011727463 |
| **ENSMUST00000049787.2** | Lrrn4 | 1,633936313 | 3,07014468 | 16,99350495 | 3,75E-05 | 0,011783735 |
| **ENSMUST00000141708.1** | Plxdc1 | 2,818761143 | 1,055704752 | 16,95217177 | 3,83E-05 | 0,011950139 |
| **ENSMUST00000023105.4** | Endou | 0,787140835 | 6,800767368 | 16,81315244 | 4,13E-05 | 0,012664233 |
| **ENSMUST00000029875.3** | Pip4p2 | 1,332829542 | 3,840615409 | 16,72690092 | 4,32E-05 | 0,013139886 |
| **ENSMUST00000037099.8** | Clic4 | 0,643789061 | 8,882647916 | 16,65300859 | 4,49E-05 | 0,013574763 |
| **ENSMUST00000027752.14** | Lamc1 | 0,776612345 | 6,874243376 | 16,61247848 | 4,59E-05 | 0,013780172 |

| | | | | | | |
|---|---|---|---|---|---|---|
| **ENSMUST00000057561.8** | Wwc2 | 1,036929522 | 4,968346886 | 16,51719599 | 4,82E-05 | 0,014399082 |
| **ENSMUST00000045281.12** | Smg6 | 0,833354923 | 6,301554491 | 16,49661935 | 4,87E-05 | 0,014495414 |
| **ENSMUST00000091288.12** | Prnp | 1,667251143 | 2,917886515 | 16,39102658 | 5,16E-05 | 0,015235528 |
| **ENSMUST00000030949.3** | Tas1r3 | 1,222001363 | 4,147078432 | 16,33049231 | 5,32E-05 | 0,015628693 |
| **ENSMUST00000029078.8** | Car2 | 0,998261627 | 5,158079653 | 16,33032816 | 5,32E-05 | 0,015628693 |
| **ENSMUST00000210581.1** | Gm45353 | 1,605477738 | 3,045566074 | 16,09415714 | 6,03E-05 | 0,017487316 |
| **ENSMUST00000028179.14** | Fcnb | 1,946885528 | 2,322166731 | 16,07037795 | 6,25E-05 | 0,017998063 |
| **ENSMUST00000231927.1** | Rnaset2a | 0,915389624 | 5,625877892 | 16,01204011 | 6,29E-05 | 0,018078259 |
| **ENSMUST00000134982.7** | Gm14296 | 2,488825159 | 1,326195281 | 15,95959731 | 6,48E-05 | 0,018525725 |
| **ENSMUST00000234364.1** | 1810073O08Rik | 1,838884233 | 2,505484297 | 15,92594077 | 6,59E-05 | 0,018805713 |
| **ENSMUST00000178993.2** | Gm4617 | 0,621350107 | 9,037674858 | 15,88756742 | 6,72E-05 | 0,01915249 |
| **ENSMUST00000094897.4** | Dnaaf3 | 2,194825466 | 1,820541701 | 15,86551964 | 6,8E-05 | 0,019299718 |
| **ENSMUST00000132092.1** | 1110051M20Rik | 2,140659711 | 1,926134182 | 15,85193673 | 6,85E-05 | 0,019390703 |
| **ENSMUST00000199915.1** | Gm43737 | 0,959646607 | 5,304041997 | 15,84909436 | 6,86E-05 | 0,019390703 |
| **ENSMUST00000081649.9** | Ifitm2 | 1,675778866 | 2,842725493 | 15,79724437 | 7,05E-05 | 0,019819627 |
| **ENSMUST00000034983.6** | Atp1b3 | 0,616451805 | 9,068331327 | 15,70587445 | 7,41E-05 | 0,020660522 |
| **ENSMUST00000045487.3** | Rhou | 2,175048853 | 2,104316282 | 16,1961656 | 7,6E-05 | 0,021097427 |
| **ENSMUST00000090246.4** | Sgms2 | 2,375894668 | 1,583909448 | 15,85498177 | 7,62E-05 | 0,021131314 |
| **ENSMUST00000183583.7** | Ntng2 | 2,713081499 | 1,010883844 | 15,50033431 | 8,25E-05 | 0,022513142 |
| **ENSMUST00000022304.9** | Thrb | 2,465009772 | 1,332511772 | 15,48328466 | 8,32E-05 | 0,022673688 |
| **ENSMUST00000028593.10** | Prrg4 | 1,623305896 | 2,919716397 | 15,45181677 | 8,46E-05 | 0,02296651 |
| **ENSMUST00000160523.7** | Vamp1 | 1,381188592 | 3,541026352 | 15,41915374 | 8,61E-05 | 0,023322464 |
| **ENSMUST00000212475.1** | Gm7807 | 2,354005543 | 1,500624243 | 15,34909554 | 8,94E-05 | 0,024157467 |
| **ENSMUST00000170998.8** | Scn2b | 1,489581799 | 3,331547338 | 15,4954795 | 9,07E-05 | 0,024477231 |
| **ENSMUST00000144519.1** | Gps1 | 2,190738575 | 1,729844653 | 15,29938823 | 9,18E-05 | 0,024668887 |
| **ENSMUST00000060148.5** | Hivep1 | 0,627201462 | 8,653686716 | 15,21455299 | 9,6E-05 | 0,025648846 |
| **ENSMUST00000211473.1** | Gm9347 | 1,07377183 | 4,506600577 | 15,01030004 | 0,000106938 | 0,028313904 |
| **ENSMUST00000064187.11** | Thra | 1,46464195 | 3,245568694 | 14,93263581 | 0,000111431 | 0,029394289 |
| **ENSMUST00000056977.13** | Runx3 | 0,904869666 | 5,476641029 | 14,90572358 | 0,000113032 | 0,029712636 |
| **ENSMUST00000181211.1** | Gm17552 | 1,245442491 | 3,877798069 | 14,90655133 | 0,000113264 | 0,029712636 |
| **ENSMUST00000029158.3** | Aar2 | 2,141757873 | 1,767565526 | 14,88871784 | 0,000114056 | 0,029865364 |
| **ENSMUST00000094303.5** | Fcrl6 | 1,573454053 | 2,967723721 | 14,86673159 | 0,000115393 | 0,030049776 |
| **ENSMUST00000198190.4** | Rnf216 | 2,1680711 | 1,738065333 | 14,84510431 | 0,000116723 | 0,030212319 |
| **ENSMUST00000156997.7** | Dvl1 | 2,123140792 | 1,823795163 | 14,84503948 | 0,000116727 | 0,030212319 |
| **ENSMUST00000098799.4** | Ehd2 | 0,591317189 | 9,243669189 | 14,84331481 | 0,000116865 | 0,030212319 |
| **ENSMUST00000038053.13** | Lpp | 0,817505107 | 6,091819532 | 14,83191869 | 0,000117542 | 0,030253495 |
| **ENSMUST00000052457.14** | Mtss2 | 1,960211438 | 2,117583032 | 14,79164572 | 0,000120079 | 0,030693586 |
| **ENSMUST00000165774.7** | Gbp2 | 1,119464739 | 4,302242568 | 14,78943983 | 0,00012022 | 0,030693586 |
| **ENSMUST00000046206.4** | Rprd1a | 0,740099793 | 6,758436388 | 14,70886236 | 0,000125469 | 0,031858334 |
| **ENSMUST00000231568.1** | Ermard | 2,613916207 | 1,065845372 | 14,68957757 | 0,000126759 | 0,032014705 |
| **ENSMUST00000138611.7** | Ino80dos | 1,464950471 | 3,210541081 | 14,66560241 | 0,000128381 | 0,032309915 |
| **ENSMUST00000058524.2** | Zc3hav1l | 1,191379259 | 4,019837718 | 14,61840745 | 0,000131637 | 0,032978986 |
| **ENSMUST00000105818.7** | Kif17 | 2,290716976 | 1,498257089 | 14,61700719 | 0,000131734 | 0,032978986 |
| **ENSMUST00000190826.1** | Ly6m | 1,59735376 | 2,861653911 | 14,46330722 | 0,000142931 | 0,035554569 |
| **ENSMUST00000087867.5** | Uprt | 1,090461689 | 4,368119862 | 14,44429954 | 0,000144381 | 0,035767856 |
| **ENSMUST00000098513.5** | Plekhf1 | 1,96504943 | 2,049646196 | 14,4279329 | 0,000145641 | 0,035958993 |
| **ENSMUST00000049931.5** | Spn | 0,678072467 | 7,511650538 | 14,32202139 | 0,000154121 | 0,037892903 |
| **ENSMUST00000212276.1** | 2900026A02Rik | 2,519530348 | 1,106689538 | 14,31217287 | 0,000154876 | 0,037971555 |

| | | | | | | |
|---|---|---|---|---|---|---|
| **ENSMUST00000073109.11** | Ctdspl | 1,492178379 | 3,093966786 | 14,30662304 | 0,000155411 | 0,03803725 |
| **ENSMUST00000052678.8** | Flnb | 0,806014717 | 6,064554135 | 14,27437444 | 0,000158017 | 0,038586161 |
| **ENSMUST00000044492.9** | Akap9 | 0,760705726 | 6,422783521 | 14,2357382 | 0,000161295 | 0,039207817 |
| **ENSMUST00000120711.1** | Gm1848 | 0,760785311 | 6,414004482 | 14,20654002 | 0,000163817 | 0,039564816 |
| **ENSMUST00000178353.1** | Gm21992 | 1,498680905 | 3,067317817 | 14,20554851 | 0,000163903 | 0,039564816 |
| **ENSMUST00000026565.6** | Ifitm3 | 1,375575329 | 3,426334712 | 14,2246013 | 0,00016576 | 0,039750707 |
| **ENSMUST00000192833.1** | (None) lncRNA | 0,701764191 | 7,061849304 | 14,03283519 | 0,000179664 | 0,042583006 |
| **ENSMUST00000020350.14** | Lgr5 | 0,756865418 | 6,39686742 | 13,98456814 | 0,000184372 | 0,043462699 |
| **ENSMUST00000208484.1** | D7Bwg0826e | 1,940997967 | 2,034448473 | 13,976648 | 0,000185113 | 0,043462699 |
| **ENSMUST00000062211.3** | Gpat2 | 2,272488105 | 1,443188796 | 13,97570754 | 0,000185206 | 0,043462699 |
| **ENSMUST00000051803.7** | Aldh3b1 | 0,912361699 | 5,236743549 | 13,95996895 | 0,000186763 | 0,043684131 |
| **ENSMUST00000058154.14** | Tmtc3 | 1,475000559 | 3,093075609 | 13,9460486 | 0,000188151 | 0,043875928 |
| **ENSMUST00000205979.1** | Qpctl | 1,751379181 | 2,438628057 | 13,94372899 | 0,000188383 | 0,043875928 |
| **ENSMUST00000118960.1** | Car15 | 1,450243285 | 3,149629562 | 13,94249496 | 0,000188507 | 0,043875928 |
| **ENSMUST00000096338.4** | Gpr152 | 2,382783071 | 1,262397684 | 13,86321668 | 0,000196628 | 0,045468949 |
| **ENSMUST00000156839.1** | Ttpal | 2,198145953 | 1,565111647 | 13,85379841 | 0,000197616 | 0,045623333 |
| **ENSMUST00000059341.4** | Zc2hc1c | 1,568068621 | 2,864413198 | 13,8423194 | 0,000198883 | 0,045841677 |
| **ENSMUST00000005671.9** | Igf1r | 1,049573733 | 4,43546616 | 13,83779409 | 0,000199349 | 0,045874786 |
| **ENSMUST00000067458.6** | Sema5a | 3,639412476 | 4,416924321 | 34,1782988 | 0,000204773 | 0,04689568 |
| **ENSMUST00000112707.2** | Lrrc8b | 0,770240361 | 6,244318037 | 13,7875638 | 0,00020531 | 0,046928667 |
| **ENSMUST00000003561.9** | Phyhip | 2,096810671 | 1,721291854 | 13,77959715 | 0,000205576 | 0,046928667 |
| **ENSMUST00000135091.1** | Mtln | 1,405864455 | 3,247832623 | 13,75955962 | 0,000207781 | 0,047280357 |
| **ENSMUST00000053078.4** | Map10 | 1,113478994 | 4,180729456 | 13,75524674 | 0,000208258 | 0,047313452 |
| **ENSMUST00000231574.1** | Gm2792 | 0,953075215 | 4,927914037 | 13,72149338 | 0,000212034 | 0,048009217 |
| **ENSMUST00000053686.8** | Uck2 | 0,790459387 | 6,064749939 | 13,71885993 | 0,000212332 | 0,048009217 |
| **ENSMUST00000119848.7** | Eme2 | 0,944359867 | 4,972499349 | 13,70129894 | 0,000214326 | 0,04830686 |
| **ENSMUST00000119978.1** | Gm12671 | 0,638325412 | 7,902025085 | 13,66813018 | 0,000218145 | 0,049089967 |

| Downregulated in Dido1ΔE16 preB | | | | | | |
|---|---|---|---|---|---|---|
| **TranscriptID** | **Symbol** | **logFC** | **logCPM** | **F** | **PValue** | **FDR** |
| **ENSMUST00000087517.9** | Dido1 | -6,321931699 | 6,127988892 | 360,3444673 | 2,51E-80 | 3,57E-75 |
| **ENSMUST00000087543.4** | B3gatl4 | -3,661303587 | 4,557737119 | 102,7008546 | 3,92E-24 | 1,86E-20 |
| **ENSMUST00000060185.2** | Fndc9 | -1,953593451 | 6,973307638 | 83,07163692 | 7,94E-20 | 2,36E-16 |
| **ENSMUST00000038552.12** | Coro7 | -1,869896301 | 6,500700051 | 68,86100539 | 1,06E-16 | 2,32E-13 |
| **ENSMUST00000033056.4** | Pycard | -1,883482372 | 5,753463343 | 54,41246882 | 1,63E-13 | 2,79E-10 |
| **ENSMUST00000222616.1** | Gm19951 | -4,80983627 | 3,183063661 | 56,41428902 | 5,84E-12 | 8,07E-09 |
| **ENSMUST00000103510.1** | Ighv-26 | -1,56806827 | 6,193893519 | 45,3781159 | 1,63E-11 | 2,07E-08 |
| **ENSMUST00000154177.1** | Gm12678 | -1,655462674 | 5,889877298 | 45,31532015 | 1,68E-11 | 2,1E-08 |
| **ENSMUST00000135559.7** | Iffo1 | -4,15081223 | 2,8172429 | 45,28881682 | 1,7E-11 | 2,11E-08 |
| **ENSMUST00000112751.1** | Bcl2 | -2,270625247 | 4,416905344 | 44,30505812 | 5,2E-11 | 6,07E-08 |
| **ENSMUST00000232755.1** | Brwd1 | -1,388856764 | 6,525235209 | 40,566014 | 1,9E-10 | 2,07E-07 |
| **ENSMUST00000124462.2** | Arhgap27os3 | -1,889261018 | 4,729862382 | 37,95072549 | 7,32E-10 | 6,99E-07 |
| **ENSMUST00000025786.8** | Pacs1 | -1,692871768 | 5,242494037 | 37,59600881 | 8,9E-10 | 8,18E-07 |
| **ENSMUST00000103328.2** | Igkv10-96 | -1,692240036 | 5,090174833 | 35,70522519 | 2,3E-09 | 1,99E-06 |
| **ENSMUST00000048935.5** | DMrt3 | -2,045531931 | 4,255897118 | 34,85501199 | 3,59E-09 | 2,96E-06 |
| **ENSMUST00000102600.3** | Fndc5 | -1,837701248 | 4,602258186 | 34,4097588 | 4,47E-09 | 3,62E-06 |
| **ENSMUST00000159080.5** | Clec2i | -7,983593355 | 1,558483814 | 33,37174352 | 7,62E-09 | 6,03E-06 |

| | | | | | | |
|---|---|---|---|---|---|---|
| **ENSMUST00000139071.7** | Iffo1 | -7,890355654 | 1,463687259 | 31,36565171 | 2,14E-08 | 1,55E-05 |
| **ENSMUST00000138127.7** | Zfp318 | -4,442870304 | 1,956464378 | 29,51947886 | 5,54E-08 | 3,8E-05 |
| **ENSMUST00000144920.3** | Cmas | -2,003662183 | 3,954036653 | 29,30672633 | 6,31E-08 | 4,28E-05 |
| **ENSMUST00000139806.1** | Casz1 | -2,62217115 | 3,113146752 | 28,85298785 | 7,81E-08 | 5,23E-05 |
| **ENSMUST00000159745.1** | Pip4p1 | -1,716446429 | 4,448499287 | 28,85859076 | 7,82E-08 | 5,23E-05 |
| **ENSMUST00000108809.7** | Trim11 | -1,328841978 | 6,363271883 | 31,2363446 | 8,09E-08 | 5,36E-05 |
| **ENSMUST00000076957.6** | Zdhhc8 | -1,179685888 | 6,382338265 | 28,64868403 | 8,68E-08 | 5,72E-05 |
| **ENSMUST00000130486.1** | Gm15675 | -1,339694373 | 5,647834982 | 28,47516073 | 9,58E-08 | 6,23E-05 |
| **ENSMUST00000117906.1** | Gm14127 | -1,420073761 | 5,320221421 | 28,24872281 | 1,07E-07 | 6,91E-05 |
| **ENSMUST00000103507.1** | Ighv1-22 | -1,800719921 | 4,202435221 | 27,47968675 | 1,59E-07 | 9,84E-05 |
| **ENSMUST00000103495.2** | Ighv10-3 | -1,495800386 | 4,954604049 | 27,25163869 | 1,79E-07 | 0,000110187 |
| **ENSMUST00000103544.2** | Ighv1-75 | -1,464349621 | 5,054374643 | 27,18513212 | 1,85E-07 | 0,000113064 |
| **ENSMUST00000192591.1** | Ighv8-8 | -1,398220873 | 5,268371238 | 26,98035366 | 2,06E-07 | 0,000123576 |
| **ENSMUST00000201831.3** | Gm20559 | -1,182468775 | 6,205358653 | 26,98167247 | 2,22E-07 | 0,000131625 |
| **ENSMUST00000052124.8** | Nlrc4 | -1,870336605 | 4,02341972 | 26,68329429 | 2,4E-07 | 0,00014055 |
| **ENSMUST00000103526.2** | Ighv1-55 | -1,310757938 | 5,559107453 | 26,45566013 | 2,7E-07 | 0,000156834 |
| **ENSMUST00000026408.6** | Gdf11 | -1,261248622 | 5,657364824 | 25,54015348 | 4,34E-07 | 0,000241657 |
| **ENSMUST00000001327.10** | Itgb7 | -2,61163875 | 3,107993862 | 26,35748488 | 4,57E-07 | 0,000252715 |
| **ENSMUST00000219633.1** | Icosl | -7,525804932 | 1,161075383 | 25,31362351 | 4,9E-07 | 0,000267191 |
| **ENSMUST00000057725.9** | Samhd1 | -0,792015264 | 9,539454369 | 24,98671412 | 5,77E-07 | 0,000310391 |
| **ENSMUST00000171330.6** | Slamf6 | -1,13563657 | 6,058165089 | 24,0025037 | 9,62E-07 | 0,000493113 |
| **ENSMUST00000124485.7** | Fam129c | -1,066469298 | 6,676448923 | 24,608023 | 9,92E-07 | 0,000504511 |
| **ENSMUST00000030417.9** | Cdc42 | -1,590191046 | 4,399796876 | 23,90182956 | 1,21E-06 | 0,000602967 |
| **ENSMUST00000035218.8** | Nckipsd | -1,283554478 | 5,317747107 | 23,40961893 | 1,31E-06 | 0,000645474 |
| **ENSMUST00000136582.1** | Dpm1 | -1,543326311 | 4,409283137 | 23,36333687 | 1,34E-06 | 0,000656648 |
| **ENSMUST00000103515.1** | Ighv1-39 | -1,547760676 | 4,378250288 | 23,00184139 | 1,62E-06 | 0,000781701 |
| **ENSMUST00000029569.8** | Slc35a3 | -1,230029896 | 5,472174724 | 22,83947362 | 1,76E-06 | 0,000841061 |
| **ENSMUST00000224965.1** | Blk | -1,250333269 | 5,365670422 | 22,82565724 | 1,78E-06 | 0,000845437 |
| **ENSMUST00000108846.1** | Galnt10 | -2,251345495 | 3,090909269 | 22,57263474 | 2,03E-06 | 0,000942717 |
| **ENSMUST00000103386.2** | Igkv6-23 | -1,356995082 | 4,870950986 | 22,1892887 | 2,47E-06 | 0,001128871 |
| **ENSMUST00000127842.7** | Ctsh | -3,440887594 | 1,979518078 | 22,14112208 | 2,54E-06 | 0,001150331 |
| **ENSMUST00000177669.1** | Pfn1 | -1,038391608 | 6,305071193 | 22,02354857 | 2,7E-06 | 0,001218565 |
| **ENSMUST00000054279.14** | Sp100 | -1,679650835 | 4,002008163 | 21,97562283 | 2,76E-06 | 0,001241119 |
| **ENSMUST00000028755.7** | Ehd4 | -0,852689493 | 7,956604028 | 21,84851254 | 2,95E-06 | 0,001313675 |
| **ENSMUST00000094646.5** | Vps4b | -0,9528883 | 6,769731742 | 21,2618847 | 4,01E-06 | 0,001683743 |
| **ENSMUST00000110031.3** | Auh | -1,699163386 | 3,82382454 | 20,55017493 | 5,81E-06 | 0,002357878 |
| **ENSMUST00000229230.1** | Gm18724 | -1,248267124 | 5,065038697 | 20,40789211 | 6,26E-06 | 0,002511213 |
| **ENSMUST00000098486.3** | Bcl2a1d | -3,109744396 | 2,036965161 | 20,12783914 | 7,27E-06 | 0,002883393 |
| **ENSMUST00000119311.7** | Auh | -1,774167059 | 3,641515153 | 20,09724599 | 7,36E-06 | 0,002912898 |
| **ENSMUST00000028205.9** | BC005624 | -1,062442735 | 5,875275434 | 19,95054576 | 7,95E-06 | 0,003119156 |
| **ENSMUST00000124563.7** | Napa | -2,896490967 | 2,171873297 | 19,93717851 | 8,02E-06 | 0,003136808 |
| **ENSMUST00000191475.1** | Gm8369 | -3,285587564 | 1,914526522 | 19,76182214 | 8,77E-06 | 0,003386816 |
| **ENSMUST00000105520.7** | Enpp1 | -2,7745023 | 2,294940484 | 19,70209371 | 9,05E-06 | 0,003475495 |
| **ENSMUST00000236643.1** | H2-Ab1 | -1,809995864 | 3,510553429 | 19,50810809 | 1,01E-05 | 0,003834755 |
| **ENSMUST00000028928.7** | Gzf1 | -1,002841441 | 6,123570802 | 19,40183321 | 1,06E-05 | 0,004012909 |
| **ENSMUST00000089688.5** | Mmp14 | -3,049193137 | 2,042926237 | 19,38315756 | 1,07E-05 | 0,004041594 |
| **ENSMUST00000032288.5** | Klra1 | -2,381443479 | 2,679955902 | 19,16790685 | 1,2E-05 | 0,004488191 |
| **ENSMUST00000200318.1** | Reln | -1,801471893 | 3,508443606 | 19,06073033 | 1,27E-05 | 0,004699707 |

| | | | | | | |
|---|---|---|---|---|---|---|
| **ENSMUST00000194738.5** | Igha | -2,28159457 | 2,761208115 | 19,01513182 | 1,3E-05 | 0,004802556 |
| **ENSMUST00000103521.2** | Ighv1-50 | -1,339319172 | 4,520063 | 18,8886975 | 1,39E-05 | 0,005037705 |
| **ENSMUST00000103321.2** | Igkv1-110 | -1,244154495 | 4,83149238 | 18,69219071 | 1,54E-05 | 0,005531082 |
| **ENSMUST00000033611.4** | Xkrx | -1,877057128 | 4,45365182 | 22,18038806 | 1,65E-05 | 0,005898442 |
| **ENSMUST00000149244.7** | Lrmp | -1,320774437 | 4,50840102 | 18,5478536 | 1,66E-05 | 0,005919815 |
| **ENSMUST00000059644.12** | Rbm33 | -0,933005737 | 6,391954558 | 18,50911402 | 1,69E-05 | 0,006016918 |
| **ENSMUST00000103330.1** | Igkv10-94 | -1,581022172 | 3,863354835 | 18,47096412 | 1,73E-05 | 0,006113882 |
| **ENSMUST00000171415.7** | Ndufb8 | -1,546232139 | 3,937086795 | 18,44760915 | 1,75E-05 | 0,006158626 |
| **ENSMUST00000190715.6** | Cep70 | -2,040272535 | 2,979809197 | 17,91385147 | 2,31E-05 | 0,007766293 |
| **ENSMUST00000185851.1** | Ms4a1 | -1,574366466 | 3,801611177 | 17,7073616 | 2,58E-05 | 0,008535729 |
| **ENSMUST00000189288.1** | F730311O21 | -1,301374978 | 4,811238435 | 18,38186761 | 2,59E-05 | 0,008572595 |
| **ENSMUST00000107802.7** | Trim59 | -0,855661493 | 6,779539165 | 17,35938084 | 3,09E-05 | 0,009971639 |
| **ENSMUST00000049872.8** | Gpr183 | -1,171565745 | 5,567322821 | 18,39201523 | 3,19E-05 | 0,010252892 |
| **ENSMUST00000123453.1** | Gmip | -2,080640912 | 2,83817036 | 17,09162441 | 3,56E-05 | 0,011327043 |
| **ENSMUST00000074259.14** | Nrm | -0,765176205 | 7,74999753 | 17,0492499 | 3,64E-05 | 0,011556825 |
| **ENSMUST00000077605.11** | Eif4a2 | -0,857011307 | 6,70463286 | 17,09189787 | 3,65E-05 | 0,011557296 |
| **ENSMUST00000088785.5** | Zfp566 | -1,521634845 | 3,830695557 | 16,99129999 | 3,76E-05 | 0,011783735 |
| **ENSMUST00000025266.5** | Lta | -2,498746182 | 2,32040539 | 16,90669591 | 3,96E-05 | 0,012262889 |
| **ENSMUST00000089497.6** | Lsy1 | -1,005583807 | 5,690951661 | 16,88235191 | 3,98E-05 | 0,012290273 |
| **ENSMUST00000034466.9** | Gnpat | -1,1617632 | 4,901555007 | 16,86739962 | 4,01E-05 | 0,01236066 |
| **ENSMUST00000014597.4** | Blk | -0,648216171 | 9,463687199 | 16,77374043 | 4,21E-05 | 0,012867344 |
| **ENSMUST00000103369.1** | Igkv12_41 | -1,843511231 | 3,176795449 | 16,64238842 | 4,51E-05 | 0,013622056 |
| **ENSMUST00000168846.2** | Prkag1 | -0,95133699 | 5,941714353 | 16,53153497 | 4,79E-05 | 0,01435078 |
| **ENSMUST00000063140.14** | Hcrtr2 | -4,031617164 | 1,218501868 | 16,50007809 | 4,87E-05 | 0,014495414 |
| **ENSMUST00000049295.14** | Ehbp1l1 | -0,986878979 | 5,72915136 | 16,49113643 | 4,89E-05 | 0,014507106 |
| **ENSMUST00000106357.7** | Ypel3 | -1,209641194 | 4,602325694 | 16,37671719 | 5,19E-05 | 0,015313957 |
| **ENSMUST00000027266.3** | Ormdl1 | -0,9487111 | 5,913806377 | 16,29367741 | 5,43E-05 | 0,015901113 |
| **ENSMUST00000030964.5** | CD38 | -0,651117231 | 9,203340782 | 16,23521185 | 5,6E-05 | 0,016365734 |
| **ENSMUST00000103492.1** | Ighv10_1 | -1,084158731 | 5,162109654 | 16,20811053 | 5,68E-05 | 0,016567508 |
| **ENSMUST00000096243.6** | B3gat3 | -1,085825454 | 5,11984947 | 16,08608819 | 6,05E-05 | 0,017500942 |
| **ENSMUST00000153442.7** | Hnrnpdl | -1,096473772 | 5,059817253 | 16,08560917 | 6,06E-05 | 0,017500942 |
| **ENSMUST00000074733.10** | Sept11 | -0,708824 | 8,200756884 | 16,06479354 | 6,12E-05 | 0,017652724 |
| **ENSMUST00000103493.2** | Ighv1_4 | -1,602674675 | 3,567852863 | 15,9900552 | 6,37E-05 | 0,01825261 |
| **ENSMUST00000015460.4** | Slamf1 | -2,286564693 | 2,455558045 | 15,87475288 | 6,78E-05 | 0,019273272 |
| **ENSMUST00000103534.1** | Ighv1-63 | -1,756876208 | 3,266879808 | 15,83304593 | 6,93E-05 | 0,019498865 |
| **ENSMUST00000120267.8** | Atg16l2 | -0,927330632 | 5,944811642 | 15,76609552 | 7,17E-05 | 0,020061185 |
| **ENSMUST00000161705.2** | Mcoln1 | -1,22439226 | 4,453059577 | 15,67899419 | 7,51E-05 | 0,020896092 |
| **ENSMUST00000173103.1** | H2-Ab1 | -1,822872128 | 3,052427033 | 15,58433112 | 7,92E-05 | 0,021770549 |
| **ENSMUST00000070720.7** | Sorcs2 | -1,316907882 | 4,181814742 | 15,51468541 | 8,19E-05 | 0,022385759 |
| **ENSMUST00000219038.1** | Icosl | -3,030945151 | 1,624124969 | 15,22695931 | 9,53E-05 | 0,025528804 |
| **ENSMUST00000164035.7** | Arid3b | -1,747583514 | 3,159753895 | 15,22848471 | 9,53E-05 | 0,025528804 |
| **ENSMUST00000140896.1** | Coro1a | -1,402439401 | 3,905593784 | 15,2313424 | 9,66E-05 | 0,025761121 |
| **ENSMUST00000103548.2** | Ighv1_81 | -1,022871062 | 5,272466495 | 15,10425623 | 0,000101745 | 0,027039536 |
| **ENSMUST00000037913.8** | Rmi2 | -0,853572767 | 6,324823147 | 15,08240218 | 0,000106569 | 0,028268881 |
| **ENSMUST00000195325.1** | Ighv1_2 | -1,506154793 | 3,650215269 | 14,99575642 | 0,000107765 | 0,028480012 |
| **ENSMUST00000123602.1** | Dctn5 | -1,632783615 | 3,336595336 | 14,90422859 | 0,000113244 | 0,029712636 |
| **ENSMUST00000200926.1** | Nabp2l1 | -2,27234845 | 2,326272308 | 14,88002122 | 0,000114583 | 0,029948313 |
| **ENSMUST00000117892.1** | Slc48a1 | -0,771435071 | 6,987704437 | 14,85486262 | 0,000116121 | 0,030184274 |

| | | | | | | |
|---|---|---|---|---|---|---|
| **ENSMUST00000035842.6** | Rassf1 | -0,654480122 | 8,580568303 | 14,80443381 | 0,000119268 | 0,030611202 |
| **ENSMUST00000204885.1** | Hnrnpa2b1 | -0,994640097 | 5,339542027 | 14,72942994 | 0,000124107 | 0,031625416 |
| **ENSMUST00000113481.8** | Zfp318 | -0,799788028 | 6,608211956 | 14,71730276 | 0,000125115 | 0,031825347 |
| **ENSMUST00000103384.1** | Igkv8_24 | -1,313846169 | 4,059313433 | 14,53626051 | 0,000137502 | 0,034362495 |
| **ENSMUST00000207327.1** | Fcer2a | -3,283156331 | 1,372850445 | 14,44130764 | 0,000145658 | 0,035958993 |
| **ENSMUST00000072729.9** | Ms4a4c | -2,331771874 | 2,251738081 | 14,33233054 | 0,000158196 | 0,038586161 |
| **ENSMUST00000119728.1** | Npm3-ps1 | -0,924176794 | 5,638522107 | 14,15437794 | 0,000168422 | 0,040274585 |
| **ENSMUST00000001455.12** | Mef2d | -0,595348928 | 9,43343979 | 14,15338483 | 0,000168511 | 0,040274585 |
| **ENSMUST00000172057.7** | Ralgps2 | -0,67314841 | 8,049282892 | 14,1192891 | 0,000171592 | 0,040942466 |
| **ENSMUST00000186927.1** | Ly6e | -1,186612293 | 4,339905695 | 13,99950832 | 0,000182937 | 0,04321498 |
| **ENSMUST00000192106.1** | Gm8146 | -1,425706995 | 3,686588118 | 13,98002222 | 0,000184781 | 0,043462699 |
| **ENSMUST00000150208.7** | Napa | -1,487700985 | 3,537511902 | 13,9375477 | 0,000190788 | 0,044299427 |
| **ENSMUST00000143378.7** | RIKEN cDNA 4930426D05 | -1,063285171 | 5,357939252 | 16,77561341 | 0,0000422 | 0,012867344 |
| **ENSMUST00000128652.1** | RIKEN cDNA 4930597A21 | -1,00947143 | 5,618464323 | 16,30293479 | 0,0000572 | 0,016656537 |
