## Supplementary material for "The chromatin reader Dido3 is a regulator of the gene network that controls B cell differentiation": Suppl3mentary Table 3

### Supplementary Table 3

Differentially expresed Ighv (variable) regions in pre B cells. RNA-seq data.

| | logFC | logCPM | LR | PValue | FDR |
|---|---|---|---|---|---|
| ENSMUSG00000076613_Ighg2b | 5,177375763 | 11,0114599 | 201,453585 | 1,01E-45 | 2,80E-43 |
| ENSMUSG00000094028_Ighd4-1 | 3,435788362 | 11,6479589 | 114,786492 | 8,76E-27 | 1,22E-24 |
| ENSMUSG00000094057_Ighd2-7 | 4,13279968 | 8,67285432 | 92,4153468 | 7,03E-22 | 6,51E-20 |
| ENSMUSG00000095897_Ighd2-5 | 2,705912053 | 9,02573405 | 60,6345883 | 6,87E-15 | 4,78E-13 |
| ENSMUSG00000095444_Ighd2-4 | 2,203923512 | 9,67438993 | 54,5985616 | 1,48E-13 | 8,22E-12 |
| ENSMUSG00000076630_Ighd1-1 | 1,793919186 | 11,253159 | 30,6397651 | 3,11E-08 | 1,44E-06 |
| ENSMUSG00000096568_Ighd2-3 | 1,798574118 | 9,51005007 | 29,510394 | 5,56E-08 | 2,21E-06 |
| ENSMUSG00000096150_Ighv1-85 | 2,001683257 | 8,37291067 | 21,7637665 | 3,08E-06 | 0,00010716 |
| ENSMUSG00000093955_Ighv1-34 | -1,683806396 | 8,63910452 | 20,0707275 | 7,46E-06 | 0,00023052 |
| ENSMUSG00000076633_Ighv5-2 | 1,149102373 | 10,7760075 | 18,7699476 | 1,47E-05 | 0,00040997 |
| ENSMUSG00000096464_Ighv2-2 | 1,196086747 | 11,7479314 | 17,8036236 | 2,45E-05 | 0,00061897 |
| ENSMUSG00000094117_Igkv3-12 | 1,177994346 | 11,2784643 | 16,1877401 | 5,74E-05 | 0,00132894 |
| ENSMUSG00000095416_Ighv1-12 | -1,180049493 | 9,75657498 | 15,9939 | 6,35E-05 | 0,00135893 |
| ENSMUSG00000103439_Gm7019 | 1,111729657 | 12,0007745 | 15,0978134 | 0,00010208 | 0,00202705 |
| ENSMUSG00000095130_Ighv1-39 | -1,073056122 | 10,0470355 | 14,7446491 | 0,0001231 | 0,00228139 |
| ENSMUSG00000105630_Gm42543 | 1,958377857 | 7,62244402 | 13,8997266 | 0,00019283 | 0,00331427 |
| ENSMUSG00000094051_Ighv1-36 | -1,729868909 | 8,40680503 | 13,7040139 | 0,000214 | 0,00331427 |
| ENSMUSG00000076620_Ighj2 | 1,173126641 | 9,80407887 | 13,6987894 | 0,00021459 | 0,00331427 |
| ENSMUSG00000094561_Ighv1-22 | -0,968802278 | 10,0309777 | 11,9406639 | 0,00054922 | 0,00803594 |
| ENSMUSG00000076531_Igkv4-92 | 0,947768886 | 9,64418052 | 11,4133208 | 0,00072919 | 0,01013579 |
| ENSMUSG00000094940_Ighv1-84 | -1,569747231 | 7,86347906 | 11,0600177 | 0,00088209 | 0,01167723 |
| ENSMUSG00000076598_Igkv3-7 | 0,974039758 | 11,8250008 | 10,7807443 | 0,00102561 | 0,01296004 |
| ENSMUSG00000096074_Ighv1-72 | 1,078454326 | 9,63555533 | 10,0677939 | 0,00150884 | 0,01823723 |
| ENSMUSG00000095866_Ighv2-4 | 0,955057938 | 9,2066517 | 9,66627868 | 0,00187681 | 0,02173973 |
| ENSMUSG00000096649_Ighv1-31 | -1,603278868 | 7,74617826 | 9,53362458 | 0,00201741 | 0,02243363 |
| ENSMUSG00000094652_Ighv1-42 | -0,975189499 | 9,26275221 | 9,32514755 | 0,0022603 | 0,02379573 |
| ENSMUSG00000076695_Ighv1-18 | -0,92584829 | 10,3922373 | 9,2754911 | 0,0023224 | 0,02379573 |
| ENSMUSG00000095863_Ighv1-67 | -1,144229293 | 8,51612379 | 9,21783174 | 0,00239669 | 0,02379573 |
| ENSMUSG00000093896_Ighv1-76 | -0,834787482 | 10,7236013 | 8,36705245 | 0,00382085 | 0,03662744 |
| ENSMUSG00000094546_Ighv1-26 | -0,911194794 | 11,9062431 | 8,2095574 | 0,00416703 | 0,03857043 |
| ENSMUSG00000106124_Gm42539 | 1,869388279 | 6,46688357 | 8,15214777 | 0,00430102 | 0,03857043 |
| ENSMUSG00000103537_Gm37838 | 1,567022347 | 6,59522279 | 7,97100381 | 0,00475326 | 0,04129392 |
| ENSMUSG00000094694_Ighv1-9 | -0,759644961 | 11,0040389 | 7,40626887 | 0,0064997 | 0,05475503 |
| ENSMUSG00000094951_Ighv5-6 | 0,726667156 | 11,0872131 | 7,30006217 | 0,00689522 | 0,05490461 |
| ENSMUSG00000095700_Ighv10-3 | -0,783609818 | 10,5656279 | 7,29557865 | 0,00691245 | 0,05490461 |
| ENSMUSG00000095612_Ighv5-4 | 0,74439873 | 11,5734534 | 7,11169352 | 0,00765827 | 0,05913888 |
| ENSMUSG00000000001_Gm30996 | 2,102964555 | 4,81116565 | 6,75350437 | 0,00935637 | 0,07029924 |
| ENSMUSG00000103297_Ighv3-2 | -1,472817484 | 7,33039719 | 6,6933679 | 0,00967722 | 0,07079652 |
| ENSMUSG00000104760_Igkv3-8 | 1,658945266 | 5,00512581 | 6,47040267 | 0,01096857 | 0,0781862 |
| ENSMUSG00000102942_Ighv1-33 | -2,126420546 | 3,91093715 | 6,39753437 | 0,0114279 | 0,07942388 |
| ENSMUSG00000094533_Ighv11-1 | -1,669326819 | 5,8759403 | 6,31963143 | 0,01194084 | 0,08005876 |
| ENSMUSG00000076596_Igkv3-10 | 0,673171372 | 10,4226837 | 6,29685931 | 0,01209521 | 0,08005876 |
| ENSMUSG00000106098_Igkv14-118-2 | 1,495531483 | 5,31228066 | 6,05846689 | 0,01383978 | 0,08947577 |
| ENSMUSG00000076501_Igkv2-137 | 0,646926589 | 10,8553969 | 5,99636998 | 0,01433534 | 0,08980706 |
| ENSMUSG00000095197_Ighv1-59 | -0,721533304 | 9,91442933 | 5,94771485 | 0,01473638 | 0,08980706 |
| ENSMUSG00000102654_Ighv1-21 | -1,201134567 | 7,45384167 | 5,93297185 | 0,01486016 | 0,08980706 |
| ENSMUSG00000094087_Ighv1-61 | -0,759887613 | 9,16421927 | 5,71295115 | 0,01684021 | 0,09960803 |

| | | | | | |
|---|---|---|---|---|---|
| ENSMUSG00000094164_Ighv2-3 | 0,685587426 | 9,60526463 | 5,34609232 | 0,02076877 | 0,12028578 |
| ENSMUSG00000103168_Gm30948 | 2,033469731 | 3,00730278 | 5,22282817 | 0,02229227 | 0,12408686 |
| ENSMUSG00000106601_Ighv1-70 | -1,381132688 | 5,69079483 | 5,22083902 | 0,02231778 | 0,12408686 |
| ENSMUSG00000096594_Igkv8-19 | 0,632206719 | 11,1831626 | 5,11593087 | 0,02370714 | 0,12805691 |
| ENSMUSG00000095170_Ighv8-11 | -1,36949941 | 6,25424398 | 5,09802581 | 0,02395309 | 0,12805691 |
| ENSMUSG00000094075_Ighv1-80 | -0,620116103 | 10,0480177 | 4,95710873 | 0,02598362 | 0,13629142 |
| ENSMUSG00000095981_Ighv10-1 | -0,588829595 | 10,554939 | 4,81117145 | 0,02827582 | 0,1455681 |
| ENSMUSG00000102888_Ighv1-11 | -0,92138606 | 8,27124633 | 4,77202889 | 0,02892572 | 0,14620636 |
| ENSMUSG00000095442_Ighv1-4 | -0,663938949 | 9,15982116 | 4,71780225 | 0,02985187 | 0,14819323 |
| ENSMUSG00000076589_Igkv8-18 | 0,947540018 | 8,36825993 | 4,62057115 | 0,03159072 | 0,15407403 |
| ENSMUSG00000102952_Ighv1-25 | -1,307576367 | 5,74547552 | 4,29499462 | 0,03822472 | 0,18321502 |
| ENSMUSG00000095771_Igkv14-111 | 0,533274469 | 10,2864999 | 4,17979124 | 0,04090874 | 0,19275643 |
| ENSMUSG00000105462_Igkv4-77 | 0,531311477 | 9,93846955 | 4,13771285 | 0,0419378 | 0,19431179 |
| ENSMUSG00000076676_Ighv12-3 | -0,687180345 | 9,13269173 | 4,10008879 | 0,04288096 | 0,19542471 |
| ENSMUSG00000094198_Ighv1-50 | -0,585001217 | 10,1161914 | 4,01208031 | 0,04517538 | 0,2005046 |
| ENSMUSG00000094689_Ighv1-81 | -0,584217099 | 10,9978384 | 3,99755463 | 0,04556633 | 0,2005046 |
| ENSMUSG00000076621_Ighj1 | 0,884264822 | 14,4042261 | 3,97577047 | 0,04615933 | 0,2005046 |
| ENSMUSG00000095589_Ighv1-55 | -0,555040787 | 10,9645996 | 3,94440051 | 0,04702761 | 0,20113346 |
| ENSMUSG00000094552_Ighd3-2 | 0,773590698 | 8,43986094 | 3,91042688 | 0,04798741 | 0,20212877 |
| ENSMUSG00000096672_Ighv1-63 | -0,763362468 | 8,82568421 | 3,76300446 | 0,05239835 | 0,21741404 |
| ENSMUSG00000106599_Ighv1-73 | -1,398410132 | 4,48734055 | 3,71656662 | 0,05387505 | 0,22025386 |
| ENSMUSG00000102301_Ighv8-2 | -0,936167637 | 7,69514926 | 3,5994213 | 0,05779969 | 0,23287411 |
| ENSMUSG00000104213_Ighd | -0,659177666 | 9,88175883 | 3,47103318 | 0,06245229 | 0,24647816 |
| ENSMUSG00000104769_Igkv8-34 | 0,941357387 | 7,02804306 | 3,443371 | 0,063506 | 0,24647816 |
| ENSMUSG00000104452_Ighv8-8 | -0,527043772 | 11,3098304 | 3,43480628 | 0,06383607 | 0,24647816 |
| ENSMUSG00000095429_Ighv5-12 | 0,49803958 | 10,6063664 | 3,34159822 | 0,06754897 | 0,25724128 |
| ENSMUSG00000076680_Ighv6-6 | -0,527580202 | 9,7000838 | 3,21736697 | 0,07286074 | 0,27067816 |
| ENSMUSG00000073028_Igkv4-71 | -0,679573852 | 8,77448633 | 3,18170461 | 0,07446701 | 0,27067816 |
| ENSMUSG00000076581_Igkv8-26 | 1,030033169 | 6,3041543 | 3,16804027 | 0,0750925 | 0,27067816 |
| ENSMUSG00000096250_Ighd2-6 | -0,587084644 | 10,4542299 | 3,10255832 | 0,07816936 | 0,27067816 |
| ENSMUSG00000076606_Igkj3 | 0,640924259 | 11,5116641 | 3,09446147 | 0,07855914 | 0,27067816 |
| ENSMUSG00000095127_Ighv1-82 | -0,504245407 | 11,5938878 | 3,08994271 | 0,07877758 | 0,27067816 |
| ENSMUSG00000102678_Ighv1-21-1 | -1,004043452 | 5,873329 | 3,08886053 | 0,07882999 | 0,27067816 |
| ENSMUSG00000094787_Ighv1-54 | -0,5071661 | 9,60429213 | 3,08810393 | 0,07886666 | 0,27067816 |
| ENSMUSG00000094420_Igkv10-96 | -0,453739549 | 10,5607213 | 3,06213137 | 0,08013649 | 0,27168225 |
| ENSMUSG00000095571_Ighv5-17 | 0,463357616 | 10,6474763 | 2,91762159 | 0,08761692 | 0,29001116 |
| ENSMUSG00000094262_Igkv4-62 | 0,874528166 | 7,16048478 | 2,9054264 | 0,0882819 | 0,29001116 |
| ENSMUSG00000106403_Igkv20-101-2 | -1,082974466 | 6,07234901 | 2,88025985 | 0,08967155 | 0,29001116 |
| ENSMUSG00000095761_Ighv1-20 | -1,055689119 | 7,01745686 | 2,8776134 | 0,08981905 | 0,29001116 |
| ENSMUSG00000103203_Gm37327 | 1,326136828 | 5,73368861 | 2,85504232 | 0,09108784 | 0,29001116 |
| ENSMUSG00000103271_Ighv4-2 | -0,806116273 | 7,60655197 | 2,80778994 | 0,09380753 | 0,29001116 |
| ENSMUSG00000096078_Ighv1-62-2 | 1,001763931 | 7,45787547 | 2,80751714 | 0,09382348 | 0,29001116 |
| ENSMUSG00000095794_Igkv6-17 | 0,429882915 | 10,4922438 | 2,79456216 | 0,0945846 | 0,29001116 |
| ENSMUSG00000094134_Ighv5-15 | 0,492452966 | 9,2415781 | 2,78869158 | 0,09493171 | 0,29001116 |
| ENSMUSG00000076731_Ighv8-12 | -0,505490423 | 10,6615352 | 2,76956572 | 0,09607222 | 0,2903052 |
| ENSMUSG00000076666_Ighv14-4 | -0,439885409 | 10,9026793 | 2,64081998 | 0,10414978 | 0,31132944 |
| ENSMUSG00000096844_Igkv6-14 | 0,598552395 | 8,88294692 | 2,58305221 | 0,10801321 | 0,31944331 |
| ENSMUSG00000095630_Igkv6-23 | -0,436742903 | 9,76588655 | 2,49506662 | 0,11420354 | 0,33403597 |
| ENSMUSG00000094502_Ighv1-69 | -0,406074466 | 10,2378681 | 2,47934131 | 0,11535055 | 0,33403597 |
| ENSMUSG00000103873_Ighv1-38 | -0,847369758 | 7,1242255 | 2,44680215 | 0,11776457 | 0,33688302 |
| ENSMUSG00000104103_Gm9517 | 0,568057076 | 8,59524348 | 2,43363375 | 0,11875732 | 0,33688302 |
| ENSMUSG00000093838_Ighv3-1 | -0,524645867 | 8,89825088 | 2,36655136 | 0,12396063 | 0,34809146 |
| ENSMUSG00000094094_Igkv5-45 | 0,397046687 | 10,0367108 | 2,30828563 | 0,12868591 | 0,35774684 |

| | | | | | |
|---|---|---|---|---|---|
| ENSMUSG00000094345_Igkv14-126 | -0,501278722 | 9,03501953 | 2,27166813 | 0,13175802 | 0,36266068 |
| ENSMUSG00000096499_Ighv1-5 | -0,41639089 | 11,0010588 | 2,2420177 | 0,13430563 | 0,36604867 |
| ENSMUSG00000105906_Iglc1 | 0,697952416 | 15,6446864 | 2,2132972 | 0,13682596 | 0,36929724 |
| ENSMUSG00000103989_Ighv5-21 | 1,185740316 | 3,696691 | 2,18464116 | 0,13939355 | 0,37260968 |
| ENSMUSG00000095335_Igkv3-5 | 0,404687686 | 9,63444883 | 2,11418872 | 0,14593937 | 0,38639186 |
| ENSMUSG00000095285_Ighv5-9 | 0,490247096 | 8,82247985 | 2,09121541 | 0,14814817 | 0,3869217 |
| ENSMUSG00000076672_Ighv3-6 | -0,439935206 | 11,6608676 | 2,08324695 | 0,1489231 | 0,3869217 |
| ENSMUSG00000095204_Ighv1-52 | -0,383154594 | 9,90330242 | 2,05735825 | 0,15147257 | 0,38990163 |
| ENSMUSG00000105955_Igkv12-40 | -0,831343442 | 7,37205575 | 2,01372719 | 0,15588195 | 0,39520584 |
| ENSMUSG00000094993_Igkv4-51 | 0,54121124 | 8,50420195 | 2,00809557 | 0,15646162 | 0,39520584 |
| ENSMUSG00000076543_Igkv4-74 | -0,517388412 | 8,59620808 | 1,99520211 | 0,15779801 | 0,39520584 |
| ENSMUSG00000105757_Ighv1-83 | -0,840099243 | 5,46038142 | 1,97360712 | 0,16006551 | 0,39730545 |
| ENSMUSG00000096498_Ighv2-5 | 0,376127475 | 9,99479459 | 1,93448397 | 0,16426871 | 0,4041301 |
| ENSMUSG00000106668_Iglj1 | 0,65977613 | 16,2949127 | 1,8530564 | 0,17342833 | 0,42292173 |
| ENSMUSG00000104712_Igkv4-75 | -0,776275593 | 6,01635123 | 1,81471057 | 0,17794418 | 0,43016072 |
| ENSMUSG00000076934_Iglv1 | 0,621829349 | 15,5288016 | 1,79752389 | 0,18001213 | 0,43140839 |
| ENSMUSG00000103254_Ighv1-15 | -0,420562261 | 9,70403143 | 1,6777382 | 0,19522526 | 0,46370708 |
| ENSMUSG00000094433_Igkv5-43 | 0,335543925 | 10,727831 | 1,65992101 | 0,19761399 | 0,46370708 |
| ENSMUSG00000076578_Igkv6-29 | -0,702788094 | 7,42870741 | 1,65342567 | 0,19849332 | 0,46370708 |
| ENSMUSG00000096355_Ighv8-4 | -0,679160692 | 6,59248066 | 1,60774962 | 0,20480843 | 0,47447285 |
| ENSMUSG00000095519_Ighv1-66 | -0,500453107 | 8,57595717 | 1,57623443 | 0,20930393 | 0,4808801 |
| ENSMUSG00000095889_Ighv1-58 | -0,361396915 | 9,36345882 | 1,54469072 | 0,21392095 | 0,48435744 |
| ENSMUSG00000076534_Igkv12-89 | -0,410829361 | 8,95769102 | 1,54201517 | 0,2143181 | 0,48435744 |
| ENSMUSG00000094174_Ighv6-4 | 0,870134735 | 4,21543851 | 1,52390026 | 0,21703022 | 0,48435744 |
| ENSMUSG00000102765_Ighv1-62 | -0,586863251 | 7,43619533 | 1,50140182 | 0,2204558 | 0,48435744 |
| ENSMUSG00000105599_Gm9238 | -0,786277568 | 5,27895761 | 1,46407662 | 0,2262826 | 0,48435744 |
| ENSMUSG00000076940_Iglv2 | 0,535480051 | 13,8034696 | 1,46273654 | 0,22649521 | 0,48435744 |
| ENSMUSG00000094319_Igkv4-54 | 0,346279984 | 9,38795044 | 1,45309118 | 0,22803259 | 0,48435744 |
| ENSMUSG00000096108_Ighv11-2 | -0,720556782 | 8,95932044 | 1,4509443 | 0,22837649 | 0,48435744 |
| ENSMUSG00000076607_Igkj4 | 0,531183988 | 14,8029883 | 1,42745702 | 0,2321799 | 0,48435744 |
| ENSMUSG00000093906_Igkv9-129 | 0,331344172 | 10,0123279 | 1,41938817 | 0,23350413 | 0,48435744 |
| ENSMUSG00000104575_Igkv9-119 | 1,009895791 | 4,47058237 | 1,41429226 | 0,23434516 | 0,48435744 |
| ENSMUSG00000076532_Igkv4-91 | 0,43127226 | 8,69238204 | 1,40767817 | 0,23544221 | 0,48435744 |
| ENSMUSG00000076526_Igkv12-98 | 0,471417481 | 8,36153569 | 1,39581348 | 0,23742579 | 0,48435744 |
| ENSMUSG00000076505_Igkv1-131 | 0,413437289 | 8,77375358 | 1,39337613 | 0,23783578 | 0,48435744 |
| ENSMUSG00000105928_Gm9256 | 0,711054143 | 6,66474411 | 1,38232606 | 0,23970533 | 0,48435744 |
| ENSMUSG00000104422_Ighv1-14 | -0,92659312 | 3,8392428 | 1,38067141 | 0,23998681 | 0,48435744 |
| ENSMUSG00000076733_Ighv8-13 | -0,90725106 | 3,57763276 | 1,37803332 | 0,24043643 | 0,48435744 |
| ENSMUSG00000076535_Igkv1-88 | 0,334989463 | 9,52801973 | 1,3625797 | 0,2430909 | 0,48494172 |
| ENSMUSG00000093894_Ighv1-53 | -0,331863888 | 10,3608133 | 1,35609585 | 0,24421525 | 0,48494172 |
| ENSMUSG00000076583_Igkv8-24 | -0,30269033 | 9,87016226 | 1,33342668 | 0,24819652 | 0,48882087 |
| ENSMUSG00000094322_Ighv9-4 | 0,350445863 | 9,15944924 | 1,32506266 | 0,24968548 | 0,48882087 |
| ENSMUSG00000096490_Igkv10-94 | -0,364783491 | 9,55513238 | 1,2797716 | 0,25794151 | 0,49923956 |
| ENSMUSG00000106428_Gm42667 | 0,596237557 | 6,8861137 | 1,27213745 | 0,25936608 | 0,49923956 |
| ENSMUSG00000076533_Igkv4-90 | -0,677187307 | 6,17758504 | 1,26657052 | 0,26041104 | 0,49923956 |
| ENSMUSG00000096020_Ighv1-75 | -0,323557453 | 10,70344 | 1,25715362 | 0,26219056 | 0,49923956 |
| ENSMUSG00000076514_Igkv17-121 | 0,308582629 | 11,3131073 | 1,24171107 | 0,26514154 | 0,50142414 |
| ENSMUSG00000094356_Igkv8-28 | -0,310896373 | 9,53487803 | 1,21455404 | 0,27043208 | 0,50797377 |
| ENSMUSG00000076550_Igkv4-63 | 0,347530991 | 8,90454381 | 1,12611662 | 0,28860519 | 0,53513966 |
| ENSMUSG00000076522_Igkv16-104 | 0,271050448 | 10,2715059 | 1,12546645 | 0,28874442 | 0,53513966 |
| ENSMUSG00000106039_Iglc4 | 0,626748129 | 6,10958567 | 1,08547482 | 0,29747602 | 0,54428012 |
| ENSMUSG00000106372_Igkv14-126-1 | 0,640671539 | 5,10650691 | 1,08495374 | 0,29759201 | 0,54428012 |
| ENSMUSG00000076580_Igkv8-27 | 0,315112972 | 10,7556421 | 1,06093665 | 0,30300165 | 0,55055202 |

| | | | | | |
|---|---|---|---|---|---|
| ENSMUSG00000103290_Ighv1-23 | -0,552986541 | 6,51221135 | 0,97457853 | 0,32354096 | 0,57933155 |
| ENSMUSG00000106494_Igkv2-107 | -0,679545597 | 4,49058185 | 0,97354072 | 0,32379872 | 0,57933155 |
| ENSMUSG00000076652_Ighv7-3 | -0,255794517 | 10,0408208 | 0,96834802 | 0,32509252 | 0,57933155 |
| ENSMUSG00000094088_Ighv1-64 | -0,276951909 | 10,8868825 | 0,95644203 | 0,32808494 | 0,5809402 |
| ENSMUSG00000094124_Ighv1-74 | -0,256971563 | 10,1565174 | 0,9407982 | 0,33207265 | 0,58263025 |
| ENSMUSG00000105231_Iglj3 | 0,445552361 | 15,6411654 | 0,92919457 | 0,3350723 | 0,58263025 |
| ENSMUSG00000079543_Igkv13-85 | 0,263507337 | 9,54336062 | 0,92231842 | 0,33686697 | 0,58263025 |
| ENSMUSG00000094194_Ighv5-16 | 0,26056316 | 9,81106463 | 0,92019966 | 0,33742255 | 0,58263025 |
| ENSMUSG00000094797_Igkv6-15 | 0,271672034 | 10,5824756 | 0,89871219 | 0,34312725 | 0,58882331 |
| ENSMUSG00000076562_Igkv4-50 | -0,24157172 | 10,1554421 | 0,87299281 | 0,3501281 | 0,597151 |
| ENSMUSG00000095079_Igha | 0,622972057 | 7,81467079 | 0,84951311 | 0,35669011 | 0,60358819 |
| ENSMUSG00000096452_Ighv1-77 | -0,290538096 | 9,18020159 | 0,84403681 | 0,35824479 | 0,60358819 |
| ENSMUSG00000095592_Ighd5-7 | -0,649330507 | 5,00801704 | 0,81973547 | 0,36525752 | 0,61064709 |
| ENSMUSG00000095200_Ighv1-7 | -0,260085741 | 10,1568173 | 0,80668439 | 0,36910233 | 0,61064709 |
| ENSMUSG00000076564_Igkv12-46 | -0,244995 | 9,76198843 | 0,80287265 | 0,37023588 | 0,61064709 |
| ENSMUSG00000076594_Igkv6-13 | 0,272287622 | 9,23379843 | 0,79116689 | 0,37374752 | 0,61064709 |
| ENSMUSG00000094505_Ighv8-6 | 0,472182721 | 6,48449882 | 0,78362143 | 0,37603586 | 0,61064709 |
| ENSMUSG00000096805_Ighv9-1 | 0,259736521 | 10,0325341 | 0,77611813 | 0,37833099 | 0,61064709 |
| ENSMUSG00000076523_Igkv15-103 | 0,241075804 | 9,57426012 | 0,77426322 | 0,37890141 | 0,61064709 |
| ENSMUSG00000076577_Igkv8-30 | 0,251965001 | 9,6721933 | 0,77067926 | 0,380007 | 0,61064709 |
| ENSMUSG00000095007_Igkv12-41 | -0,233280903 | 9,65325154 | 0,73154946 | 0,39238141 | 0,62690823 |
| ENSMUSG00000076538_Igkv13-84 | 0,308077386 | 8,80405391 | 0,69754677 | 0,40360925 | 0,6348118 |
| ENSMUSG00000076525_Igkv1-99 | -0,481681387 | 6,00351146 | 0,69718639 | 0,40373073 | 0,6348118 |
| ENSMUSG00000105123_Ighv1-79 | -0,491407698 | 5,3351094 | 0,69326074 | 0,40505749 | 0,6348118 |
| ENSMUSG00000094006_Igkv4-59 | 0,21216088 | 10,3393884 | 0,68912478 | 0,40646223 | 0,6348118 |
| ENSMUSG00000076709_Ighv1-47 | -0,229032973 | 9,68623755 | 0,67518375 | 0,41125013 | 0,63707238 |
| ENSMUSG00000076608_Igkj5 | 0,377073531 | 15,486796 | 0,67160409 | 0,41249291 | 0,63707238 |
| ENSMUSG00000102524_Ighv1-2 | -0,254762567 | 9,13168914 | 0,64354316 | 0,42243067 | 0,64881617 |
| ENSMUSG00000118182_Gm50427 | -0,442921533 | 6,01716163 | 0,61649833 | 0,43235151 | 0,65967763 |
| ENSMUSG00000096638_Ighv2-9 | 0,234766291 | 9,19876365 | 0,60414049 | 0,43700259 | 0,65967763 |
| ENSMUSG00000076615_Ighg3 | -0,561090699 | 4,2901392 | 0,60395397 | 0,43707337 | 0,65967763 |
| ENSMUSG00000104975_Iglj2 | 0,753599928 | 14,6635427 | 0,59375423 | 0,44097104 | 0,65967763 |
| ENSMUSG00000095682_Igkv3-1 | 0,312557013 | 8,17792018 | 0,59272568 | 0,44136705 | 0,65967763 |
| ENSMUSG00000076518_Igkv2-112 | 0,263756771 | 8,85308283 | 0,57980687 | 0,44638803 | 0,66361429 |
| ENSMUSG00000094478_Igkv3-3 | 0,416937855 | 5,93614304 | 0,55996014 | 0,4542763 | 0,6665364 |
| ENSMUSG00000104679_Igkv4-60 | 0,31541137 | 7,98018412 | 0,55786806 | 0,45512051 | 0,6665364 |
| ENSMUSG00000105781_Igkv8-31 | -0,558734137 | 3,96971686 | 0,55489596 | 0,45632408 | 0,6665364 |
| ENSMUSG00000076591_Igkv8-16 | 0,20319541 | 10,7337683 | 0,55091499 | 0,45794407 | 0,6665364 |
| ENSMUSG00000105547_Iglc3 | 0,339635191 | 15,1081855 | 0,54347477 | 0,46099623 | 0,66748412 |
| ENSMUSG00000094315_Igkv4-78 | 0,248892032 | 8,89086979 | 0,5287429 | 0,46713585 | 0,67286925 |
| ENSMUSG00000076617_Ighm | 0,348183033 | 17,3514804 | 0,51663129 | 0,47228223 | 0,67677556 |
| ENSMUSG00000095565_Ighv2-9-1 | 0,184258808 | 10,0521685 | 0,48983886 | 0,48399919 | 0,6861848 |
| ENSMUSG00000106630_Igkv2-116 | 0,338122031 | 7,19670815 | 0,4792497 | 0,48876237 | 0,6861848 |
| ENSMUSG00000106256_Igkv12-47 | 0,405542736 | 5,95829757 | 0,47543499 | 0,49049739 | 0,6861848 |
| ENSMUSG00000104533_Igkv5-40-1 | 0,375379991 | 6,26698921 | 0,47494428 | 0,49072132 | 0,6861848 |
| ENSMUSG00000076540_Igkv4-80 | -0,281333597 | 8,40302214 | 0,47191206 | 0,49210884 | 0,6861848 |
| ENSMUSG00000105606_Igkv2-109 | 0,245014201 | 8,63256957 | 0,4685432 | 0,49365813 | 0,6861848 |
| ENSMUSG00000096715_Igkv3-4 | 0,198274599 | 9,23296256 | 0,45416514 | 0,5003636 | 0,68954181 |
| ENSMUSG00000076710_Ighv1-49 | -0,340366322 | 7,30480647 | 0,45274563 | 0,50103397 | 0,68954181 |
| ENSMUSG00000076547_Igkv4-70 | 0,246010923 | 8,4109251 | 0,41864495 | 0,51761396 | 0,70727918 |
| ENSMUSG00000096326_Ighv1-78 | -0,317656718 | 11,8539838 | 0,41585893 | 0,51901062 | 0,70727918 |
| ENSMUSG00000076609_Igkc | 0,291222181 | 16,7502319 | 0,37662463 | 0,53941523 | 0,73149968 |
| ENSMUSG00000094102_Ighv9-2 | -0,170329386 | 9,43557026 | 0,34528378 | 0,55679508 | 0,75140308 |

| | | | | | |
|---|---|---|---|---|---|
| ENSMUSG00000076512_Igkv9-123 | 0,271666192 | 7,48862618 | 0,33483406 | 0,56282639 | 0,75352941 |
| ENSMUSG00000076556_Igkv4-57 | 0,154728655 | 10,5094676 | 0,33318133 | 0,56379179 | 0,75352941 |
| ENSMUSG00000105432_Gm43218 | 0,148564981 | 10,3498374 | 0,30389826 | 0,58144882 | 0,76992341 |
| ENSMUSG00000096833_Igkv4-55 | 0,14409403 | 10,0273607 | 0,30214115 | 0,58254321 | 0,76992341 |
| ENSMUSG00000076549_Igkv4-68 | -0,141048878 | 10,5684415 | 0,29922876 | 0,58436633 | 0,76992341 |
| ENSMUSG00000076555_Igkv4-57-1 | -0,262436959 | 7,71067725 | 0,2867031 | 0,59234105 | 0,77674911 |
| ENSMUSG00000076552_Igkv4-61 | -0,186117332 | 8,66037059 | 0,2755353 | 0,59964258 | 0,7826321 |
| ENSMUSG00000094509_Ighv14-1 | -0,189970368 | 8,69376466 | 0,26431842 | 0,60716874 | 0,78875192 |
| ENSMUSG00000105605_Ighv1-86 | -0,361815371 | 4,23368518 | 0,2555706 | 0,61317967 | 0,79285558 |
| ENSMUSG00000076688_Ighv15-2 | -0,149383467 | 9,48570109 | 0,24924357 | 0,61760821 | 0,79488464 |
| ENSMUSG00000076563_Igkv5-48 | -0,139570571 | 9,40559736 | 0,24004421 | 0,62417419 | 0,79963329 |
| ENSMUSG00000106239_Gm9260 | 0,287001875 | 5,16385006 | 0,21634036 | 0,64184264 | 0,81408531 |
| ENSMUSG00000102364_Ighv8-5 | -0,161869405 | 8,99789554 | 0,21510133 | 0,64279808 | 0,81408531 |
| ENSMUSG00000076655_Ighv4-1 | -0,121571431 | 10,5384202 | 0,20891783 | 0,64761693 | 0,81408531 |
| ENSMUSG00000105363_Igkv11-114 | 0,240278976 | 6,34800144 | 0,20184216 | 0,653238 | 0,81408531 |
| ENSMUSG00000098814_Igkv19-93 | 0,124634903 | 11,778665 | 0,19659474 | 0,65748363 | 0,81408531 |
| ENSMUSG00000096670_Ighv2-6 | 0,155854184 | 8,76412662 | 0,19629158 | 0,65773097 | 0,81408531 |
| ENSMUSG00000076677_Ighv6-3 | -0,120845008 | 11,0785089 | 0,19193504 | 0,66131095 | 0,81408531 |
| ENSMUSG00000076618_Ighj4 | -0,186959068 | 14,1884986 | 0,1901091 | 0,66282583 | 0,81408531 |
| ENSMUSG00000094335_Igkv1-117 | 0,114967801 | 10,8050238 | 0,18932363 | 0,66348015 | 0,81408531 |
| ENSMUSG00000094491_Igkv1-133 | 0,139406554 | 9,00614621 | 0,18474283 | 0,66732862 | 0,81408531 |
| ENSMUSG00000093876_Ighd5-3 | 0,276261958 | 6,71784344 | 0,18210758 | 0,66956824 | 0,81408531 |
| ENSMUSG00000076665_Ighv7-1 | -0,120348616 | 9,50395325 | 0,1809064 | 0,67059545 | 0,81408531 |
| ENSMUSG00000106387_Igkv4-73 | -0,137780003 | 9,07078641 | 0,17708438 | 0,67389094 | 0,81452905 |
| ENSMUSG00000076605_Igkj2 | 0,176885429 | 15,0225811 | 0,1568367 | 0,69208575 | 0,82639415 |
| ENSMUSG00000076530_Igkv11-106 | -0,18736427 | 7,45791349 | 0,15563006 | 0,6932121 | 0,82639415 |
| ENSMUSG00000096632_Igkv9-124 | -0,108582598 | 10,6291738 | 0,15468461 | 0,69409819 | 0,82639415 |
| ENSMUSG00000095497_Igkv1-122 | -0,12694671 | 9,0595918 | 0,14999015 | 0,69854477 | 0,82639415 |
| ENSMUSG00000076587_Igkv6-20 | -0,162908938 | 8,54752757 | 0,14996313 | 0,6985706 | 0,82639415 |
| ENSMUSG00000096580_Igkv1-132 | 0,186139319 | 7,36224049 | 0,13940096 | 0,70887726 | 0,833794 |
| ENSMUSG00000095753_Igkv4-53 | -0,095777546 | 10,2749611 | 0,13745493 | 0,71082438 | 0,833794 |
| ENSMUSG00000076541_Igkv4-79 | 0,121632105 | 8,9398792 | 0,12051305 | 0,72847871 | 0,85091211 |
| ENSMUSG00000106405_Iglj3p | 0,191227185 | 7,8650562 | 0,11487126 | 0,734665 | 0,85454758 |
| ENSMUSG00000076573_Igkv1-35 | 0,209845865 | 4,76780112 | 0,11113314 | 0,73885774 | 0,85584355 |
| ENSMUSG00000076576_Igkv6-32 | -0,080550462 | 10,1078751 | 0,10026057 | 0,75151717 | 0,86499685 |
| ENSMUSG00000105499_Igkv3-11 | 0,180479765 | 5,86457245 | 0,09896565 | 0,75307444 | 0,86499685 |
| ENSMUSG00000076604_Igkj1 | 0,137928839 | 15,023093 | 0,09648083 | 0,75609437 | 0,86499685 |
| ENSMUSG00000102381_Ighv8-7 | 0,18465745 | 5,86355052 | 0,08499911 | 0,77063371 | 0,8780171 |
| ENSMUSG00000076548_Igkv4-69 | -0,093399629 | 9,04179871 | 0,08116068 | 0,77573061 | 0,88021678 |
| ENSMUSG00000096577_Ighv1-71 | 0,092346246 | 8,9955719 | 0,07753822 | 0,78066167 | 0,88037455 |
| ENSMUSG00000095633_Igkv4-58 | -0,094153211 | 9,70899731 | 0,076424 | 0,78220329 | 0,88037455 |
| ENSMUSG00000076674_Ighv3-8 | 0,073361619 | 10,1637542 | 0,07076587 | 0,79022492 | 0,88581665 |
| ENSMUSG00000095351_Igkv3-2 | -0,065360601 | 10,2199948 | 0,06734516 | 0,79524219 | 0,88669299 |
| ENSMUSG00000096767_Ighv1-62-3 | -0,086999961 | 8,90785895 | 0,06591124 | 0,79738578 | 0,88669299 |
| ENSMUSG00000076619_Ighj3 | -0,101747365 | 14,1264234 | 0,05606735 | 0,81282324 | 0,89282621 |
| ENSMUSG00000076571_Igkv5-37 | -0,144791447 | 5,6630405 | 0,05508074 | 0,81444712 | 0,89282621 |
| ENSMUSG00000095642_Ighv14-3 | -0,059290379 | 10,1187981 | 0,05482572 | 0,81486936 | 0,89282621 |
| ENSMUSG00000104742_Igkv9-128 | -0,1921414 | 4,40318622 | 0,05429723 | 0,81574769 | 0,89282621 |
| ENSMUSG00000076572_Igkv18-36 | 0,109422805 | 6,83412762 | 0,04664122 | 0,82901443 | 0,90073515 |
| ENSMUSG00000091087_Ighv8-14 | -0,185163176 | 2,99762025 | 0,046398 | 0,82945395 | 0,90073515 |
| ENSMUSG00000094872_Igkv9-120 | 0,055844358 | 11,1235654 | 0,04353472 | 0,83472167 | 0,9029285 |
| ENSMUSG00000076614_Ighg1 | -0,138190691 | 3,9784508 | 0,03993818 | 0,8416015 | 0,90497231 |
| ENSMUSG00000076536_Igkv4-86 | -0,053758181 | 9,86559969 | 0,03916523 | 0,84312168 | 0,90497231 |

| | | | | | |
|---|---|---|---|---|---|
| ENSMUSG00000096422_Igkv12-44 | 0,048912709 | 10,0671197 | 0,03659978 | 0,84828218 | 0,90700941 |
| ENSMUSG00000094902_Igkv10-95 | -0,049115534 | 10,055638 | 0,03466202 | 0,85230565 | 0,90781982 |
| ENSMUSG00000093861_Igkv1-110 | 0,043759949 | 10,0831495 | 0,02841619 | 0,86613407 | 0,91902776 |
| ENSMUSG00000106667_Igkv3-6 | -0,105053915 | 4,13129792 | 0,02306254 | 0,87929456 | 0,92760834 |
| ENSMUSG00000076508_Igkv17-127 | 0,03972502 | 11,4242513 | 0,02131987 | 0,88391092 | 0,92760834 |
| ENSMUSG00000076586_Igkv8-21 | 0,04034298 | 9,71480056 | 0,01913409 | 0,88998276 | 0,92760834 |
| ENSMUSG00000102901_Ighv5-1 | 0,071624359 | 6,66648494 | 0,01799088 | 0,89329972 | 0,92760834 |
| ENSMUSG00000102535_Ighv1-41 | -0,0722863 | 6,75060593 | 0,01768225 | 0,89421346 | 0,92760834 |
| ENSMUSG00000076939_Iglv3 | 0,055580193 | 13,3533477 | 0,01767294 | 0,89424114 | 0,92760834 |
| ENSMUSG00000103939_Ighv3-4 | 0,077553904 | 6,27476052 | 0,01499839 | 0,90252844 | 0,93272456 |
| ENSMUSG00000096459_Ighv9-3 | -0,029310753 | 12,2720502 | 0,00813855 | 0,92811722 | 0,95561699 |
| ENSMUSG00000095210_Ighv5-9-1 | 0,022084337 | 9,94400644 | 0,00651178 | 0,93568404 | 0,959853 |
| ENSMUSG00000076569_Igkv5-39 | -0,013586154 | 11,1590319 | 0,00255699 | 0,95967083 | 0,98084003 |
| ENSMUSG00000076545_Igkv4-72 | -0,01190012 | 9,09632534 | 0,00147271 | 0,96938801 | 0,98401258 |
| ENSMUSG00000106016_Igkv4-56 | 0,017566271 | 7,48033277 | 0,00142818 | 0,96985412 | 0,98401258 |
| ENSMUSG00000095583_Ighv14-2 | 0,006201916 | 10,4775504 | 0,00052578 | 0,98170627 | 0,98975896 |
| ENSMUSG00000076539_Igkv4-81 | 0,011010796 | 7,27230913 | 0,00047355 | 0,9826384 | 0,98975896 |
| ENSMUSG00000094930_Igkv6-25 | 0,003325808 | 9,72610993 | 0,00015357 | 0,99011252 | 0,99366989 |
| ENSMUSG00000076646_Ighv2-6-8 | -0,003638691 | 7,72934721 | 6,29E-05 | 0,99366989 | 0,99366989 |
