## Supplementary Table 4 for "The chromatin reader Dido3 is a regulator of the gene network that controls B cell differentiation"

**Supplementary Table 4. ChIPseeker annotations of the H3K27me3 ChIP-seq data.**

| ID | #Peaks | Condition | Percentage of H3K27me3 peaks | | | | | | | | |
|---|---|---|---|---|---|---|---|---|---|---|---|
| | | | Promoter | 5'-UTR | 3'-UTR | 1st Exon | Other Exon | 1st Intron | Other Intron | Down-stream | Distal Intergenic |
| HR45_S1 | 898 | WT (replicate 1) | 1.89 | 0.22 | 1.22 | 2.56 | 4.12 | 10.8 | 21.49 | 0.11 | 57.57 |
| HR60_S4 | 894 | WT (replicate 2) | 2.01 | 0.34 | 1.12 | 2.46 | 3.47 | 10.51 | 21.92 | - | 58.16 |
| HR61_S5 | 1212 | WT (replicate 3) | 1.89 | 0.25 | 1.24 | 2.72 | 3.96 | 9.65 | 22.52 | - | 57.76 |
| HR56_S2 | 827 | *Dido1*ΔE16 (replicate 1) | 1.57 | 0.36 | 1.33 | 2.66 | 4.35 | 10.16 | 23.58 | - | 55.98 |
| HR58_S3 | 1356 | *Dido1*ΔE16 (replicate 2) | 1.99 | 0.29 | 1.4 | 2.36 | 3.69 | 10.84 | 21.75 | - | 57.67 |
| HR62_S6 | 973 | *Dido1*ΔE16 (replicate 3) | 1.85 | 0.31 | 1.34 | 2.36 | 4.01 | 9.97 | 23.23 | - | 56.94 |

Numbers in columns (promoter, 5'-UTR, 3'-UTR, 1st exon, other exon, 1st intron, other intron, downstream, distal intergenic) indicate the percentage of H3K27me3 peaks overlapping each genomic region. ID: identifier of biological replicates in the GEO dataset GSE272156. #Peaks: total number of H3K27me3 peaks detected with the callpeak function in MACS version 3.0.0b3. Condition: source of the LSK cells and their biological replicates in parenthesis.
