## Supplementary Table 5 for "The chromatin reader Dido3 is a regulator of the gene network that controls B cell differentiation"

**Supplementary Table 5. ChIPseeker annotations of the H3K27me3-enriched genomic regions.**

| **H3K27me3 enrichment** | **Percentage of H3K27me3 peaks** | | | | | | | | | **Gene symbols[a]** |
|---|---|---|---|---|---|---|---|---|---|---|
| | **Promoter** | **5'-UTR** | **3'-UTR** | **1st Exon** | **Other Exon** | **1st Intron** | **Other Intron** | **Down-stream** | **Distal Intergenic** | |
| WT | 1.72 | - | 3.45 | - | - | 8.62 | 15.52 | - | 70.69 | Cd180, Cpeb4, Hmgxb4, Snrpe |
| *Dido1*ΔE16 | - | - | - | - | - | 6.82 | 20.45 | - | 72.73 | Alx1,Atg7,Camk1d,Dclk2,Eya1, Marchf1,Snrpn,Spin4,Tet2,Zxda |
| Common regions | 1.38 | 0.55 | 1.1 | 2.75 | 5.37 | 10.47 | 21.76 | - | 56.61 | *Blk*,Cdk18,Cr2,Erdr1x,Ezh2,Fgfr2,*Gpm6a*, *Gramd2b(Gramd3)*,Hdac9,Hira,Jarid2,*Klra1*, Pou5f1-rs8,Snrpe,Srsf4,Trim30a,Zfp607a, Zfp746,Zfp825,Zfp945,Zfp958,Zfp980, Zfp982 |
| **Control dataset annotations** | | | | | | | | | | |
| Control IgG | 1.36 | 0.34 | 1.9 | 2.07 | 3.11 | 12.16 | 28.11 | 0.07 | 50.87 | |
| Control H3K4me3 | 32.21 | 0.99 | 2.21 | 3.01 | 4.08 | 10.18 | 17.06 | 0.09 | 30.16 | |

Numbers in columns (promoter, 5'-UTR, 3'-UTR, 1st exon, other exon, 1st intron, other intron, downstream, distal intergenic) indicate the percentage of H3K27me3 peaks overlapping each genomic region. [a]The complete list of genes within 1.0 kb upstream to 200 bp downstream of transcriptional start sites is available in Supplementary File section. Text in italics and underline indicates genes that were differentially expressed in the RNA-seq of the pre-B cells.
