## Supplementary Table 6 for "The chromatin reader Dido3 is a regulator of the gene network that controls B cell differentiation"

**Supplementary Table 6. Summary of H3K27me3 ChIP-seq data in LSK cells.**

| Study | LSK cells | GEO dataset | #Peaks | H3K27me3 Ab | Bioinformatic pipeline[a] |
|---|---|---|---|---|---|
| This study | WT (replicate 3) | GSE272156 | 1212 | Abcam (ab195477) | MACS3 callpeak -B --extsize 147 -g mm -q 0.05 –broad --broad-cutoff 0.1 --nomodel --extsize 147 MACS3 bdgdiff -g 73 -l 147 (b) |
| | *Dido1*dE16 (replicate 2) | GSE272156 | 1356 | | |
| Yang *et al.* Blood (2016) DOI:10.1182/blood-2015-11-679431 | Ezh2-WT | GSM2091489 | 25541 | Millipore (07-449) | Strand NGS Version 2.0 (Strand Genomics) with MACS method |
| | Ezh2-KO | GSM2091491 | 29626 | | |
| Hasemann *et al.* PLoS Genet (2014) DOI:10.1371/journal.pgen.1004079 | Cebpa-WT | GSM1054811 | 239 | Cell Signaling (C36B11) | MACS2 callpeak -g mm -p 0.001 --to-large |
| | Cebpa-KO | GSM1054814 | 597 | | |

[a]For more details see https://github.com/macs3-project/MACS/wiki/Call-differential-binding-events
