## Supplementary Table 7 for "The chromatin reader Dido3 is a regulator of the gene network that controls B cell differentiation"

**Supplementary Table 7. ChIPseeker annotations of H3K27me3 ChIP-seq data in LSK cells.**

| Dataset | Promoter | 5'-UTR | 3'-UTR | 1st Exon | Other Exon | 1st Intron | Other Intron | Down-stream | Distal Intergenic |
|---|---|---|---|---|---|---|---|---|---|
| WT (replicate 3) | 1.89 | 0.25 | 1.24 | 2.72 | 3.96 | 9.65 | 22.52 | - | 57.76 |
| dE16 (replicate 2) | 1.99 | 0.29 | 1.4 | 2.36 | 3.69 | 10.84 | 21.75 | - | 57.67 |
| Ezh2-WT (GSM2091489) | 0.99 | 0.05 | 0.48 | 0.52 | 0.24 | 12.15 | 25.87 | 0.1 | 59.59 |
| Ezh2-KO (GSM2091491) | 1.02 | 0.05 | 0.46 | 0.43 | 0.25 | 11.99 | 25.79 | 0.08 | 59.93 |
| Cebpa-WT (GSM1054811) | 1.25 | - | 0.84 | 8.37 | 3.35 | 12.55 | 24.27 | - | 49.37 |
| Cebpa-KO (GSM1054814) | 1.17 | 0.17 | 1.17 | 4.86 | 2.68 | 17.59 | 26.29 | - | 46.06 |

Numbers in columns (promoter, 5'-UTR, 3'-UTR, 1st exon, other exon, 1st intron, other intron, downstream, distal intergenic) indicate the percentage of H3K27me3 peaks overlapping each genomic region.
Dataset: identifier of H3K27me3 ChIP-seq data and their GEO accession number in parenthesis.
