## Supplementary Table 8 for "The chromatin reader Dido3 is a regulator of the gene network that controls B cell differentiation"

**Supplementary Table 8. H3K27me3 peaks overlap enrichment analysis.**

| qSample[a] | GEO acc. | tSample[b] | qLen[c] | tLen[d] | N_OL[e] | N_OL(%qLen)[f] | p-value[g] | p.adjust[h] |
|---|---|---|---|---|---|---|---|---|
| WT (replicate 3) | GSM2091489 | Ezh2-WT | 1212 | 25541 | 107 | 8.8% | $4.9 \times 10^{-4}$ | $5.9 \times 10^{-4}$ |
| | GSM2091491 | Ezh2-KO | 1212 | 29626 | 106 | 8.7% | $1.02 \times 10^{-2}$ | $1.02 \times 10^{-2}$ |
| | GSM1054811 | Cebpa-WT | 1212 | 239 | 117 | 9.6% | $9.9 \times 10^{-5}$ | $1.5 \times 10^{-4}$ |
| | GSM1054814 | Cebpa-KO | 1212 | 597 | 146 | **12%** | $9.9 \times 10^{-5}$ | **$1.5 \times 10^{-4}$** |
| dE16 (replicate 2) | GSM2091489 | Ezh2-WT | 1356 | 25541 | 83 | 6.1% | $1.3 \times 10^{-2}$ | $1.3 \times 10^{-2}$ |
| | GSM2091491 | Ezh2-KO | 1356 | 29626 | 86 | 6.3% | $1.3 \times 10^{-2}$ | $1.3 \times 10^{-2}$ |
| | GSM1054811 | Cebpa-WT | 1356 | 239 | 117 | 8.6% | $9.9 \times 10^{-5}$ | $1.5 \times 10^{-4}$ |
| | GSM1054814 | Cebpa-KO | 1356 | 597 | 144 | **10.6%** | $9.9 \times 10^{-5}$ | **$1.5 \times 10^{-4}$** |

[a]Query ChIP-seq sample, [b]Target ChIP-seq sample, [c]Number of query peaks, [d]Number of target peaks, [e]Number of overlapped peaks between query and target, [f]Percentage of overlapped peaks, [g]calculated p-value by ChIPseeker, [h]p-value correction (FDR) according to the Benjamini and Hochberg method. GEO database accession of tSample (GEO acc.). The values were obtained using the ChIPseeker command enrichPeakOverlap(queryPeak=file1, targetPeak=file-list, TxDb=TxDb.Mmusculus.UCSC.mm10.knownGene, pAdjustMethod="BH", nShuffle=10000, chainFile=NULL, verbose=FALSE) and a number of randomly permutations in the genomic locations of 10000.
