## Supplementary Table 9 for "The chromatin reader Dido3 is a regulator of the gene network that controls B cell differentiation"

**Supplementary Table 9. ChIP-seq H3K27me3 peaks and ATAC-seq chromatin-accessible regions overlap enrichment analysis.**

| **qSample**[a] | **tSource** | **tSample**[b] | **qLen**[c] | **tLen**[d] | **N_OL**[e] | **N_OL(%qLen)**[f] | **p-value**[g] | **p.adjust**[h] | Gene symbols |
|---|---|---|---|---|---|---|---|---|---|
| WT(replicate 3) | ATAC-seq | WT(open) | 1212 | 26327 | 272 | 22.4% | $1.8 \times 10^{-2}$ | $2.3 \times 10^{-2}$ | (i) |
| | | WT(open)/MUT(open) | 1212 | 9197 | 219 | 18.1% | $2.2 \times 10^{-3}$ | $3.7 \times 10^{-3}$ | (i) |
| | | MUT(open) | 1212 | 9795 | **333** | **27.5%** | $3.9 \times 10^{-4}$ | **$9.9 \times 10^{-4}$** | (i) |
| | | WT(open)/MUT(close) | 1212 | 11007 | 7 | 0.6% | 0.6 | 0.6 | - |
| | | WT(close)/MUT(open) | 1212 | 81 | **11** | **0.9%** | $9.9 \times 10^{-5}$ | **$4.9 \times 10^{-4}$** | 4930448K20Rik,Adam29,B230307C23Rik, Bcas1,G530011O06Rik,Gm12018,Gm17019, Mid1,Pdia3,Rimbp2,Speer4d |
| dE16(replicate 2) | | WT(open) | 1356 | 26327 | 276 | 20.3% | $1.9 \times 10^{-2}$ | $2.4 \times 10^{-2}$ | (i) |
| | | WT(open)/MUT(open) | 1356 | 9197 | 219 | 16.2% | $2.9 \times 10^{-3}$ | $4.9 \times 10^{-3}$ | (i) |
| | | MUT(open) | 1356 | 9795 | **346** | **25.5%** | $2.9 \times 10^{-4}$ | **$7.5 \times 10^{-4}$** | (i) |
| | | WT(open)/MUT(close) | 1356 | 11007 | 8 | 0.6% | 0.6 | 0.6 | - |
| | | WT(close)/MUT(open) | 1356 | 81 | **14** | **1%** | $9.9 \times 10^{-5}$ | **$4.9 \times 10^{-4}$** | 4930448K20Rik,Adam29,1700064M15Rik, B230307C23Rik,Bcas1,G530011O06Rik, Gm12018,Gm17019,Gm21190,Mid1,Pdia3, Rimbp2,Speer4d |

[a]Query ChIP-seq sample, [b]Target ChIP-seq sample, [c]Number of query peaks, [d]Number of target peaks, [e]Number of overlapped peaks between query and target, [f]Percentage of overlapped peaks, [g]calculated p-value by ChIPseeker,[h]p-value correction (FDR) according to the Benjamini and Hochberg method. [i]The complete list of genes within 1.0 kb upstream to 200 bp downstream of transcriptional start sites is available in Supplementary File section. The values were obtained using the ChIPseeker command enrichPeakOverlap(queryPeak=file1, targetPeak=file-list, TxDb=TxDb.Mmusculus.UCSC.mm10.knownGene, pAdjustMethod="BH", nShuffle=10000, chainFile=NULL, verbose=FALSE) and a number of randomly permutations in the genomic locations of 10000.
