## Supplementary Table 10 for "The chromatin reader Dido3 is a regulator of the gene network that controls B cell differentiation"

**Supplementary Table 10. ChIP-seq H3K4me3 peaks and ATAC-seq chromatin-accessible regions overlap enrichment analysis.**

| **qSample[a]** | **tSource** | **tSample[b]** | **qLen[c]** | **tLen[d]** | **N_OL[e]** | **N_OL(%qLen)[f]** | **p-value[g]** | **p.adjust[h]** |
|---|---|---|---|---|---|---|---|---|
| H3K4me3 | ATAC-seq | WT(open) | 47243 | 26327 | 17663 | **37.4%** | $9.9 \times 10^{-5}$ | $9.9 \times 10^{-5}$ |
| H3K4me3 | | WT(open)/MUT(open) | 47243 | 9197 | 8301 | 17.6% | $9.9 \times 10^{-5}$ | $9.9 \times 10^{-5}$ |
| H3K4me3 | | MUT(open) | 47243 | 9795 | 8995 | **19%** | $9.9 \times 10^{-5}$ | $9.9 \times 10^{-5}$ |
| H3K4me3 | | WT(open)/MUT(close) | 47243 | 11007 | 4990 | 10.6% | $9.9 \times 10^{-5}$ | $9.9 \times 10^{-5}$ |
| H3K4me3 | | WT(close)/MUT(open) | 47243 | 81 | 17 | 0.04% | $9.9 \times 10^{-5}$ | $9.9 \times 10^{-5}$ |
| H3K4me3 | ChIP-seq (H3K27me3) | WT (enrichment) | 47243 | 58 | 41 | 0.09% | $9.9 \times 10^{-5}$ | $9.9 \times 10^{-5}$ |
| H3K4me3 | | dE16 (enrichment) | 47243 | 44 | 8 | 0.02% | $9.9 \times 10^{-5}$ | $9.9 \times 10^{-5}$ |
| H3K4me3 | | Common | 47243 | 726 | 645 | 1.4% | $9.9 \times 10^{-5}$ | $9.9 \times 10^{-5}$ |
| H3K4me3 | FANTOM5 | F5.mm10.enhancers | 47243 | 49797 | 7771 | 16.4% | $9.9 \times 10^{-5}$ | $9.9 \times 10^{-5}$ |

[a]Query ChIP-seq sample, [b]Target ChIP-seq sample, [c]Number of query peaks, [d]Number of target peaks, [e]Number of overlapped peaks between query and target, [f]Percentage of overlapped peaks, [g]calculated p-value by ChIPseeker, [h]p-value correction (FDR) according to the Benjamini and Hochberg method. The values were obtained using the ChIPseeker command enrichPeakOverlap(queryPeak=file1, targetPeak=file-list, TxDb=TxDb.Mmusculus.UCSC.mm10.knownGene, pAdjustMethod="BH", nShuffle=10000, chainFile=NULL, verbose=FALSE) and a number of randomly permutations in the genomic locations of 10000.
