## Supplementary Table 11 for "The chromatin reader Dido3 is a regulator of the gene network that controls B cell differentiation"

**Supplementary Table 11. List of bivalent genes with chromatin accessibility alterations and H3K4me3 marks in the promoter that are differently expressed.**

| **Gene** | **ChIP-seq** | | **ATAC-seq (WT/MUT)** | | | **RNA-seq** | | **Description** |
|---|---|---|---|---|---|---|---|---|
| | H3K4me3 | H3K27me3 | (cl/op) | (op/cl) | (op/op) | DOWN | UP | |
| Ctsh | 1 | 0 | 0 | 0 | 1 | 1 | 0 | cathepsin H |
| Atg16l2 | 1 | 0 | 0 | 0 | 1 | 1 | 0 | autophagy related 16 like 2 |
| Casz1 | 1 | 0 | 0 | 0 | 0 | 1 | 0 | castor zinc finger 1 |
| Rassf1 | 1 | 0 | 0 | 0 | 1 | 1 | 0 | Ras association (RalGDS/AF-6) domain family member 1 |
| **Sorcs2** | 1 | 0 | 0 | 1 | 0 | 1 | 0 | sortilin-related VPS10 domain containing receptor 2 |
| **Reln** | 1 | 0 | 0 | 1 | 0 | 1 | 0 | Reelin |
| Mef2d | 1 | 0 | 0 | 0 | 1 | 1 | 0 | myocyte enhancer factor 2D |
| H2-Ab1 | 1 | 0 | 1 | 0 | 0 | 1 | 0 | histocompatibility 2, class II antigen A, beta 1 |
| Runx3 | 1 | 0 | 0 | 0 | 1 | 0 | 1 | runt related transcription factor 3 |
| Macf1 | 1 | 0 | 0 | 0 | 1 | 0 | 1 | microtubule-actin crosslinking factor 1 |
| Pdzd2 | 1 | 0 | 0 | 0 | 1 | 0 | 1 | PDZ domain containing 2 |
| Nfkb2 | 1 | 0 | 0 | 0 | 1 | 0 | 1 | nuclear factor of kappa light polypeptide gene enhancer in B cells 2, p49/p100 |
| Phactr2 | 1 | 0 | 0 | 1 | 0 | 0 | 1 | phosphatase and actin regulator 2 |
| Arhgap31 | 1 | 0 | 0 | 1 | 0 | 0 | 1 | Rho GTPase activating protein 31 |
| Sox18 | 1 | 0 | 0 | 1 | 0 | 0 | 1 | SRY (sex determining region Y)-box 18 |
| Adm | 1 | 0 | 0 | 1 | 0 | 0 | 1 | adrenomedullin |
| Cnnm2 | 1 | 0 | 0 | 0 | 1 | 0 | 1 | cyclin M2 |
| Lgr5 | 1 | 0 | 0 | 1 | 0 | 0 | 1 | leucine rich repeat containing G protein coupled receptor 5 |
| Nefh | 1 | 0 | 0 | 1 | 0 | 0 | 1 | neurofilament, heavy polypeptide |
| Cplx1 | 1 | 0 | 0 | 1 | 0 | 0 | 1 | complexin 1 |
| Plxna2 | 1 | 0 | 0 | 1 | 0 | 0 | 1 | plexin A2 |
| Heyl | 1 | 0 | 0 | 1 | 0 | 0 | 1 | hairy/enhancer-of-split related with YRPW motif-like |
| Tnfrsf21 | 1 | 0 | 0 | 0 | 1 | 0 | 1 | tumor necrosis factor receptor superfamily, member 21 |
| Nhsl2 | 1 | 0 | 0 | 1 | 0 | 0 | 1 | NHS-like 2 |
| Thrb | 1 | 0 | 0 | 0 | 1 | 0 | 1 | thyroid hormone receptor beta |
| Apba1 | 1 | 0 | 0 | 0 | 1 | 0 | 1 | amyloid beta precursor protein binding family A member 1 |
| Gpr157 | 1 | 0 | 0 | 0 | 1 | 0 | 1 | G protein-coupled receptor 157 |
| Lrrc8b | 1 | 0 | 0 | 0 | 1 | 0 | 1 | leucine rich repeat containing 8 family, member B |
| Arid5a | 1 | 0 | 0 | 0 | 1 | 0 | 1 | AT-rich interaction domain 5A |
| Epb41l4b | 1 | 0 | 0 | 1 | 0 | 0 | 1 | erythrocyte membrane protein band 4.1 like 4b |
| Begain | 1 | 0 | 0 | 1 | 0 | 0 | 1 | brain-enriched guanylate kinase-associated |
| Mypop | 1 | 0 | 0 | 0 | 1 | 0 | 1 | Myb-related transcription factor, partner of profilin |
| Ctif | 1 | 0 | 0 | 1 | 0 | 0 | 1 | CBP80/20-dependent translation initiation factor |

The chromatin regions that are either close (cl) or open (op) are indicated using ATAC-seq data. One denotes the existence of a differentially expressed gene, a chromatin accessible region, or a histone mark, whereas cero denotes their absence.
